# Characterization of three resistance-breaking isolates of sugarcane mosaic virus from Rwanda and implications for maize lethal necrosis

**DOI:** 10.1101/2024.07.23.604352

**Authors:** Jennifer R. Wilson, Kristin J. Willie, Lucy R. Stewart, Margaret G. Redinbaugh, Erik W. Ohlson

## Abstract

Maize lethal necrosis (MLN) is a devastating disease of maize caused by synergy between two viruses: maize chlorotic mottle virus (MCMV) and a potyvirus, most often sugarcane mosaic virus (SCMV). Throughout the 2010s, severe MLN outbreaks occurred in sub-Saharan East Africa including Kenya, Rwanda, and Ethiopia. In this study, we assessed the virulence of SCMV isolates collected from Rwanda by screening a panel of maize near isogenic lines containing different combinations of major potyvirus resistance loci. We discovered that the three Rwandan SCMV isolates tested could overcome all three potyvirus resistance loci even when used in combination, including one isolate that could asymptomatically infect all resistant lines tested. To understand how SCMV virulence may contribute to MLN, each of the three isolates were co-inoculated with MCMV on a panel of SCMV and MCMV resistant maize lines. No significant differences in MLN severity were observed for the Rwandan isolates compared to the reference SCMV isolates, indicating that increased virulence in SCMV single infection does not necessarily correlate with increased MLN severity in co-infection with MCMV. For all SCMV isolates tested, at least two potyvirus resistance loci were needed to reduce MLN severity, and maize lines with a combination of SCMV and MCMV resistance were most effective. Surprisingly, in some cases co-infection with MCMV facilitated SCMV infection of potyvirus resistant lines that SCMV could not infect alone. These results underscore the challenges of developing durable MLN resistance and highlight the importance of incorporating strong, multigenic potyvirus resistance into MLN resistance breeding programs.

## Introduction

Maize lethal necrosis (MLN) disease is a classic example of viral synergy. It is caused by the co-infection of maize chlorotic mottle virus (MCMV) and one of several viruses in the *Potyviridae* family, including sugarcane mosaic virus (SCMV), maize dwarf mosaic virus (MDMV), or wheat streak mosaic virus (WSMV) (Niblett & Claflin, 1978, Wangai *et al*., 2012, Stewart *et al*., 2017, Scheets, 1998). While single infection by any of these potyviruses only causes mosaic and stunting, co-infection with MCMV leads to more severe mosaic, severe stunting, chlorosis, necrosis, and eventual plant death (reviewed in (Ohlson & Wilson, 2022). MLN was first described in the United States in 1977 (Niblett & Claflin, 1978) but has since spread to several other continents, including sub-Saharan East Africa, Southeast Asia, and South America (Mahuku *et al*., 2015b, Adams *et al*., 2014, Deng *et al*., 2014, Wangai et al., 2012, Lukanda *et al*., 2014, Quito-Avila *et al*., 2016, Xie *et al*., 2011). The disease has been particularly devastating in East Africa, where it has caused a combined $300 million in damages each year throughout the 2010s in Ethiopia, Kenya, Rwanda, and Tanzania (Pratt *et al*., 2017).

Management of MLN relies heavily on host plant resistance. Significant efforts have been made to identify and develop MCMV resistant maize lines. However, resistance to MCMV remains poorly described and quantitative in nature. Quantitative trait loci associated with MCMV resistance have been identified on every maize chromosome (Jones *et al*., 2018, Ohlson *et al*., 2022, Sitonik *et al*., 2019) but, to date, not a single causal gene has been reported, though fine mapping projects are progressing (Jones et al., 2018, Ohlson et al., 2022, Sitonik et al., 2019). Moreover, MCMV is readily detected in most MCMV “resistant” lines via enzyme linked immunosorbent assay (ELISA)(Jones et al., 2018) and reverse transcriptase polymerase chain reaction (RT-PCR)(Gentzel *et al*., 2024) even though the plants show no symptoms, indicating that the resistance is incomplete, which is often termed tolerance.

Potyvirus resistance, on the other hand, is much better understood. Three major potyvirus resistance loci have been identified on chromosomes 6 (*Scmv1*), 3 (*Scmv2*), and 10 (*Scmv3*) in maize (Jones *et al*., 2011, Lübberstedt *et al*., 2006, Xia *et al*., 1999, Zhang *et al*., 2003). These same loci co-localize in several potyvirus resistant lines, including FAP1360A, Pa405, and Oh1VI which are used in this study (Jones et al., 2011, McMullen, 1994, Lübberstedt et al., 2006, Zambrano *et al*., 2014). These same three loci confer resistance to SCMV, MDMV, and WSMV and it is debated whether these resistance genes are pleiotropic or part of a virus resistance gene cluster (Jones et al., 2011, Lübberstedt et al., 2006). Regardless, these three loci have efficacy against all potyviruses that have been identified as contributing to MLN (Jones et al., 2011, Stewart et al., 2017). While fine mapping and cloning have identified the gene conferring resistance to SCMV at the *Scmv1* locus to be an atypical h-type thioredoxin (*TrxH*)(Liu *et al*., 2017), and at the *Scmv2* locus to be an auxin-binding protein (*auxin-binding protein 1, ABP1*)(Leng *et al*., 2017), the exact mechanism of the resistance remains to be elucidated. Based on observations of virus localization in infected plants, the resistance mechanism for *Scmv2* and *Scmv3* is hypothesized to be movement-based, with resistant plants restricting long-distance virus movement and systemic infection (Jones et al., 2011, Lei, 1986). A causal gene at the *Scmv3* locus has yet to be identified.

Among the causal viruses of MLN, MCMV has been shown to have very little sequence diversity, with >99% nucleotide identity among East African isolates and at least 95% nucleotide identity across all isolates deposited in GenBank from around the globe (Asiimwe *et al*., 2020, Mahuku *et al*., 2015a, Mwatuni *et al*., 2020). In contrast, SCMV is more diverse, with isolates from East Africa showing as low as 90% nucleotide identity with each other and as low as 78% identity with isolates from other countries (Mwatuni et al., 2020, Mahuku et al., 2015a). Some partial SCMV sequences show as little as 65.5% nucleotide identity with other sequences deposited in GenBank (Asiimwe et al., 2020).

This high level of diversity in SCMV and potential differential virulence among isolates may present challenges for breeding durable resistance to the virus. Hence, in this study, we evaluated the efficacy of known sources of potyvirus resistance against diverse SCMV isolates. To this end, we challenged a panel of common potyvirus resistance genes with three diverse SCMV isolates from Rwanda. We also sought to understand how the virulence of SCMV isolates may contribute to overall disease severity in MLN so we screened both potyvirus and MLN resistant maize genotypes with these three isolates in co-infection with MCMV. Finally, we fully sequenced the genomes of the SCMV isolates to get a fuller picture of their genomic diversity and how it may correlate with virulence.

## Materials and Methods

### Virus isolates

The Rwandan SCMV isolates used in this study were collected in 2015 as part of a previous MLN survey (Asiimwe et al., 2020). Collection location information for each isolate used in this study is contained in **Supplementary Table 1**. Briefly, leaf samples collected from fields in Rwanda were lyophilized and shipped to the USDA-ARS Corn, Soybean and Wheat Quality Research Unit in Wooster, Ohio, USA for diagnostics and sequencing. A subset of 11 samples were selected for this study based on phylogenetic analysis performed in Asiimwe et al. (2020); isolates representing each major subgroup/clade of SCMV were selected based on sample availability and quality. The lyophilized tissue was ground in 0.01 M KHPO_4_, pH 7.0 buffer, 1:5 (w/v), combined with carborundum and rub-inoculated onto sorghum cv. Sart ([PI 671955](https://npgsweb.ars-grin.gov/gringlobal/accessiondetail?id=1026580)). Since these samples were originally co-infected with SCMV and MCMV, to isolate SCMV, a serial inoculation series was conducted using sorghum cv. Sart, which is susceptible to SCMV but not MCMV. Immunostrips (Agdia) were used to confirm the presence of SCMV and absence of MCMV before each passage. After four passages, inoculation of the corn inbred Oh28 ([PI 701842](https://npgsweb.ars-grin.gov/gringlobal/accessiondetail?id=1125055)), which is susceptible to both viruses, resulted in only infection by SCMV, indicating that MCMV had been successfully eliminated. Thereafter, these isolates were continually maintained in sorghum cv. Sart through rub-inoculation.

Two other well described SCMV isolates were used in this study for reference: an SCMV isolate originally collected from Ohio, USA (SCMV-OH) and one originally collected from Germany (the Seehausen isolate, SCMV-See, (Fuchs & Grüntzig, 1995)). The virulence of SCMV-OH and SCMV-See on many of the maize genotypes used in this study have been previously characterized (Jones et al., 2011, Zambrano et al., 2014). The SCMV-OH isolate was maintained in the greenhouse through serial rub-inoculation on sorghum cv. Sart as previously described (Jones *et al*., 2007). SCMV-See was generously provided by Stephan Hentrup (Danish Institute of Agricultural Sciences, Slagelse, Denmark) in 2010, placed in long-term storage in liquid nitrogen, vascular puncture inoculated into Oh28 as previously described (Louie, 1995, Bernardo *et al*., 2023), and subsequently maintained through serial rub-inoculation on sorghum cv. Sart. The MCMV isolate used in this study was originally collected from Kansas and maintained through rub-inoculation (Nault *et al*., 1978). Virulence of this MCMV isolate on Oh28 during single infection or co-infection with SCMV-OH has been previously described (Stewart & Willie, 2021, Jones et al., 2018). All experiments were conducted in the growth chamber under containment conditions (APHIS permits P526P-20-00592, P526P-21-03148, P526P-21-06846).

### Maize germplasm

All inbred and near isogenic lines (NILs) used in this study were maintained in fields near Wooster, Ohio, USA. The susceptible check, Oh28, is a dent corn inbred with high SCMV and MCMV susceptibility (Jones et al., 2018, Nault *et al*., 1971, Louie *et al*., 1990). The three potyvirus resistance loci used in this study, *Scmv1* (chromosome 6), *Scmv2* (chromosome 3), and *Scmv3* (chromosome 10), are derived from the dent corn inbred Pa405 ([Ames 22757](https://npgsweb.ars-grin.gov/gringlobal/accessiondetail?id=1517111)), which has elite potyvirus resistance, but is susceptible to MCMV (Jones et al., 2018, Mikel, 1984, McMullen, 1991, McMullen, 1994). NILs carrying the three potyvirus resistance loci in all possible combinations were generated by introgression of Pa405 into the susceptible Oh28 genetic background by marker assisted backcross selection using Pa405 as the donor parent and Oh28 as the recurrent parent. The desired combinations of potyvirus resistance loci were fixed by selfing and selection of homozygous resistant individuals at the BC_6_ generation as previously described (Jones et al., 2011). All NILs are homozygous for the various combinations of *Scmv1*, *Scmv2*, and *Scmv3* loci and their phenotypes and genotypes are confirmed at the time of each seed increase.

Several other virus resistant lines were used in this study. FAP1360A is an early maturing European inbred with good resistance to SCMV (Kuntze *et al*., 2006). Oh1VI ([PI 614734](https://npgsweb.ars-grin.gov/gringlobal/accessiondetail?id=1602987)) is a tropically adapted line selected from the Virgin Island population (PI 504148) with elite SCMV resistance and quantitative MCMV resistance (Louie *et al*., 2002, Jones et al., 2018, Zambrano et al., 2014). KS23-6 was derived from a broad-base synthetic population and has elite MCMV resistance and moderate SCMV resistance (Jones et al., 2018). N211 (PI 596354) was also derived from a synthetic population and has strong MCMV resistance and moderate SCMV resistance (Kaeppler *et al*., 1998).

### SCMV single infection experiments

Single infection experiments were conducted to assess the effect of each of the SCMV isolates on major potyvirus resistance loci. The experiments were conducted in a walk-in growth chamber with 14 hours of light, 26°C day/20°C night temperature, 50% humidity, and a light intensity of 640 umols/m^2^/s. In a preliminary experiment, only the NIL containing *Scmv1* and the parental lines, Pa405 and Oh28, were inoculated with 10 Rwandan isolates as well as the reference isolates SCMV-OH and SCMV-See. Based on the results, 3 Rwandan isolates (SCMV-Rw001, SCMV-Rw043, and SCMV-Rw145) were selected for further experiments using the entire NIL panel, as well as the parental lines (Oh28, Pa405) and two other potyvirus resistant inbreds, FAP1360A and Oh1VI. SCMV-OH and SCMV-See isolates were also included in these experiments for reference. Each experiment was conducted as follows: 6 seeds per line were planted in 10 cm plastic pots. After 10 days, all seedlings that had emerged (V2-V3 growth stage) were rub-inoculated with inoculum of the respective SCMV isolate or mock-inoculated and then inoculated again 2-3 days later. SCMV inoculum consisted of infected and symptomatic sorghum plants 14 dpi, all inoculated at the same age (10 days). Inoculum was prepared by grinding tissue 1:5 (w/v) in 0.01 M KHPO_4_, pH 7.0 buffer plus carborundum. Mock inoculum consisted of just buffer plus carborundum.

Thereafter, symptoms of each individual plant were assessed every 2-3 days for 4 weeks and rated on a 0-2 scale: 0 indicates no symptoms, 1 indicates an incomplete mosaic only covering part of the leaf (delayed limited mosaic) and 2 indicates a full mosaic across the entire leaf (systemic mosaic, **Figure 1**). At 4 weeks post inoculation (wpi), pooled tissue samples were collected from the youngest fully emerged leaf from each plant. The entire experiment was repeated independently twice.

**Figure 1.**
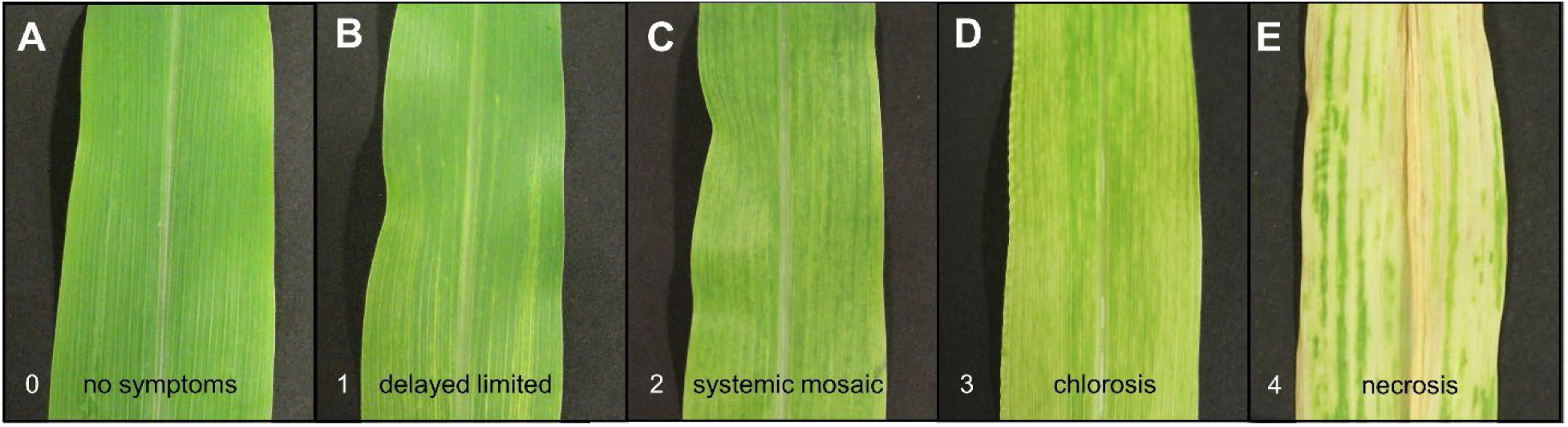
Symptom severity score rating system. Plants were rated on a scale of 0-4 based on symptom severity. Shown are representative photos of leaves exemplifying each score: A) a score of 0 indicates no symptoms, B) a score of 1 indicates mosaic is present but not yet over the entire leaf (delayed limited), C) a score of 2 means mosaic is present across the entire leaf surface (systemic mosaic), D) a score of 3 is given when there is widespread chlorosis present among the mosaic, and D) a score of 4 indicates widespread necrosis. Shown are pictures of inbred Oh28 A) mock-inoculated, B, D, E) infected with SCMV-Rw001 and MCMV or C) singly infected with SCMV-Rw001.

Double-antibody sandwich ELISA was conducted on all pooled samples using a commercially available antibody set for SCMV (Agdia) following the manufacturer’s instructions. Samples were considered positive for SCMV if the absorbance value at 405 nm after 20 minutes of development was more than twice the absorbance of the uninfected plant control. In treatments in which the tissue samples were positive for SCMV via ELISA but no symptoms were observed, SCMV infection was confirmed via RT-PCR using the primers SCMV 8679F and SCMV 959R. RT-PCR was performed following nucleic extraction in grape extraction buffer (GEB) as previously described (Xie *et al*., 2021). Briefly, leaf tissue is ground in GEB (0.015 M sodium carbonate, 0.035 M sodium bicarbonate, 2% polyvinylpyrrolidone, 0.2% bovine serum albumin, 0.05% Tween 20) followed by denaturation in GES (0.1 M glycine-NaOH, pH 9.0, 50 mM NaCl, 1 mM EDTA, 0.5% Triton X-100, 1mM DTT) and used as template in an RT-PCR reaction using Superscript III (ThermoFisher) for RT and Green GoTaq Polymerase (Promega) for PCR. All pooled samples were also tested for the presence of MCMV via RT-PCR using the same method to ensure that MCMV from the original inoculum from Rwanda had been eliminated. The MCMV primers used were MCMV 894F and MCMV 1553R. All primers and their sequences can be found in **Supplementary Table 2**.

In instances where SCMV was detected in potyvirus resistant lines (Oh28*^Scmv1Scmv2Scmv3^*, Pa405, FAP1360A, or Oh1VI), infection by the correct SCMV isolate was confirmed by amplification and Sanger sequencing of the entire coat protein (CP) gene. Tissue was homogenized in TRI Reagent (Zymo Research) and total RNA was extracted using the Direct-zol RNA Miniprep Plus kit (Zymo Research) following manufacturer’s instructions including DNaseI digestion. cDNA synthesis was performed using Superscript III and SCMV 9595R as the gene specific reverse primer. PCR was performed with PrimeSTAR GXL (Takara Bio) according to the manufacturer’s instructions using SCMV 7393F and SCMV 9595R primers. PCR products were purified using the Monarch PCR and DNA Cleanup kit (New England Biolabs) and then Sanger sequenced (Genewiz) using the sequencing primers SCMV 7393F, SCMV 8899R, SCMV 8679F, and SCMV 9595R.

Disease severity ratings were used to calculate a relative area under the disease progress curve (rAUDPC) for each plant and determine the percentage of plants that became symptomatically infected. The inbred lines Pa405 and FAP1360A have variegated leaves that mask symptom detection for the subtle mosaic caused by SCMV (**Supplementary Figure S1**) so no rAUDPC is reported for these lines and the percent infection is determined based on what percent of individual plant samples tested positive via ELISA. For all other lines, percent infection is based on symptoms and only a pooled tissue sample was tested by ELISA.

### SCMV and MCMV co-infection experiments

Co-infection experiments were conducted using each of the SCMV isolates in combination with MCMV to assess maize lethal necrosis disease severity. The experiments were conducted in the same walk-in growth chamber under the same environmental conditions as the single infection experiments. For these experiments, a sub-panel of maize genotypes were used: the doubly virus susceptible inbred Oh28, the NIL containing *Scmv1*, the NIL containing *Scmv1+Scmv2+Scmv3*, and the virus resistant inbreds Pa405, FAP1360A, Oh1VI, KS23-6, and N211. All genotypes were inoculated with each of the 5 SCMV isolates (3 Rwandan isolates, SCMV-OH and SCMV-See) singly or in combination with MCMV. Each experiment also included MCMV single and mock inoculated controls. These experiments followed a similar design to the SCMV-only experiments described above: 6 seeds per line were sown per pot. At 10 days, all emerged seedlings were rub-inoculated with their respective isolate combinations, with a second rub-inoculation 2 days later. For co-infection treatments, 4 parts SCMV inoculum was combined with 1 part MCMV inoculum by volume to compensate for the much higher titer of MCMV. Every 2-3 days for 4 weeks each individual plant was scored for disease severity. The 0-2 rating scale was expanded in these experiments to include a rating of 3 for chlorosis and 4 for necrosis (**Figure 1**), two symptom types that did not occur during single SCMV or MCMV infection in these experiments but are part of the disease progression of maize lethal necrosis. It is also important to note that MCMV in isolation also causes a mosaic symptom and so, in this experiment, symptom ratings of 1 or 2 could be due to one or both viruses (**Supplementary Figure S1**). At 4 wpi, tissue samples were collected for ELISA and RT-PCR as described above. The experiment was conducted independently twice.

ELISA for both viruses was conducted on all samples pooled by treatment using commercially available antibodies for both SCMV and MCMV (Agdia) following manufacturer’s instructions. Unexpected ELISA results were confirmed via RT-PCR as described above for single infection experiments.

### Viral genome sequencing and alignment

The genome sequences of SCMV isolates Rw001, Rw043, and Rw145 were determined by Oxford Nanopore sequencing and assembled using Canu 2.2. by Plasmidsaurus Inc. The near complete genome of each virus was amplified with the primers WX263 and WX264 using PrimeStar GXL polymerase after total RNA extraction and cDNA synthesis as described for Sanger sequencing above. The entire genome of SCMV-Rw001 was subsequently cloned into the pJL89 backbone using HiFi Assembly (New England Biolabs) and the plasmid was sequenced. PCR products amplified using primers WX263 and WX264 were directly sequenced for isolates SCMV-Rw043 and SCMV-Rw145. Isolates SCMV-OH (Accession No. JX188385) and SCMV-See (JX185303) were sequenced previously.

Whole genome alignments were performed and visualized in SnapGene version 7.0.3 using MUSCLE. Nucleotide and amino acid identity and similarity scores of full-length sequences and individual open reading frames (ORFs) was calculated using MacVector version 15.5.3. after ClustalW alignment using default parameters (**Supplementary Table 3**). Nucleotide diversity (Pi) was calculated using DnaSP version 6.12.03 using ClustalW-aligned whole SCMV genome sequences as the input. Annotation of ORFs was determined based on conserved potyvirus protease cleavage sites in the polyprotein (Goh & Hahn, 2021).

### Data analysis

All statistical analyses were conducted in R version 4.1.1 (R Core Team, 2021). The relative AUDPC for each individual plant was calculated using the audpc function in the agricolae package (Mendiburu, 2021). Relative AUDPC (rAUDPC) is the proportion of maximum possible disease severity and is calculating by dividing the AUDPC by the total area of the graph (Fry, 1978, Simko & Piepho, 2012). This normalization was performed so that comparisons can be made across experiments. Statistical comparisons were made between treatments using the Kruskal-Wallis test since the data distribution was not normal but was homoscedastic. If significant, *post hoc* pairwise comparisons were conducted using Dunn’s test and adjusted for multiple pairwise comparisons using the Hommel method (Hommel, 1988). Comparisons were made between isolates within a genotype (**Figures 2A, 3A**), and between genotypes within an isolate (**Figures 2B**, **3B**). Comparisons were conducted using the PMCMRplus package (Pohlert, 2021). Each individual plant was considered a biological replicate.

**Figure 2.**
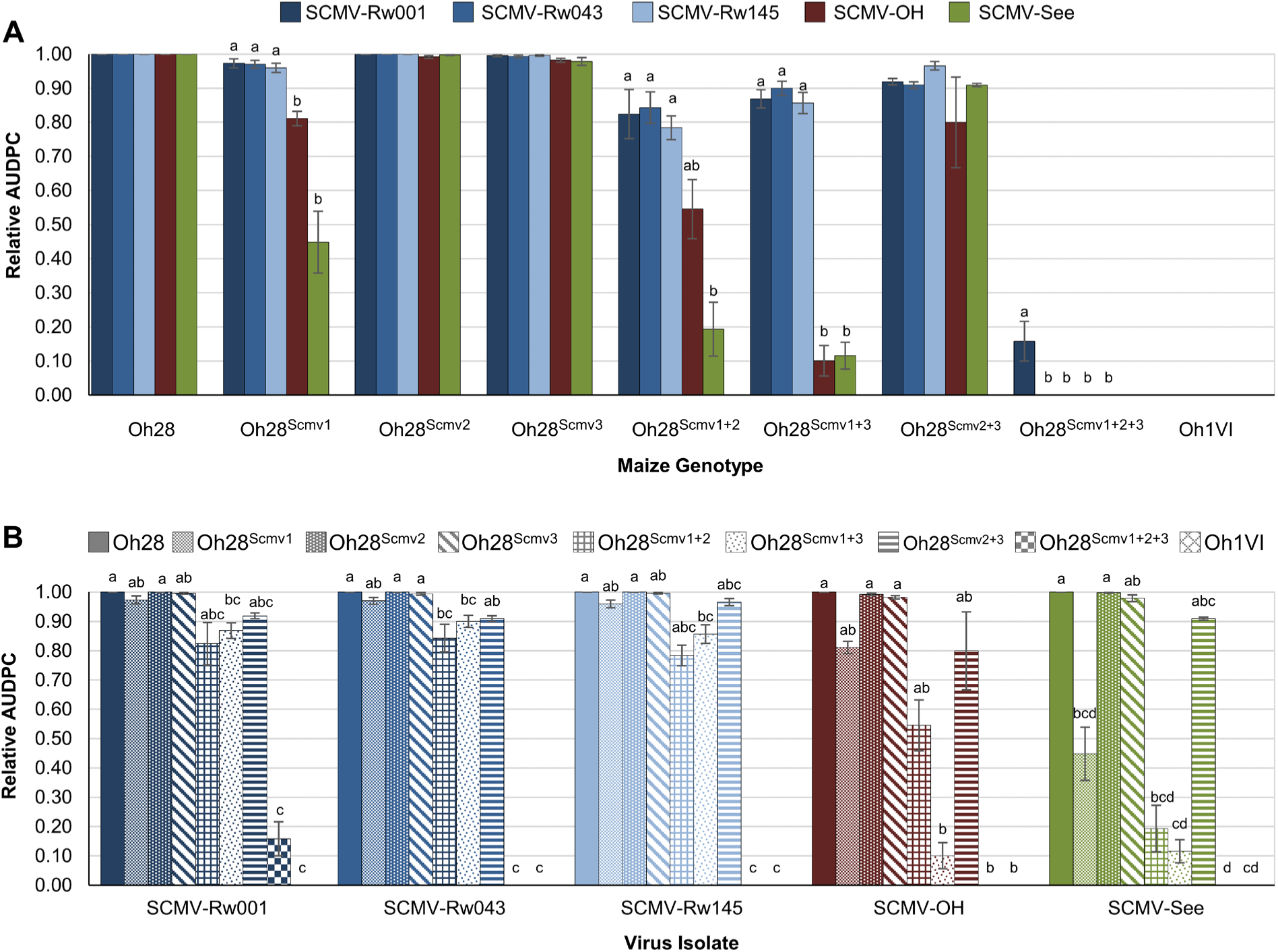
Relative area under the disease progress curve (rAUDPC) for SCMV single infection of the maize potyvirus resistance panel. Maize seedlings were rub-inoculated twice, 2 days apart, at the V3 growth stage with each SCMV isolate or buffer only (mock-inoculated) with 3-5 seedlings inoculated for each isolate-genotype combination. Symptom severity was rated on a 0-2 scale for every plant every 2-3 days for 28 days. The experiment was performed independently twice. Relative AUDPC was calculated for each plant is presented as the proportion of maximum possible disease severity. Bars show mean rAUDPC averaged across all individual plants sorted by maize genotype (A) or by virus isolate (B). Error bars represent ± one standard error across plants. Differences between isolates on each genotype (A) or between genotypes for each isolate (B) were tested using the Kruskal-Wallis test, and if significant, pairwise comparisons between isolates (A) or between genotypes (B) were conducted using Dunn’s test. Letters indicate statistically significant differences between isolates across each genotype (p < 0.05).

**Figure 3.**
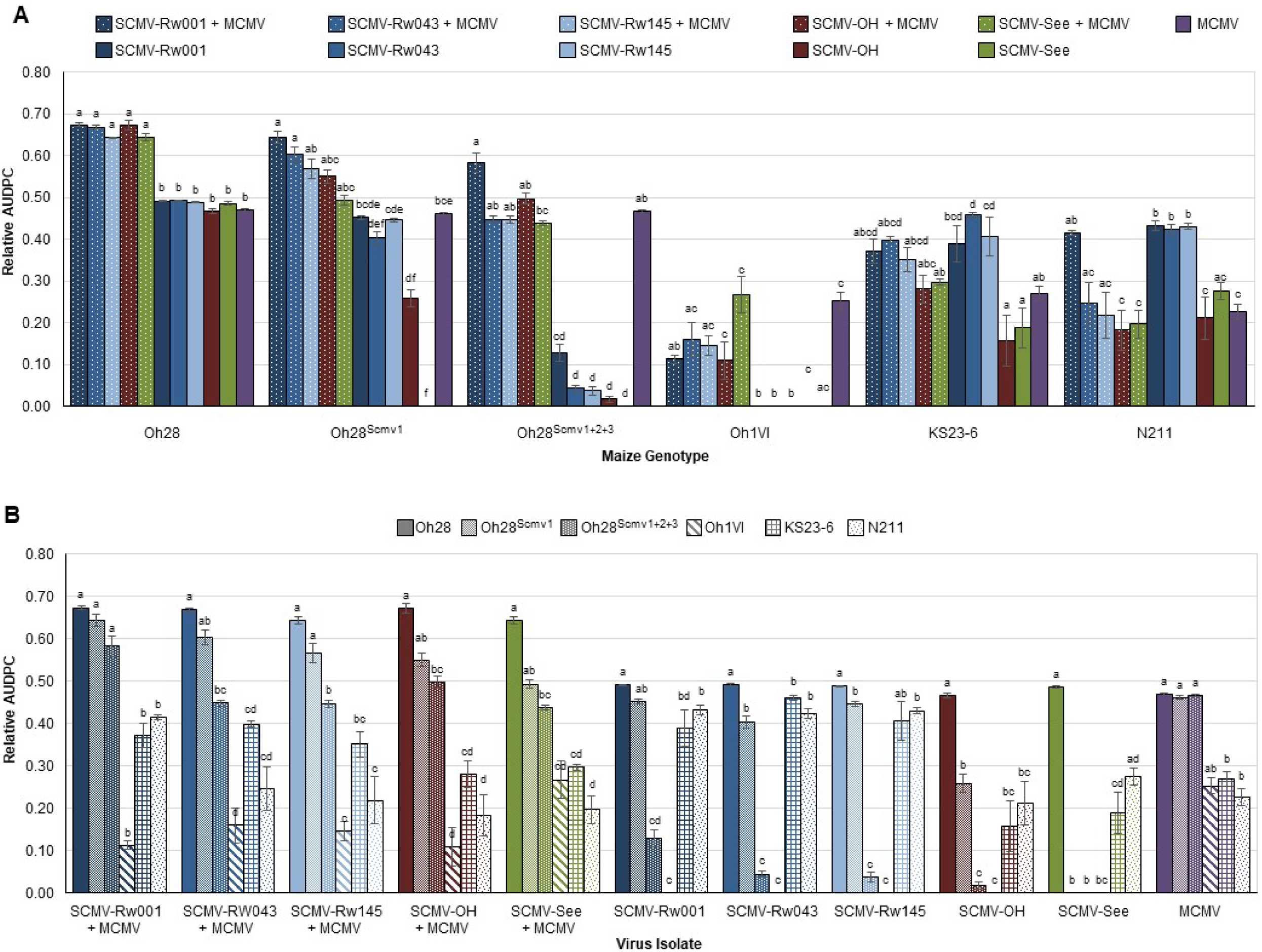
Relative area under the disease progress curve (rAUDPC) for SCMV and MCMV co-infection of a reduced maize potyvirus resistance panel. Maize seedlings were rub-inoculated twice, 2 days apart, at the V3 growth stage with MCMV mixed 1:4 with each SCMV isolate or with buffer only (mock-inoculated). Three to five seedlings were inoculated for each isolate-genotype combination. Symptom severity was rated on a 0-4 scale for every plant every 2-3 days for 28 days. The experiment was performed independently twice. Relative AUDPC was calculated for each plant and is presented as the proportion of maximum possible disease severity. Bars show mean rAUDPC averaged across all individual plants sorted by maize genotype (A) or by virus isolate (B). Error bars represent ± one standard error across plants. Differences between isolates on each genotype (A) or between genotypes for each isolate (B) were tested using the Kruskal-Wallis test, and if significant, pairwise comparisons between isolates (A) or between genotypes (B) were conducted using Dunn’s test. Letters indicate statistically significant differences between isolates across each genotype (p < 0.05).

The relative AUDPC for all individual plants within a treatment were then averaged and graphed.

In all graphs, error bars represent one standard error from the mean and letters indicate significantly different groupings by Dunn’s test with a significance threshold of *P* = 0.05.

## Results

### Single infection of the potyvirus resistance panel with diverse SCMV isolates

In the initial screening, all 10 SCMV isolates from Rwanda that were tested could infect the NIL containing *Scmv1*, with nearly all Rwandan isolates showing a higher rAUDPC than the reference isolates on OH28^Scmv1^, but these differences were only significant for isolate SCMV-Rw145 (**Supplementary Table 3**). Surprisingly, one isolate (SCMV-Rw001) could infect Pa405 as detected via ELISA. Hence SCMV-Rw145, SCMV-Rw001, and one other isolate (SCMV-Rw043) were selected for further study using the full panel of NILs containing all combinations of the potyvirus resistance loci *Scmv1*, *Scmv2*, and *Scmv3*, as well as the resistant donor parent (Pa405), the susceptible recurring parent (Oh28), and two other potyvirus resistant genotypes (FAP1360A and Oh1VI).

In single infection of the potyvirus resistance panel, no significant differences were observed in rAUDPC among SCMV isolates in the susceptible check Oh28, nor the NILs containing only *Scmv2*, only *Scmv3,* or *Scmv2* + *Scmv3* in combination (**Figure 2A**). In the NIL containing just *Scmv1*, all three Rwandan isolates had a significantly higher rAUDPC than both reference isolates, SCMV-OH and SCMV-See. In fact, only lines containing the combination of *Scmv1* and *Scmv3* caused a significant reduction in rAUDPC for all three Rwandan isolates relative to the susceptible check (**Figure 2B**). Even the combination of all three resistance loci was only able to prevent symptom development for two of the three Rwandan isolates, with SCMV-Rw001 still producing symptoms on Oh28*^Scmv1Scmv2Scmv3^*. All three isolates were detected in this line by ELISA and RT-PCR, indicating asymptomatic infection by SCMV-Rw043 and SCMV-Rw145 (**Table 1**).

**Table 1.** The number of symptomatic plants and detection of SCMV in the maize potyvirus resistance panel in single infection experiments.

| Genotype | Oh28 | Oh28<sup>Scmv1</sup> | Oh28<sup>Scmv2</sup> | Oh28<sup>Scmv3</sup> | Oh28<sup>Scmv1+2</sup> | Oh28<sup>Scmv1+3</sup> | Oh28<sup>Scmv2+3</sup> | Oh28<sup>Scmv1+2+3</sup> | Pa405 | FAP1360A | Oh1VI |
| --- | --- | --- | --- | --- | --- | --- | --- | --- | --- | --- | --- |
| **SCMV-Rw001** |  |  |  |  |  |  |  |  |  |  |  |
| Symptomatic/Total | 9/9 | 9/9 | 9/9 | 9/9 | 7/7 | 10/10 | 4/5 | 5/10 | 2/10<sup>c</sup> | 4/5<sup>c</sup> | 0/5 |
| SCMV<sup>a</sup> | + | + | + | + | + | + | + | + | + | + | +<sup>b</sup>/- |
| **SCMV-Rw043** |  |  |  |  |  |  |  |  |  |  |  |
| Symptomatic/Total | 10/10 | 9/9 | 10/10 | 10/10 | 6/6 | 10/10 | 5/5 | 0/10 | 0/10<sup>c</sup> | 2/5<sup>c</sup> | 0/5 |
| SCMV<sup>a</sup> | + | + | + | + | + | + | + | +<sup>b</sup> | - | + | - |
| **SCMV-Rw145** |  |  |  |  |  |  |  |  |  |  |  |
| Symptomatic/Total | 9/9 | 10/10 | 10/10 | 10/10 | 10/10 | 10/10 | 8/8 | 0/10 | 0/10<sup>c</sup> | 4/5<sup>c</sup> | 0/5 |
| SCMV<sup>a</sup> | + | + | + | + | + | + | + | +<sup>b</sup> | - | + | - |
| **SCMV-OH** |  |  |  |  |  |  |  |  |  |  |  |
| Symptomatic/Total | 9/9 | 8/8 | 10/10 | 9/9 | 6/6 | 9/11 | 6/6 | 0/10 | 0/10<sup>c</sup> | 0/5<sup>c</sup> | 0/5 |
| SCMV<sup>a</sup> | + | + | + | + | + | + | + | -/+<sup>b</sup> | - | - | - |
| **SCMV-See** |  |  |  |  |  |  |  |  |  |  |  |
| Symptomatic/Total | 9/9 | 7/9 | 9/9 | 9/9 | 3/3 | 5/10 | 9/9 | 0/10 | 0/10<sup>c</sup> | 0/5<sup>c</sup> | 0/5 |
| SCMV<sup>a</sup> | + | + | + | + | -/+ | -/+ | + | - | - | - | - |
| **Mock** |  |  |  |  |  |  |  |  |  |  |  |
| Symptomatic/Total | 0/9 | 0/6 | 0/8 | 0/10 | 0/6 | 0/10 | 0/4 | 0/10 | 0/9<sup>c</sup> | 0/5<sup>c</sup> | 0/4 |
| SCMV<sup>a</sup> | - | - | - | - | - | - | - | - | - | - | - |

<sup>a</sup> Indicates whether SCMV was detected via ELISA and/or RT-PCR in a pooled sample of all plants per treatment. “+/-” indicates SCMV was detected in the first replicate of the experiment but not in the second replicate of the experiment. “-/+” indicates SCMV was not detected in the first replicate of the experiment but was detected in the second replicate of the experiment.
<sup>b</sup> Indicates asymptomatic infection

Similarly, for the reference isolate SCMV-OH, only *Scmv1* and *Scmv3* in combination led to a significant decrease in rAUDPC relative to the susceptible check (**Figure 2B**), but the Rwandan isolates still had significantly higher disease severity in this line (**Figure 2A**). When inoculated with SCMV-OH, the combination of all three resistance loci abolished symptoms, though asymptomatic infection was detected in Oh28*^Scmv1Scmv2Scmv3^* in the pooled sample collected in the first replicate of the experiment (**Table 1**). In contrast, for the other reference isolate, SCMV-See, *Scmv1* alone was sufficient to significantly reduce disease severity compared to Oh28, with greater decreases observed when stacked with *Scmv2* or *Scmv3* (**Figure 2B**). Infection by SCMV-See was not detected by ELISA when all three loci are present (**Table 1**).

The ELISA and RT-PCR data also indicate that all three Rwandan isolates can consistently infect FAP1360A and that the SCMV-Rw001 isolate can infect Pa405 and Oh1VI, but detection of SCMV-Rw001 in Oh1VI was not consistent across the two single infection experiments (**Table 1**). In comparison, neither SCMV-OH nor SCMV-See were detected in Pa405, FAP1360A, or Oh1VI in any experimental replicate of the single infection experiments (**Table 1**).

### Co-infection of the potyvirus resistance and MLN tolerance panel with MCMV and diverse SCMV isolates

To assess the impact of SCMV virulence on MLN, an informative subset of the potyvirus resistance panel was screened with the three Rwandan SCMV isolates and two reference SCMV isolates in co-infection with MCMV. Single infection of each SCMV isolate as well as single infection by MCMV were included as controls, along with mock inoculation. Two MLN tolerant lines, KS23-6 and N211, were added to the panel for these co-infection experiments. Oh1VI, which was included in the single infection experiments as a potyvirus resistant line also has quantitative resistance to MCMV and is considered MLN tolerant. No differences were observed in rAUDPC in the susceptible line (Oh28) across isolates, in both single SCMV infection and co-infection with MCMV, though co-infection leads to significantly more severe disease (MLN) for all isolates tested, as expected (**Figure 3A**).

When only *Scmv1* was present, no significant differences in rAUDPC were observed among SCMV isolates in co-infection with MCMV (**Figure 3A**) and disease severity was not significantly reduced compared to the susceptible control (**Figure 3B**). This result is despite the fact that this single locus was sufficient to reduce disease progress in several SCMV single infections (**Figure 3B**), as was observed in the single infection experiments for SCMV-See (**Figure 2**). SCMV-Rw001 and SCMV-Rw043 under co-infection with MCMV were the only isolates that caused a significantly higher rAUDPC in this line than MCMV alone, evidence that the additive or synergistic effects induced by co-infection still occurred despite the presence of *Scmv1*.

However, pyramiding several potyvirus resistance loci did lead to a significant decrease in MLN severity, as observed for Oh28*^Scmv1Scmv2Scmv3^*, which had a significantly lower rAUDPC than the susceptible control for all co-infection treatments except MCMV co-infected with SCMV-Rw001 (**Figure 3B**), though for co-infection with this isolate, the rAUDPC was not significantly greater than MCMV infection alone (**Figure 3A**). Importantly, MCMV single infection of this NIL is indistinguishable from the susceptible check, indicating these three loci do not confer resistance to MCMV, and rather the observed reduced MLN disease severity is due to potyvirus resistance alone.

The potyvirus resistant and MCMV tolerant line Oh1VI also significantly reduced MLN severity in all co-infection treatments compared to Oh28 and Oh28*^Scmv1^*, even when co-infected with SCMV-Rw001 (**Figure 3B**). Moreover, for all three Rwandan SCMV isolates tested in co-infection with MCMV, Oh1VI displayed a significantly lower rAUDPC than the NIL containing *Scmv1*, *Scmv2*, and *Scmv3*, which is likely due to its MCMV tolerance or possibly other minor potyvirus resistance loci. SCMV was not detected in any Oh1VI plants in the co-infection experiments by ELISA (**Table 2**), and only detected once by ELISA in the single infection experiments (**Table 1**).

**Table 2.** The number of symptomatic plants and detection of SCMV and in the reduced maize potyvirus resistance panel in co-infection experiments.

| Genotype | Oh28 | Oh28<sup>Scmv1</sup> | Oh28<sup>Scmv1+2+3</sup> | Pa405 | FAP1360A | Oh1VI | KS23-6 | N211 |
| --- | --- | --- | --- | --- | --- | --- | --- | --- |
| **SCMV-Rw001 + MCMV** |  |  |  |  |  |  |  |  |
| Symptomatic/Total | 19/19 | 20/20 | 19/20 | n.a. | n.a. | 7/7 | 10/10 | 10/10 |
| SCMV<sup>a</sup> | + | + | + | + | + | - | + | + |
| MCMV<sup>b</sup> | + | + | + | + | + | + | + | + |
| **SCMV-Rw043 + MCMV** |  |  |  |  |  |  |  |  |
| Symptomatic/Total | 19/19 | 19/19 | 20/20 | n.a. | n.a. | 7/8 | 10/10 | 8/10 |
| SCMV<sup>a</sup> | + | + | + | -/+ | + | - | + | + |
| MCMV<sup>b</sup> | + | + | + | + | + | + | + | + |
| **SCMV-Rw145 + MCMV** |  |  |  |  |  |  |  |  |
| Symptomatic/Total | 19/19 | 15/15 | 20/20 | n.a. | n.a. | 9/10 | 10/10 | 7/10 |
| SCMV<sup>a</sup> | + | + | + | + | + | - | + | + |
| MCMV<sup>b</sup> | + | + | + | + | + | + | + | + |
| **SCMV-OH + MCMV** |  |  |  |  |  |  |  |  |
| Symptomatic/Total | 15/15 | 15/15 | 20/20 | n.a. | n.a. | 4/7 | 10/10 | 8/10 |
| SCMV<sup>a</sup> | + | + | + | -/+ | - | - | -/+ | + |
| MCMV<sup>b</sup> | + | + | + | + | + | + | + | + |
| **SCMV-See + MCMV** |  |  |  |  |  |  |  |  |
| Symptomatic/Total | 15/15 | 20/20 | 20/20 | n.a. | n.a. | 6/6 | 10/10 | 10/10 |
| SCMV<sup>a</sup> | + | + | - | -/+ | - | - | + | + |
| MCMV<sup>b</sup> | + | + | + | + | + | + | + | + |
| **MCMV** |  |  |  |  |  |  |  |  |
| Symptomatic/Total | 18/18 | 20/20 | 20/20 | n.a. | n.a. | 2/2 | 10/10 | 10/10 |
| SCMV<sup>a</sup> | - | - | - | - | - | - | - | - |
| MCMV<sup>b</sup> | + | + | + | + | + | + | + | + |
| **SCMV-Rw001** |  |  |  |  |  |  |  |  |
| Symptomatic/Total | 19/19 | 15/15 | 16/19 | n.a. | n.a. | 0/7 | 9/10 | 10/10 |
| SCMV<sup>a</sup> | + | + | + | - | - | - | + | + |
| MCMV<sup>b</sup> | - | - | - | - | - | - | - | - |
| **SCMV-Rw043** |  |  |  |  |  |  |  |  |
| Symptomatic/Total | 19/19 | 20/20 | 14/18 | n.a. | n.a. | 0/6 | 10/10 | 10/10 |
| SCMV<sup>a</sup> | + | + | + | - | - | - | + | + |
| MCMV<sup>b</sup> | - | - | - | - | - | - | - | - |
| **SCMV-Rw145** |  |  |  |  |  |  |  |  |
| Symptomatic/Total | 19/19 | 19/19 | 9/20 | n.a. | n.a. | 0/10 | 9/10 | 10/10 |
| SCMV<sup>a</sup> | + | + | + | - | + | - | + | + |
| MCMV<sup>b</sup> | - | - | - | - | - | - | - | - |
| **SCMV-OH** |  |  |  |  |  |  |  |  |
| Symptomatic/Total | 19/19 | 20/20 | 5/15 | n.a. | n.a. | 0/7 | 5/10 | 7/10 |
| SCMV<sup>a</sup> | + | + | +/- | -/+ | - | - | +/- | + |
| MCMV<sup>b</sup> | - | - | - | - | - | - | - | - |
| **SCMV-See** |  |  |  |  |  |  |  |  |
| Symptomatic/Total | 19/19 | 0/19 | 0/15 | n.a. | n.a. | 0/6 | 8/10 | 10/10 |
| SCMV<sup>a</sup> | + | - | - | - | - | - | + | + |
| MCMV<sup>b</sup> | - | - | - | - | - | - | - | - |
| **Mock** |  |  |  |  |  |  |  |  |
| Symptomatic/Total | 0/19 | 0/15 | 0/18 | n.a. | n.a. | 0/6 | 0/10 | 0/10 |
| SCMV<sup>a</sup> | - | - | - | - | - | - | - | - |
| MCMV<sup>b</sup> | - | - | - | - | - | - | - | - |

<sup>a</sup> Indicates whether SCMV was detected via ELISA and/or RT-PCR in a pooled sample of all plants per treatment.
“+/-” indicates SCMV was detected in the first replicate of the experiment but not in the second replicate of the experiment. “-/+” indicates SCMV was not detected in the first replicate of the experiment but was detected in the second replicate of the experiment.
<sup>b</sup> Indicates whether MCMV was detected via ELISA and/or RT-PCR in a pooled sample of all plants per treatment.
Abbreviations: SCMV, sugarcane mosaic virus; MCMV, maize chlorotic mottle virus; n.a., not applicable

The two other MLN tolerant lines, KS23-6 and N211, performed similarly to Oh1VI, reducing MLN disease severity relative to the susceptible line in all co-infection treatments, though Oh1VI outperformed these two lines in SCMV single infections, as KS23-6 and N211 decreased SCMV disease severity to a level similar to the *Scmv1* NIL (**Figure 2B**). Moreover, KS23-6 and N211 could be infected by all five SCMV isolates, though KS23-6 infection with SCMV-OH was detected in the pooled sample from one replicate of the experiment but not the other (**Table 2**). Interestingly, SCMV-Rw001 co-infected with MCMV caused significantly higher MLN severity than the reference isolates and a higher rAUDPC than MCMV infection alone on N211 (**Figure 3A**). This result indicates that in the context of intermediate SCMV resistance, the ability of this isolate to overcome potyvirus resistant loci may contribute to more severe disease under co-infection, though disease severity was not significantly higher than SCMV infection alone.

Interestingly, based on RT-PCR and ELISA detection data (**Table 2**), it seems that co-infection with MCMV facilitated SCMV infection of genotypes that the SCMV isolate could not infect alone. For instance, SCMV-Rw043 and SCMV-Rw145 were never detected in Pa405 when inoculated individually across all experiments, but each isolate infected Pa405 when co-inoculated with MCMV, though this was only observed in one of the two co-infection experiments for SCMV-Rw043. Surprisingly, in one instance, the SCMV-See isolate was detected in Pa405 in co-infection with MCMV even though, in single inoculation, sometimes even the presence of just *Scmv1* was enough to prevent infection with SCMV-See. This ELISA result was confirmed by RT-PCR and a region of the SCMV isolate infecting this Pa405 plant was sequenced, confirming it as the SCMV-See isolate.

### Genomic diversity of SCMV isolates

The nearly full-length genome sequences (9631-9673 bp) were determined for the three Rwandan SCMV isolates and compared to the sequences of the two reference isolates, SCMV-OH and SCMV-See. The three isolates from Rwanda were somewhat similar, with an average of 94.7% nucleotide identity between the full-length sequences and were less similar to the reference isolate from Ohio, with 90.6 - 91.4% nucleotide identity (**Table 3**). When looking at amino acid identity, this increases to an average of 97.5% identity among the Rwandan isolates and 97.6 - 98.3% amino acid identity with SCMV-OH. However, SCMV-See is quite different from the other four isolates on the nucleotide level, with only 78.6% identity, on average. Interestingly, on the amino acid level, identity jumps to around 95% between the Seehausen isolate and the isolates from Rwanda but remains just 88.7% amino acid identity with the Ohio isolate (**Table 3**).

**Table 3.** Nucleotide and amino acid identity and similarity scores for the full-length sequences of five SCMV isolates.

| **Virus Isolate** | SCMV-Rw001 | SCMV-Rw043 | SCMV-Rw145 | SCMV-OH | SCMV-See |
| --- | --- | --- | --- | --- | --- |
| **Nucleotide Identity/Similarity (%)<sup>a</sup>** |  |  |  |  |  |
| SCMV-Rw001 | ----- | 94.6 | 94.6 | 91.4 | 78.4 |
| SCMV-Rw043 | 94.6 | ----- | 94.8 | 90.6 | 78.8 |
| SCMV-Rw145 | 94.6 | 94.8 | ----- | 90.7 | 78.6 |
| SCMV-OH | 91.4 | 90.6 | 90.7 | ----- | 78.5 |
| SCMV-See | 78.4 | 78.8 | 78.6 | 78.5 | ----- |
| **Amino Acid Identity/Similarity (%)<sup>b</sup>** |  |  |  |  |  |
| SCMV-Rw001 | ----- | 97.4 | 97.7 | 98.3 | 94.8 |
| SCMV-Rw043 | 98.5 | ----- | 97.5 | 97.6 | 95.2 |
| SCMV-Rw145 | 99.0 | 98.7 | ----- | 98.1 | 94.9 |
| SCMV-OH | 96.8 | 95.8 | 96.1 | ----- | 88.7 |
| SCMV-See | 89.1 | 89.6 | 88.9 | 94.8 | ----- |

<sup>a</sup> Nucleotide identity scores (expressed as a percentage) are presented above the diagonal and nucleotide similarity scores are presented below the diagonal
<sup>b</sup> Amino acid identity scores (expressed as a percentage) are presented above the diagonal and amino acid similarity scores are presented below the diagonal

Nucleotide diversity varies across the genome, with diversity peaking in the middle of the ORF for the P1 protein and in the N-terminus of the CP (**Figure 4**). When broken down by ORF, nucleotide identity between SCMV-See and the other isolates for P1 drops to just below 70%, with 83.1– 90.8% nucleotide identity among Rwandan isolates and 80.8 - 87.4% nucleotide identity between SCMV-OH and the three isolates from Rwanda, with SCMV-Rw145 being the most divergent besides SCMV-See (**Supplementary Table 4**). The lowest nucleotide diversity is at the C-terminus of the CP and along a conserved stretch in the middle of the P3 ORF, which corresponds to the nested Pretty Interesting *Potyviridae* ORF (PIPO, **Figure 4**). Interestingly, both SCMV-See and SCMV-Rw043 share a 39 bp deletion towards the beginning of the CP, which corresponds to a 13 amino acid deletion (**Supplementary Figure S2**). SCMV-See also has a 12 bp insertion in the N-terminal region of the CP ORF that is not shared by any other isolate (**Supplementary Figure S2**).

**Figure 4.**
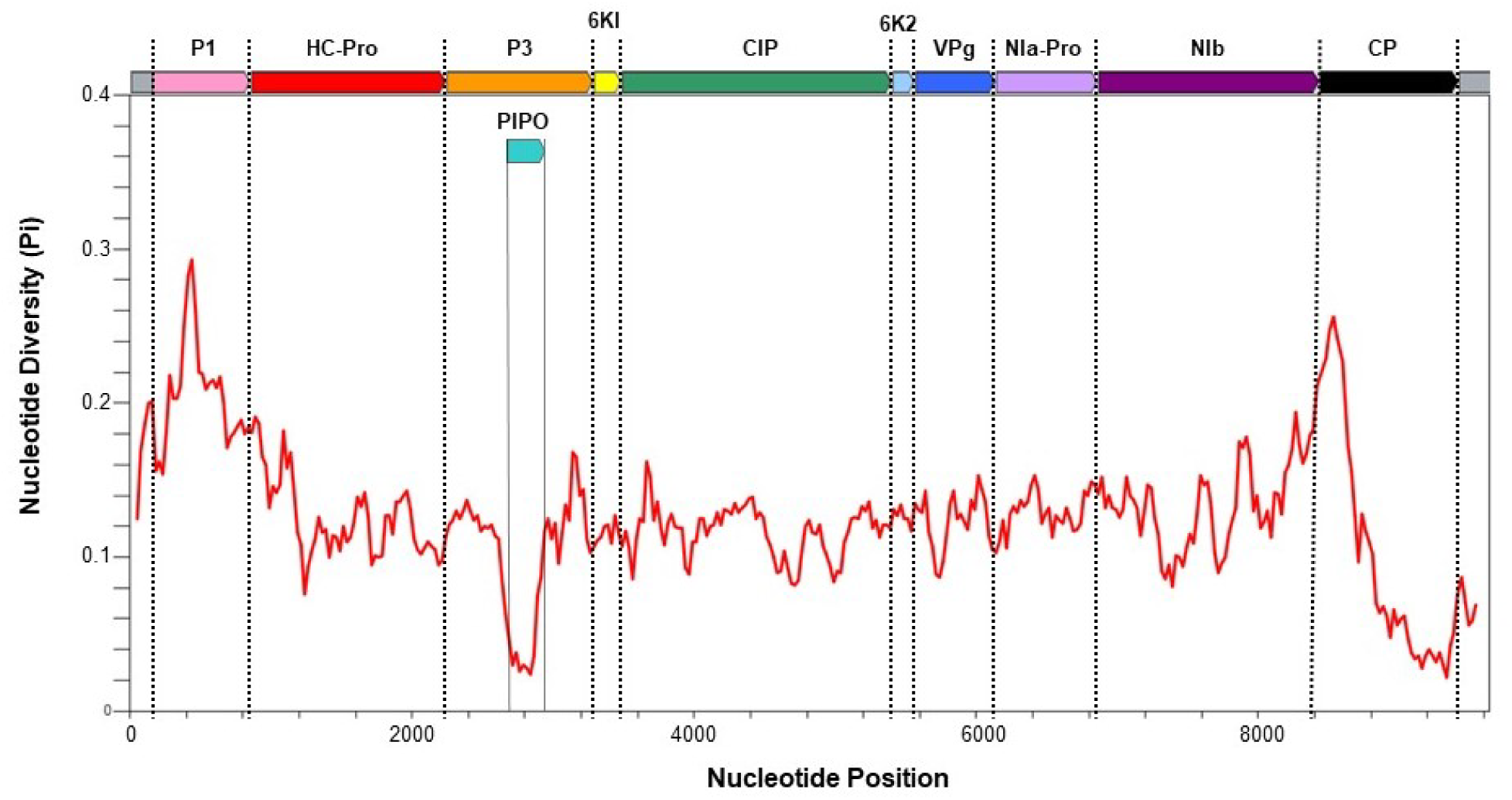
Nucleotide diversity across the genomes of the five SCMV isolates. Graphed is the nucleotide diversity (π), which is the average number of nucleotide differences per site across all possible pairs of sequences, at each position in the SCMV genome. Shown along the top is a diagram of the SCMV genome which shows the 5’ and 3’ untranslated regions (UTR, grey) as well as the open reading frames (ORF) for the viral proteins: P1 (pink), helper component protease (HC-Pro, red), P3 (orange), 6KI (yellow), cylindrical inclusion protein (CIP, green), 6K2 (light blue), viral protein genome-linked (VPg, dark blue), nuclear inclusion a protease (NIa-Pro, light purple), nuclear inclusion b (NIb, dark purple), and coat protein (CP, black) with dotted lines designating the nucleotide position of the start and end of each ORF.

## Discussion

The three SCMV isolates from Rwanda that were characterized show a marked ability to overcome the most widely used potyvirus resistance loci in maize. At least two resistance loci were required to significantly reduce disease severity in single infection, and all three isolates could asymptomatically infect the potyvirus resistant line FAP1360A, which contains all three of the known major potyvirus resistance loci. Isolate SCMV-Rw001 was particularly virulent, and able to infect all elite potyvirus resistant lines tested, though asymptomatically. The results from the SCMV single infection experiments also showed that, even though FAP1360A and Pa405 are purported to have the same three potyvirus resistance loci, the effectiveness of that resistance varies, as has been previously observed (Jones et al., 2011), which may be due to differences in the alleles at those loci or other genetic factors present in each background. The different level of SCMV resistance between the NIL Oh28*^Scmv1Scmv2Scmv3^*and Pa405, from which the loci were derived, also suggests the importance of genetic background as the same exact alleles are present at these loci in both lines. Oh1VI also likely contains the same three resistance loci, as potyvirus resistance in this line has been mapped to the same locations on chromosomes 6, 3, and 10, though finer mapping resolution is needed for confirmation (Zambrano et al., 2014).

Fortunately, the increased virulence of the isolates from Rwanda in SCMV single infection did not generally result in more severe MLN disease when these isolates were co-infected with MCMV, as disease severity was similar between the Rwandan isolates and the two reference isolates which do not have the same ability to overcome the resistance conferred by the three loci. For the NIL containing *Scmv1*, *Scmv2*, and *Scmv3*, disease severity was not significantly different between the co-infection treatments and MCMV infection alone for all SCMV isolates tested, underscoring the effectiveness of potyvirus resistance in reducing MLN severity. This effect of strong potyvirus resistance on suppressing MLN has previously been observed many times, as several potyvirus resistant lines such as Pa405 were reported to have intermediate resistance to MLN despite their MCMV susceptibility (Jones et al., 2018, Ohlson et al., 2022, Doupnik Jr *et al*., 1980, Doupnik *et al*., 1982, Foster, 1984). Importantly, in this study, a combination of potyvirus resistance loci was needed for an effect to be observed, as the incorporation of a single potyvirus resistance locus (*Scmv1*) did not significantly change MLN severity as compared to the susceptible control, highlighting the importance of stacking multiple resistance loci. This result may explain why early MLN tolerant lines that had potyvirus resistance only in the form of *Scmv1* were ineffective at stalling the MLN epidemic in East Africa, especially considering we have shown that SCMV isolates from the region can overcome *Scmv1*-mediated resistance. Resistance gene pyramiding is known to be an effective strategy to increase the durability of resistance and avoid breakdown (Pilet-Nayel *et al*., 2017). However, our study found an SCMV isolate (SCMV-Rw001) that can already overcome the resistance of these three loci when used in combination and hence may produce more severe MLN in the context of incomplete potyvirus resistance, though significantly higher disease severity during MCMV co-infection with this isolate was only observed for one line tested, N211. Therefore, identifying additional sources of potyvirus resistance is desirable to safeguard against these isolates that can evade current potyvirus resistance genes.

In some instances, the incorporation of MCMV resistance may improve the durability of SCMV resistance in co-infection contexts. In this study we observed that MCMV co-infection can facilitate SCMV infection of potyvirus resistant germplasm that the SCMV isolate cannot infect when inoculating singly. Every single SCMV isolate tested was detected at least once in Pa405 when co-inoculated with MCMV, despite the elite potyvirus resistance of this line. However, SCMV infection of Oh1VI was never observed in co-infection experiments and the potyvirus resistance loci in Oh1VI are likely the same as Pa405 – the major difference is the addition of moderate MCMV resistance in Oh1VI, though the effectiveness of the alleles at the major potyvirus resistance loci and potentially additional minor potyvirus resistance loci in Oh1VI may contribute. Even when potyvirus resistance is incomplete, such as in the MLN tolerant lines KS23-6 and N211, which phenotypically resemble lines containing *Scmv1* and *Scmv2* but not *Scmv3* (Gentzel et al., 2024), and hence were infected by all SCMV isolates, the presence of MCMV resistance in these lines still led to a significantly lower MLN disease severity, and they performed similarly to Oh1VI’s complete potyvirus resistance under co-infection when comparing rAUDPC. These results underscore yet again that the most effective and durable path forward in developing MLN resistance is the incorporation of both potyvirus and MCMV resistance genes, ensuring at least two potyvirus resistance loci are present.

An important caveat of these results is that these experiments were performed under growth chambers conditions rather than field conditions. Previous work with these NILs has shown that their resistance to potyviruses, including SCMV, is stronger in the field than in the growth chamber (Jones et al., 2011). This difference is potentially due to the higher pathogen pressure in the growth chamber where adequate inoculation is ensured or the fact that the plants may be stressed by growing in small pots due to space limitations in the growth chamber and thus are more susceptible to infection. Hence, the breakdown of potyvirus resistance, especially when all three loci are present, may be less likely to happen in the field than in these growth chamber experiments.

The sequence diversity among the Rwandan isolates used in this study is consistent with previous studies. The ∼95% nucleotide identity between the three isolates is consistent with previously observed diversity among East African isolates of 87-100% nucleotide identity in near full-length sequences (Mwatuni et al., 2020, Mahuku et al., 2015a). The high nucleotide diversity observed at the end of the NIb ORF and the beginning of the CP is consistent with previous reports that this is a hypervariable region in the SCMV genome (Wamaitha *et al*., 2018, Asiimwe et al., 2020). Large deletions in this region have been previously observed, with six isolates from Kenya sharing the same 39-bp deletion observed in the SCMV-See and SCMV-Rw043 isolates from this study (Wamaitha et al., 2018). The ∼98% amino acid similarity of the Rwandan isolates with the Ohio isolate indicates that small changes in the amino acid sequence of the virus can correspond to large changes in virulence. The mechanism of potyvirus resistance at these loci is still not known and hence how these isolates overcome that resistance is even more obscured. As for the ability of MCMV to facilitate SCMV infection of potyvirus resistant lines, if potyvirus resistance is largely a restriction of long-distance potyvirus movement, then it is possible that MCMV somehow overcomes this barrier and hence SCMV is able to overcome it as well. This enhanced movement would not be unprecedented, as in other plant virus systems it has been shown that one virus can facilitate movement of a different virus in co-infection, allowing the second virus to enter tissues it would not otherwise, with examples for both cell-to-cell and long-distance movement (Harrison *et al*., 1990, Ryabov *et al*., 1999).

The increased virulence of the SCMV isolates tested in the current study not only underscores the importance of stacking multiple sources of resistance but also that breeders must consider the diversity of the virus when breeding for resistance, especially isolates of the virus from the region where the resistance is intended to be deployed. For MLN, the target region for many breeding efforts is East Africa, a large geographic area with high SCMV diversity, including some isolates that can break all major potyvirus resistance loci, as shown in this study. It is unknown to what extent this increased SCMV virulence contributed to the severe MLN epidemics that occurred in this region, especially considering that these isolates did not generally cause heightened MLN disease severity in this study. Nevertheless, combining multiple potyvirus resistance loci with MCMV resistance remains the best path forward towards breeding for durable MLN resistance.

## Acknowledgements

The authors would like to thank Mark Jones for his technical assistance and for developing the near isogenic lines used in this study and Chris Nacci for his diligent maintenance of these lines. This work was supported by United States Department of Agriculture – Agricultural Research Service base funds, project #5082-22000-013-00D.

## Data Availability

The genome sequences of isolates SCMV-Rw001, SCMV-Rw043, and SCMV-Rw145 have been deposited in GenBank on NCBI (PP737788, PP737789, and PP737790, respectively).

---

**Supplementary Table 1.**
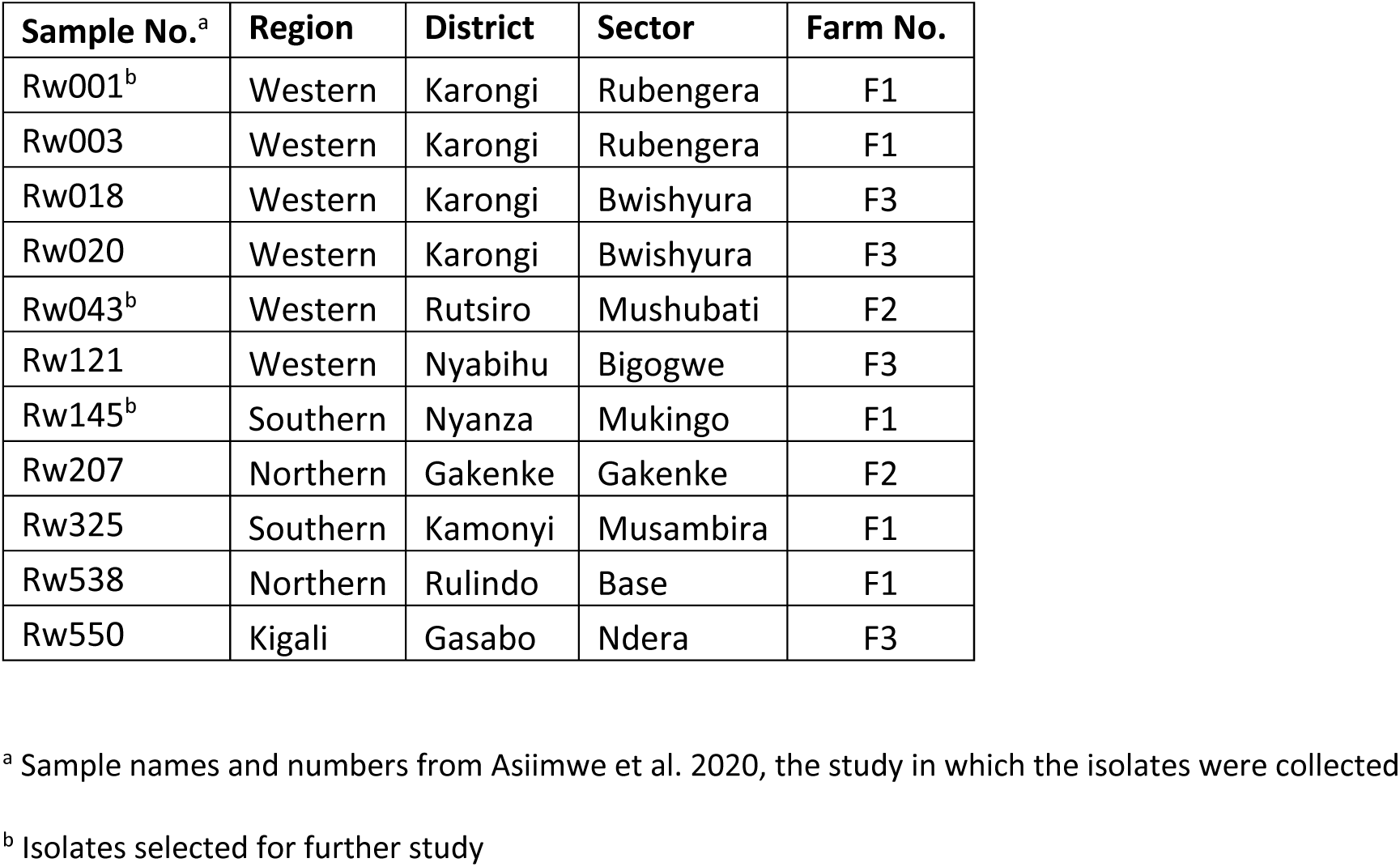
Collection locations of Rwandan SCMV isolates used in this study.

| **Sample No.<sup>a</sup>** | **Region** | **District** | **Sector** | **Farm No.** |
| --- | --- | --- | --- | --- |
| Rw001<sup>b</sup> | Western | Karongi | Rubengera | F1 |
| Rw003 | Western | Karongi | Rubengera | F1 |
| Rw018 | Western | Karongi | Bwishyura | F3 |
| Rw020 | Western | Karongi | Bwishyura | F3 |
| Rw043<sup>b</sup> | Western | Rutsiro | Mushubati | F2 |
| Rw121 | Western | Nyabihu | Bigogwe | F3 |
| Rw145<sup>b</sup> | Southern | Nyanza | Mukingo | F1 |
| Rw207 | Northern | Gakenke | Gakenke | F2 |
| Rw325 | Southern | Kamonyi | Musambira | F1 |
| Rw538 | Northern | Rulindo | Base | F1 |
| Rw550 | Kigali | Gasabo | Ndera | F3 |

<sup>a</sup> Sample names and numbers from Asimwe et al. 2020, the study in which the isolates were collected
<sup>b</sup> Isolates selected for further study

**Supplementary Table 2.**
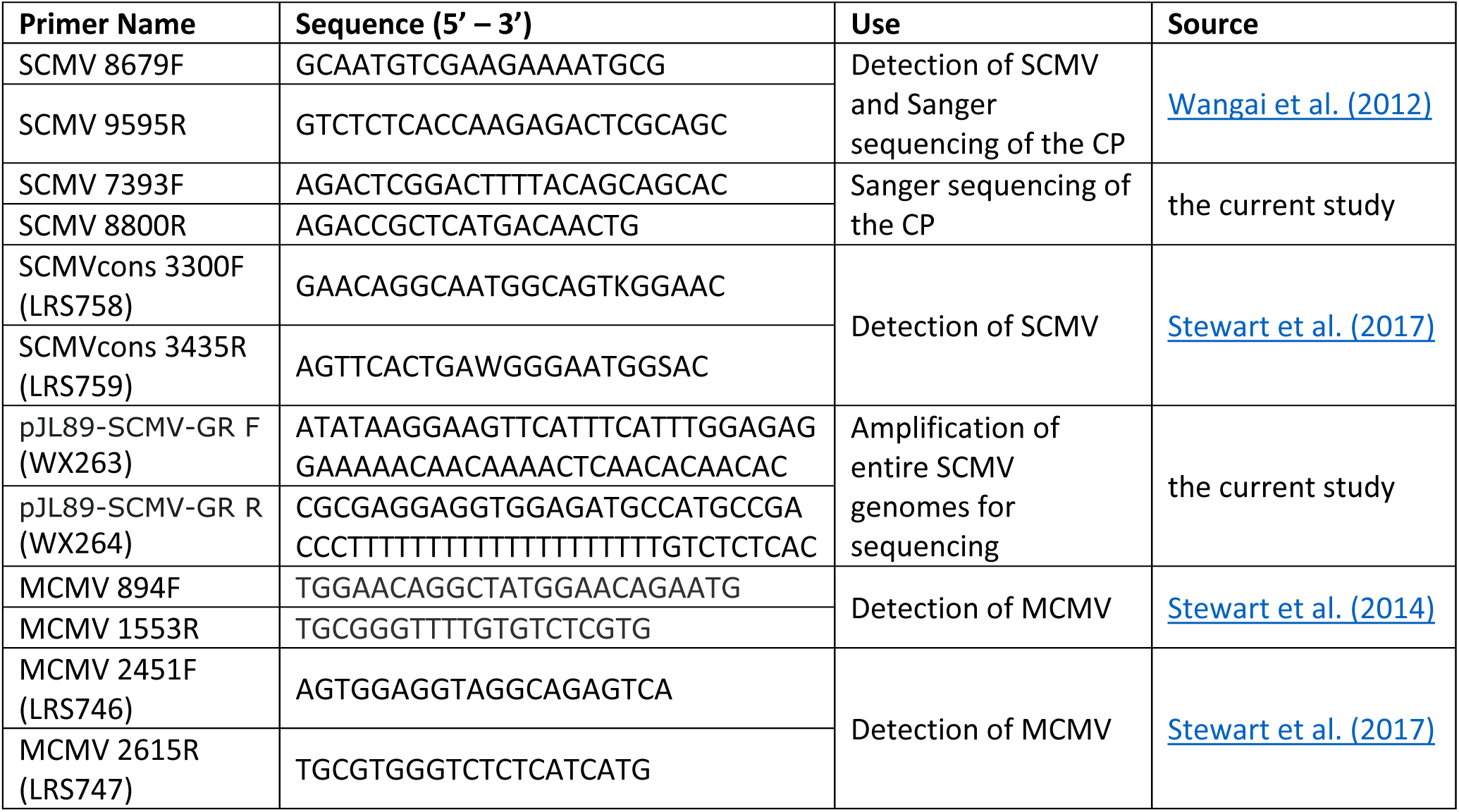
List of all primers used in this study.

**Supplementary Table 3.** Response of the near isogenic line *Scmv1* and parental lines to 10 sugarcane mosaic virus isolates from Rwanda.

| Genotype |  | Oh28 |  |  | Oh28<sup>Scmv1</sup> |  |  | Pa405 |  |
| --- | --- | --- | --- | --- | --- | --- | --- | --- | --- |
| Isolate | Symptomatic<br>/Total | rAUDPC<sup>y</sup> | SCMV<sup>z</sup> | Symptomatic<br>/Total | rAUDPC<sup>y</sup> | SCMV<sup>z</sup> | Symptomatic<br>/Total | rAUDPC<sup>y</sup> | SCMV<sup>z</sup> |
| SCMV-Rw001 | 6/6 | 1.00 ± 0.00 a | + | 5/5 | 0.90 ± 0.01 abc | + | n.a. | n.a. | + |
| SCMV-Rw003 | 5/5 | 1.00 ± 0.00 a | + | 4/4 | 0.89 ± 0.00 abc | + | n.a. | n.a. | - |
| SCMV-Rw018 | 6/6 | 1.00 ± 0.00 a | + | 6/6 | 0.90 ± 0.01 abc | + | n.a. | n.a. | - |
| SCMV-Rw020 | 6/6 | 1.00 ± 0.00 a | + | 4/4 | 0.94 ± 0.01 abc | + | n.a. | n.a. | - |
| SCMV-Rw121 | 4/4 | 1.00 ± 0.00 a | + | 5/5 | 0.91 ± 0.00 ab | + | n.a. | n.a. | - |
| SCMV-Rw145 | 6/6 | 0.97 ± 0.03 a | + | 5/5 | 0.92 ± 0.00 b | + | n.a. | n.a. | - |
| SCMV-Rw207 | 6/6 | 1.00 ± 0.00 a | + | 5/5 | 0.61 ± 0.04 ac | + | n.a. | n.a. | - |
| SCMV-Rw325 | 5/5 | 1.00 ± 0.00 a | + | 5/5 | 0.87 ± 0.01 abc | + | n.a. | n.a. | - |
| SCMV-Rw538 | 4/4 | 1.00 ± 0.00 a | + | 3/3 | 0.91 ± 0.01 abc | + | n.a. | n.a. | - |
| SCMV-Rw550 | 5/5 | 1.00 ± 0.00 a | + | 5/5 | 0.90 ± 0.01 abc | + | n.a. | n.a. | - |
| SCMV-OH | 4/5 | 0.80 ± 0.20 a | + | 3/5 | 0.49 ± 0.20 abc | + | n.a. | n.a. | - |
| SCMV-See | 6/6 | 1.00 ± 0.00 a | - | 0/5 | 0.00 ± 0.00 c | - | n.a. | n.a. | - |
| None (Mock) | 0/6 | 0.00 ± 0.00 b | - | 0/3 | 0.00 ± 0.00 ac | - | n.a. | n.a. | - |

<sup>y</sup> Shown is the mean relative AUDPC ± one standard error from the mean. Letters indicate significant differences among isolates within each treatment via Dunn's test (p < 0.05).
<sup>z</sup> Indicates whether SCMV was detected via ELISA and/or RT-PCR in a pooled sample of all plants per treatment.
Abbreviations: SCMV, sugarcane mosaic virus; n.a., not applicable

**Supplementary Table 4.** Nucleotide identity and similarity scores for each ORF across five SCMV isolates (see supplemental file)

**Supplementary Table 5.** Amino acid identity and similarity scores for each viral protein across five SCMV isolates (see supplemental file)

**Supplementary Figure S1.**
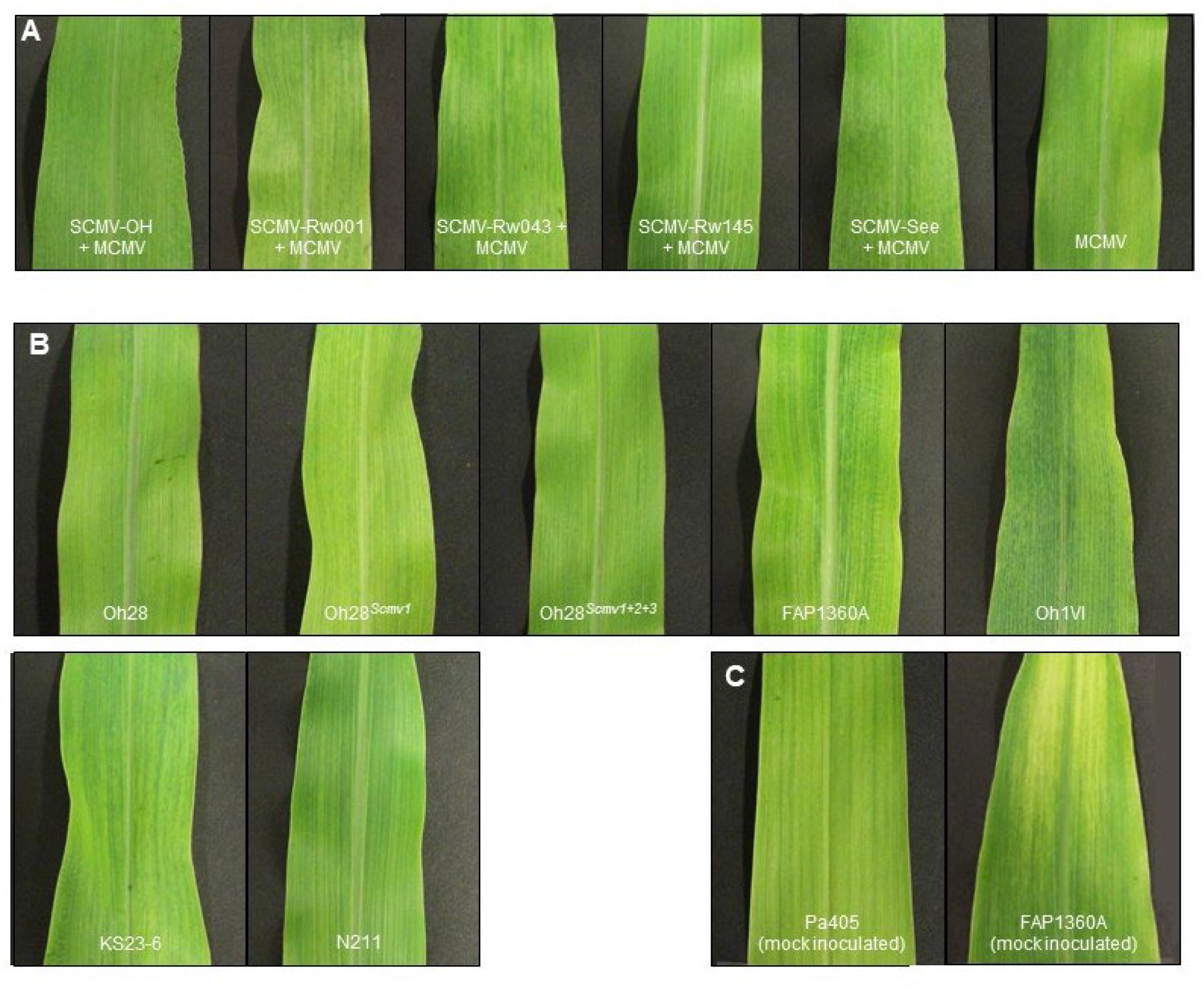
Photos of mosaic symptoms for each isolate and each genotype. Shown are representative photos of mosaic symptoms of A) each SCMV isolate in co-infection with MCMV on the susceptible inbred Oh28 and B) SCMV-Rw001 and MCMV in co-infection on each genotype, except for C) the inbreds Pa405 and FAP1360A, which are variegated even when not inoculated with virus. Mock-inoculated plants are shown for these two inbreds. All plants pictured received a symptom severity score of 2, except for C (unscorable).

**Supplementary Figure S2.** Nucleotide alignment of the near-full genome sequences of the five SCMV isolates. Bases that match the consensus sequence (bold) are shaded pink. Above the consensus sequence is a sequence logo indicating the most prevalent base at each position. Untranslated regions (UTR, grey) and open reading frames of viral genes are indicated below the alignment: P1 (bright pink), helper component protease (HC-Pro, red), P3 (orange), Pretty Interesting *Potyviridae* ORF (PIPO, teal), 6KI (yellow), cylindrical inclusion protein (CIP, green), 6K2 (light blue), viral protein genome-linked (VPg, dark blue), nuclear inclusion a protease (NIa-Pro, light purple), nuclear inclusion b (NIb, dark purple), and coat protein (CP, black). Amino acids are indicated above each position in the polyprotein based on translation of the SCMV-Rw145 sequence. Alignment was performed in SnapGene v 7.0.3 using MUSCLE and default parameters. (see supplemental file)

## Notes

### Competing Interest Statement

The authors have declared no competing interest.

