## Supplementary Figure S2 for "Characterization of three resistance-breaking isolates of sugarcane mosaic virus from Rwanda and implications for maize lethal necrosis"

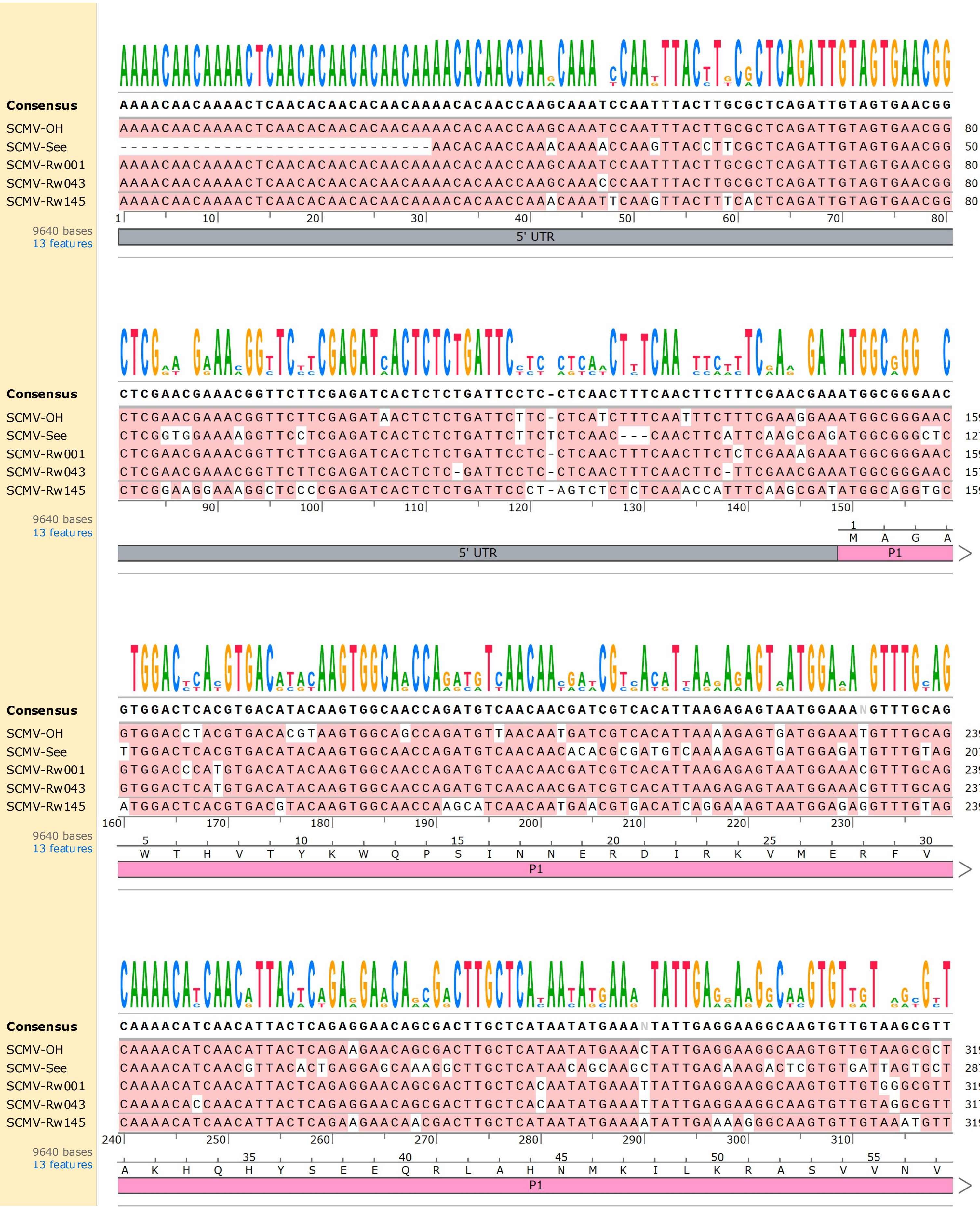

Consensus
SCMV-OH
SCMV-See
SCMV-Rw001
SCMV-Rw043
SCMV-Rw145
9640 bases
13 features
5' UTR
P1

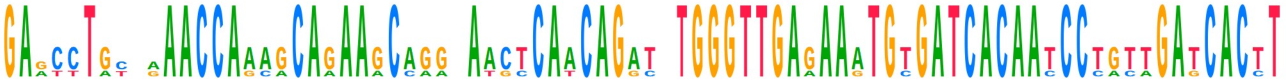

```
Consensus   GAGCCTGCNAAACCAAAGCAGAAGCAGGCAACTCAACAGATGTGGGTTGAGAAATGTGATCACAATCCTGTTGATCACTT
SCMV-OH     GAACCTGCGAAACCAAAGCAGAAGCAGGCAACTCAACAGATGTGGGTTGAGAAATGTGATCACAATCCTGTTGATCACTT  399
SCMV-See    GAGTTTATTGAACCAGCACAGAAACCAAAATGTCATCAGACATGGGTTGAAAAGTGCGATCACAACCCCACAGAGCACTT  367
SCMV-Rw001  GAGCCTGCAAAACCAAAGCAGAAGCAGGCAACTCAACAGATGTGGGTTGAGAAATGTGATCACAATCCTGTTGATCACTT  399
SCMV-Rw043  GAGCCTGCAAAACCAAAGCAGAAGCAGGCAACTCAACAGATGTGGGTTGAGAAATGTGATCACAATCCTGTTGATCACTT  397
SCMV-Rw145  GAACCTGTGGAACCAAAGCAAAAGCAGGTAACCCAACAGGTTTGGGTTGAGAAATGTGATCACAATCCTGTTGATCACCT  399
         320       330       340       350       360       370       380       390
9640 bases      60              65              70              75              80
13 features  E  P  V  E  P  K  Q  K  Q  V  T  Q  Q  V  W  V  E  K  C  D  H  N  P  V  D  H  L
                                         P1
```

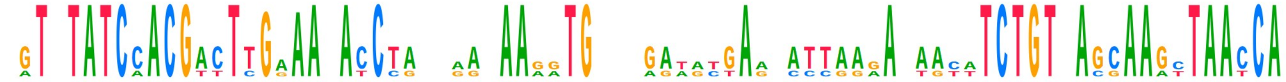

```
Consensus   AGTATATCCACGACTTGAAAGACCTATNAACAAGGTG---GATATGAANATTAAGANAACATCTGTNAGCAAGCTAACCA
SCMV-OH     AGTATATCCACGACTTGGAAGATCCGCAAACAAGGTG---GAAATGAATATTAAAAGAACATCTGTGAGCAAATTAACCA  476
SCMV-See    CATTTATCAACGTTTC---ACACCTAAAAAGAAAGTGCTTAGTACTAAGCCTGAGACAACTTCTGTAACGAAGTTAATCA  444
SCMV-Rw001  AGTATATCCACGACTTGAAAGACCTATCAGCAAAGTG---GAAACGAATATCAGAAGTGCATCTGTGAGCAAGCTAACCA  476
SCMV-Rw043  AGTATATCCACGACTTGAAAGACCTATGGATAAGATG---GATGTGAGCATTAAGAAAATATCTGTTAGCAAGCTAACCA  474
SCMV-Rw145  TGTGTATCCACGACTTGAAAAACCTATGGACAAGATG---GATGTGAGCATTAAGAAAATTTCTGTTAGCAAGCTAACCA  476
         400       410       420       430       440       450       460       470
9640 bases   85              90              95                 100             105
13 features  V  Y  P  R  L  E  K  P  M  D  K  M     D  V  S  I  K  K  I  S  V  S  K  L  T
                                         P1
```

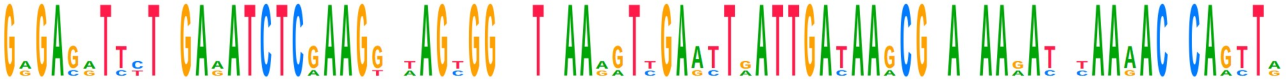

```
Consensus   GAGAGATTCTNGAAATCTCGAAGGCTAGTGGCNTNAAAGTTGAATTGATTGATAAGCGTAAAAGATCTAAAACACAGTTA
SCMV-OH     GGGAGGTTTTAGAGATCTCAAAGTCAAGCGGTCTTAAAGTTGAACTAATTGATAAACGGAAAAGATCCAAAACACAGTTA  556
SCMV-See    GGGACGTCCTTGAAATCTCGAAGGGTAGTGGAATCAAAATTGAGTTAATTGACAAGCGCATAAAACGCAAGACTCAACTA  524
SCMV-Rw001  GAGAGATTTTAGAGATCTCGAAGGTAAGCGGCCTTAAAGTTGAATTGATTGATAAGCGTAAAAGATTTAAAACGCAGTTG  556
SCMV-Rw043  GAGAGATTCTCGAAATCTCGAAGGCTAGTGGCTTGAAGGTCGAATTGATTGATAAGCGTAGAAAATCTAAAACACAGTTA  554
SCMV-Rw145  GAGAGATTCTTGAAATCTCGAAGGCTAGTGGCTTGAAGGTCGAATTGATTGATAAGCGTAAAAGATCTAAAACACAGTTG  556
           480       490       500       510       520       530       540       550
9640 bases  110             115             120             125             130             135
13 features  R  E  I  L  E  I  S  K  A  S  G  L  K  V  E  L  I  D  K  R  K  R  S  K  T  Q  L
                                         P1
```

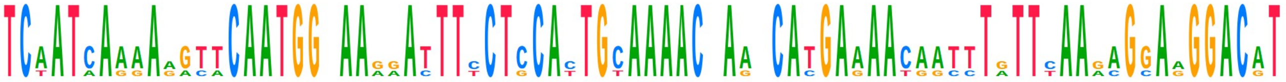

```
Consensus   TCAATCAAAAAGTTCAATGGCAAGGATTTCCTCCACTGCAAAACNAANCATGAAAACAATTTGTTCAAGAGGAAGGACAT
SCMV-OH     TCAATCAAAAGGTTCAATGGTAAAAATTTCCTCCATTGCAAAACGAATCACGAGAACAATTTATTCAAAAGGAGGGACAT  636
SCMV-See    TCTATAAGGAAACACAATGGCAAAGATTTCCTGCATTGCAAAACCAGGCATGAAAATGGCTTGTTCAAACGCAAGGACAT  604
SCMV-Rw001  TCAATCAAAAAGTTCAATGGAAAGAATTTCCTCCACTGTAAAACGAATCATGAAAACAATCTATTCAAGAGGAGGGACGT  636
SCMV-Rw043  TCAATCAAAAAGTTCAATGGCAAGGACTTTCTCCACTGCAAAACAAAACATGAAAACAATTTGTTTAAGAGGAAGGACAT  634
SCMV-Rw145  TCAATCAAAAGGTTCAATGGCAAGGATTTTCTCCACTGCAAAACAAAACATGAAAACAATTTGTTTAAGAGGAAGGACAT  636
           560       570       580       590       600       610       620       630
9640 bases          140             145             150             155             160
13 features  S  I  K  R  F  N  G  K  D  F  L  H  C  K  T  K  H  E  N  N  L  F  K  R  K  D  I
                                         P1
```

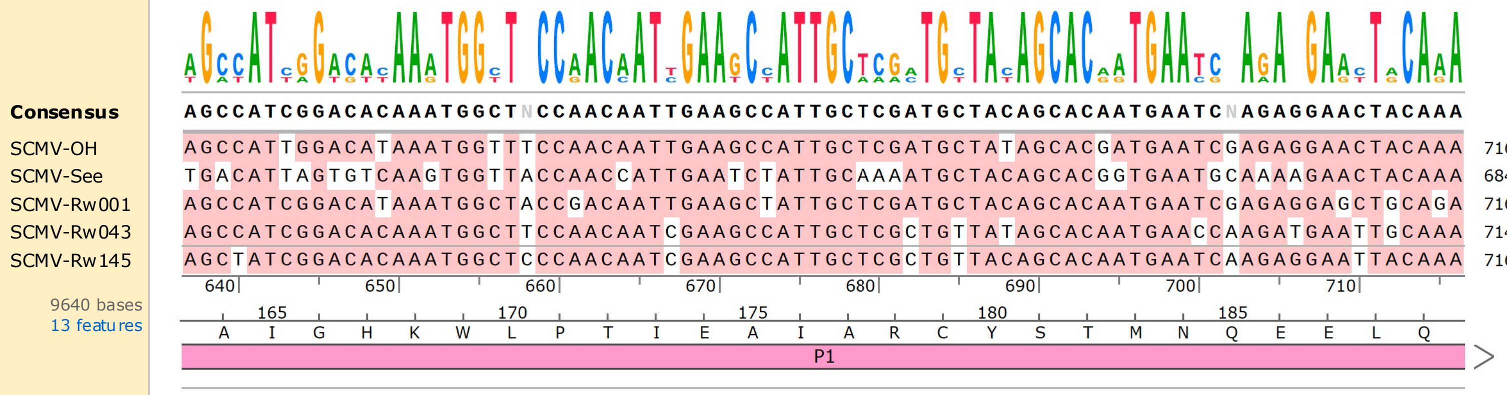
Consensus AGCCATCGGACACAAATGGCTNCCAACAATTGAAGCCATTGCTCGATGCTACAGCACAATGAATCNAGAGGAACTACAAA
SCMV-OH AGCCATTGGACATAAATGGTTTCCAACAATTGAAGCCATTGCTCGATGCTATAGCACGATGAATCGAGAGGAACTACAAA 716
SCMV-See TGACATTAGTGTCAAGTGGTTACCAACCATTGAATCTATTGCAAAATGCTACAGCACGGTGAATGCAAAAGAACTACAAA 684
SCMV-Rw001 AGCCATCGGACATAAATGGCTACCGACAATTGAAGCTATTGCTCGATGCTACAGCACAATGAATCGAGAGGAGCTGCAGA 716
SCMV-Rw043 AGCCATCGGACACAAATGGCTTCCAACAATCGAAGCCATTGCTCGCTGTTATAGCACAATGAACCAAGATGAATTGCAAA 714
SCMV-Rw145 AGCTATCGGACACAAATGGCTCCCAACAATCGAAGCCATTGCTCGCTGTTACAGCACAATGAATCAAGAGGAATTACAAA 716
640 650 660 670 680 690 700 710
9640 bases
13 features
165 170 175 180 185
A I G H K W L P T I E A I A R C Y S T M N Q E E L Q
P1

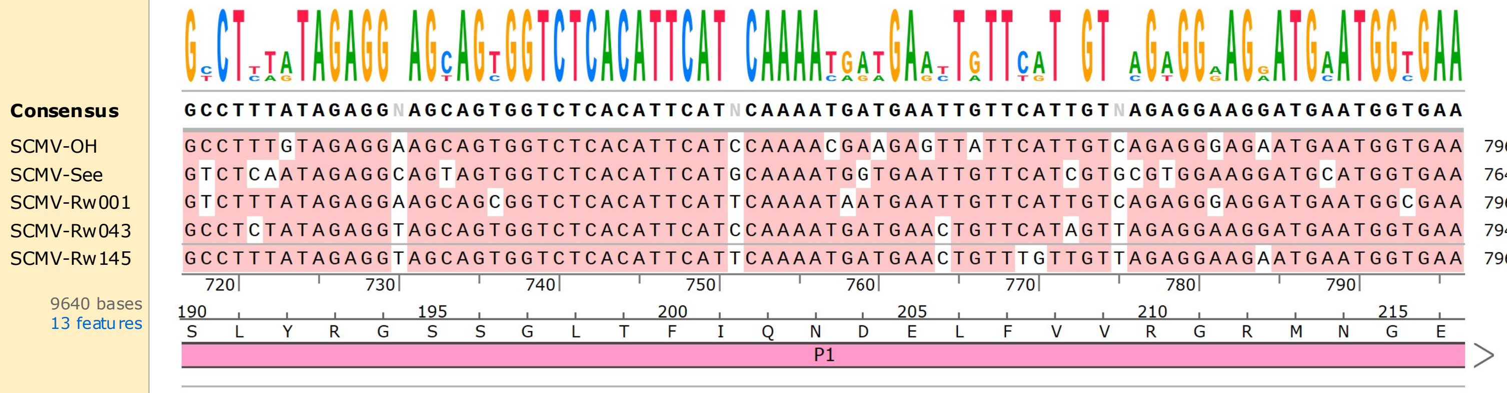
Consensus GCCTTTATAGAGGNAGCAGTGGTCTCACATTCATNCAAAATGATGAATTGTTCATTGTNAGAGGAAGGATGAATGGTGAA
SCMV-OH GCCTTTGTAGAGGAAGCAGTGGTCTCACATTCATCCAAAACGAAGAGTTATTCATTGTCAGAGGGAGAATGAATGGTGAA 796
SCMV-See GTCTCAATAGAGGCAGTAGTGGTCTCACATTCATGCAAAATGGTGAATTGTTCATCGTGCGTGGAAGGATGCATGGTGAA 764
SCMV-Rw001 GTCTTTATAGAGGAAGCAGCGGTCTCACATTCATTCAAAATAATGAATTGTTCATTGTCAGAGGGAGGATGAATGGCGAA 796
SCMV-Rw043 GCCTCTATAGAGGTAGCAGTGGTCTCACATTCATCCAAAATGATGAACTGTTCATAGTTAGAGGAAGGATGAATGGTGAA 794
SCMV-Rw145 GCCTTTATAGAGGTAGCAGTGGTCTCACATTCATTCAAAATGATGAACTGTTTGTTGTTAGAGGAAGAATGAATGGTGAA 796
720 730 740 750 760 770 780 790
9640 bases
13 features
190 195 200 205 210 215
S L Y R G S S G L T F I Q N D E L F V V R G R M N G E
P1

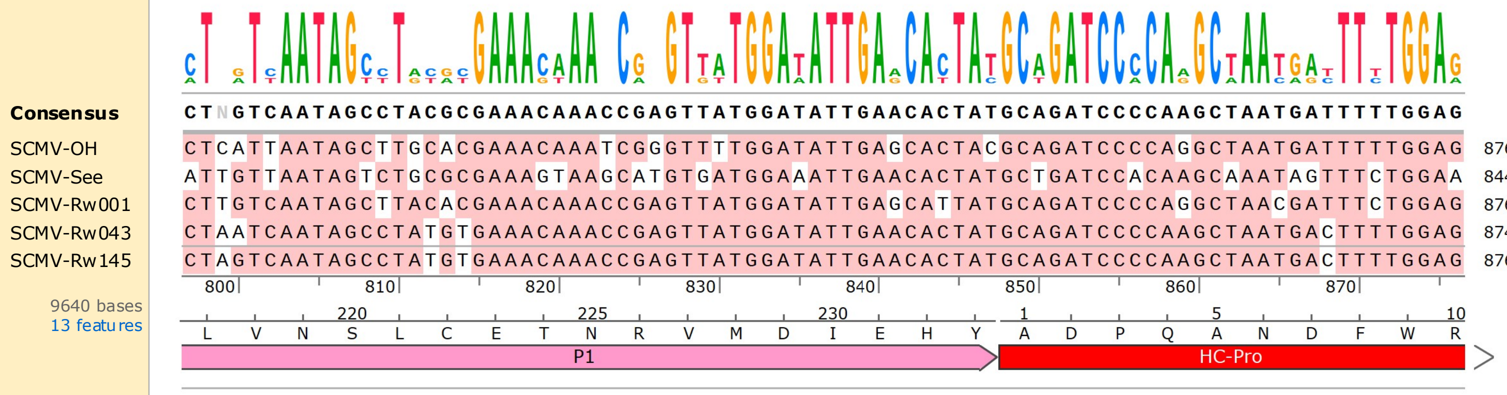
Consensus CTNGTCAATAGCCTACGCGAAACAAACCGAGTTATGGATATTGAACACTATGCAGATCCCCAAGCTAATGATTTTTGGAG
SCMV-OH CTCATTAATAGCTTGCACGAAACAAATCGGGTTTTGGATATTGAGCACTACGCAGATCCCCAGGCTAATGATTTTTGGAG 876
SCMV-See ATTGTTAATAGTCTGCGCGAAAGTAAGCATGTGATGGAAATTGAACACTATGCTGATCCACAAGCAAATAGTTTCTGGAA 844
SCMV-Rw001 CTTGTCAATAGCTTACACGAAACAAACCGAGTTATGGATATTGAGCATTATGCAGATCCCCAGGCTAACGATTTCTGGAG 876
SCMV-Rw043 CTAATCAATAGCCTATGTGAAACAAACCGAGTTATGGATATTGAACACTATGCAGATCCCCAAGCTAATGACTTTTGGAG 874
SCMV-Rw145 CTAGTCAATAGCCTATGTGAAACAAACCGAGTTATGGATATTGAACACTATGCAGATCCCCAAGCTAATGACTTTTGGAG 876
800 810 820 830 840 850 860 870
9640 bases
13 features
220 225 230 1 5 10
L V N S L C E T N R V M D I E H Y A D P Q A N D F W R
P1
HC-Pro

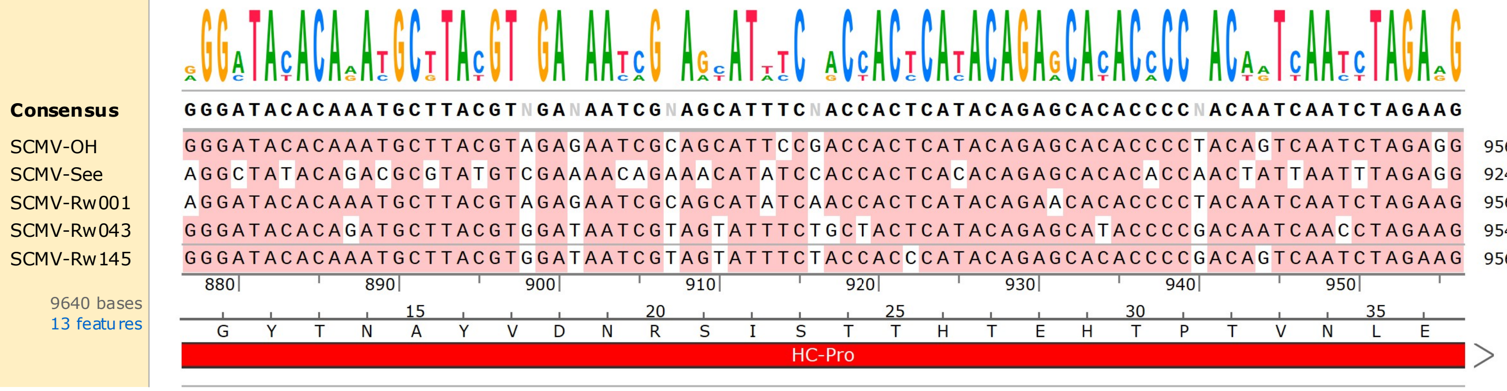
Consensus GGGATACACAAATGCTTACGTNGANAATCGNAGCATTTCNACCACTCATACAGAGCACACCCCNACAATCAATCTAGAAG
SCMV-OH GGGATACACAAATGCTTACGTAGAGAATCGCAGCATTCCGACCACTCATACAGAGCACACCCCTACAGTCAATCTAGAGG 956
SCMV-See AGGCTATACAGACGCGTATGTCGAAAACAGAAACATATCCACCACTCACACAGAGCACACACCAACTATTAATTTAGAGG 924
SCMV-Rw001 AGGATACACAAATGCTTACGTAGAGAATCGCAGCATATCAACCACTCATACAGAAACACACCCCTACAATCAATCTAGAAG 956
SCMV-Rw043 GGGATACACAGATGCTTACGTGGATAATCGTAGTATTTCTGCTACTCATACAGAGCATACCCCGACAATCAACCTAGAAG 954
SCMV-Rw145 GGGATACACAAATGCTTACGTGGATAATCGTAGTATTTCTACCACCCATACAGAGCACACCCCGACAGTCAATCTAGAAG 956
880 890 900 910 920 930 940 950
9640 bases
13 features
15 20 25 30 35
G Y T N A Y V D N R S I S T H T E H T P T V N L E
HC-Pro

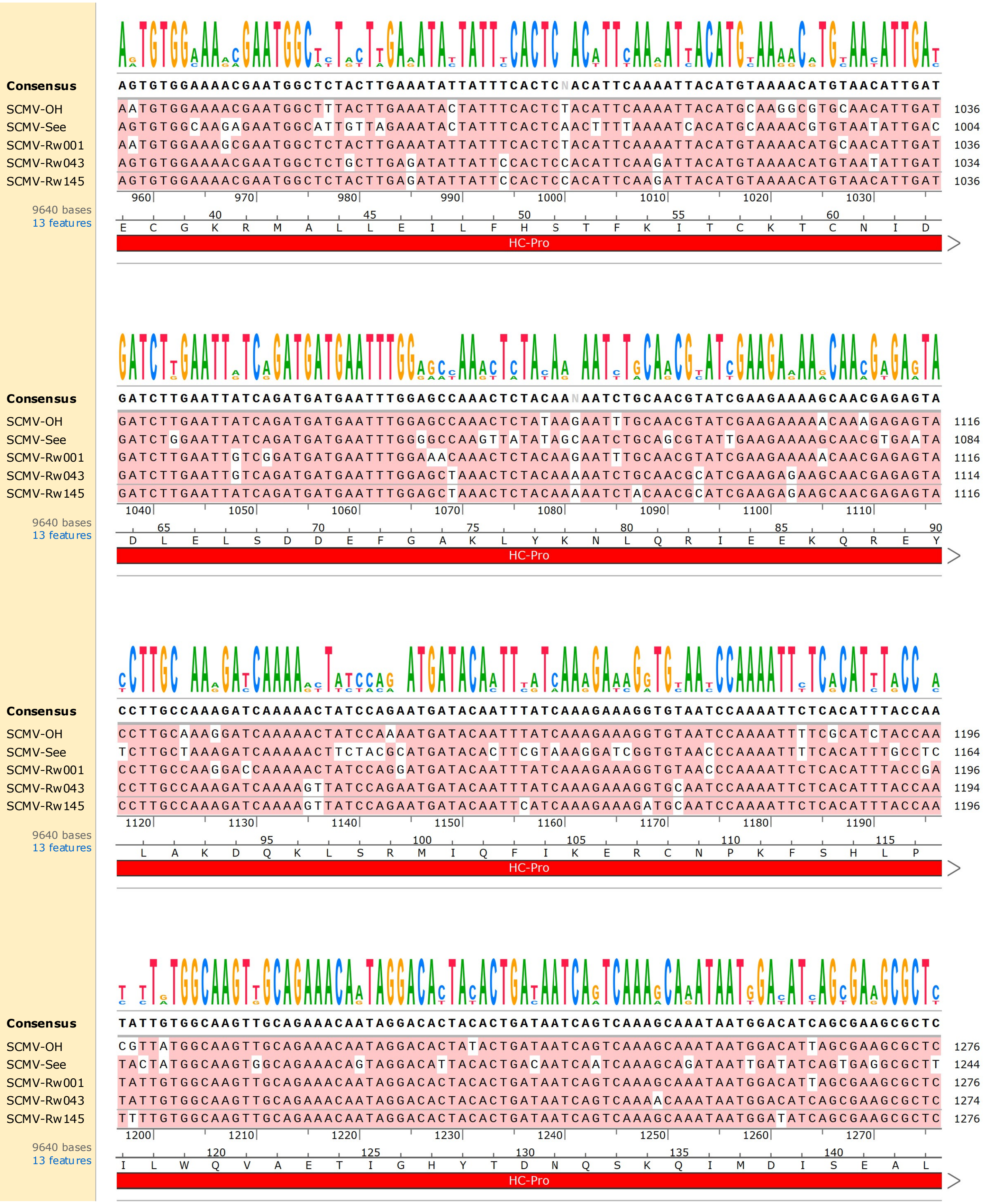

Consensus
AGTGTGGAAAACGAATGGCTCTACTTGAAATATTATTTCACTCNACATTCAAAATTACATGTAAAACATGTAACATTGAT
SCMV-OH AATGTGGAAAACGAATGGCTTTACTTGAAATACTATTTCACTCTACATTCAAAATTACATGCAAGGCGTGCAACATTGAT 1036
SCMV-See AGTGTGGCAAGAGAATGGCATTGTTAGAAATACTATTTCACTCAACTTTTAAAATCACATGCAAAACGTGTAATATTGAC 1004
SCMV-Rw001 AATGTGGAAAGCGAATGGCTCTACTTGAAATATTATTTCACTCTACATTCAAAATTACATGTAAAACATGCAACATTGAT 1036
SCMV-Rw043 AGTGTGGAAAACGAATGGCTCTGCTTGAGATATTATTCCACTCCACATTCAAGATTACATGTAAAACATGTAATATTGAT 1034
SCMV-Rw145 AGTGTGGAAAACGAATGGCTCTACTTGAGATATTATTCCACTCCACATTCAAGATTACATGTAAAACATGTAACATTGAT 1036
960 970 980 990 1000 1010 1020 1030
9640 bases
13 features
40 45 50 55 60
E C G K R M A L L E I L F H S T F K I T C K T C N I D
HC-Pro
Consensus
GATCTTGAATTATCAGATGATGAATTTGGAGCCAAACTCTACAANAATCTGCAACGTATCGAAGAAAAGCAACGAGAGTA
SCMV-OH GATCTTGAATTATCAGATGATGAATTTGGAGCCAAACTCTATAAGAATTTGCAACGTATCGAAGAAAAACAAAGAGAGTA 1116
SCMV-See GATCTGGAATTATCAGATGATGAATTTGGGGCCAAGTTATATAGCAATCTGCAGCGTATTGAAGAAAAGCAACGTGAATA 1084
SCMV-Rw001 GATCTTGAATTGTCGGATGATGAATTTGGAAACAAACTCTACAAGAATTTGCAACGTATCGAAGAAAAACAACGAGAGTA 1116
SCMV-Rw043 GATCTTGAATTGTCAGATGATGAATTTGGAGCTAAACTCTACAAAAATCTGCAACGCATCGAAGAGAAGCAACGAGAGTA 1114
SCMV-Rw145 GATCTTGAATTATCAGATGATGAATTTGGAGCTAAACTCTACAAAAATCTACAACGCATCGAAGAGAAGCAACGAGAGTA 1116
1040 1050 1060 1070 1080 1090 1100 1110
9640 bases
13 features
65 70 75 80 85 90
D L E L S D D E F G A K L Y K N L Q R I E E K Q R E Y
HC-Pro
Consensus
CCTTGCCAAAGATCAAAAACTATCCAGAATGATACAATTTATCAAAGAAAGGTGTAATCCAAAATTCTCACATTTACCAA
SCMV-OH CCTTGCAAAGGATCAAAAACTATCCAAAATGATACAATTTATCAAAGAAAGGTGTAATCCAAAATTTTCGCATCTACCAA 1196
SCMV-See TCTTGCTAAAGATCAAAAACTTCTACGCATGATACACTTCGTAAAGGATCGGTGTAACCCAAAATTTTCACATTTGCCTC 1164
SCMV-Rw001 CCTTGCCAAGGACCAAAAACTATCCAGGATGATACAATTTATCAAAGAAAGGTGTAACCCAAAATTCTCACATTTACCGA 1196
SCMV-Rw043 CCTTGCCAAAGATCAAAAGTTATCCAGAATGATACAATTTATCAAAGAAAGGTGCAATCCAAAATTCTCACATTTACCAA 1194
SCMV-Rw145 CCTTGCCAAAGATCAAAAGTTATCCAGAATGATACAATTCATCAAAGAAAGATGCAATCCAAAATTCTCACATTTACCAA 1196
1120 1130 1140 1150 1160 1170 1180 1190
9640 bases
13 features
95 100 105 110 115
L A K D Q K L S R M I Q F I K E R C N P K F S H L P
HC-Pro
Consensus
TATTGTGGCAAGTTGCAGAAACAATAGGACACTACACTGATAATCAGTCAAAGCAAATAATGGACATCAGCGAAGCGCTC
SCMV-OH CGTTATGGCAAGTTGCAGAAACAATAGGACACTATACTGATAATCAGTCAAAGCAAATAATGGACATTAGCGAAGCGCTC 1276
SCMV-See TACTATGGCAAGTGGCAGAAACAGTAGGACATTACACTGACAATCAATCAAAGCAGATAATTGATATCAGTGAGGCGCTT 1244
SCMV-Rw001 TATTGTGGCAAGTTGCAGAAACAATAGGACACTACACTGATAATCAGTCAAAGCAAATAATGGACATTAGCGAAGCGCTC 1276
SCMV-Rw043 TATTGTGGCAAGTTGCAGAAACAATAGGACACTACACTGATAATCAGTCAAAACAAATAATGGACATCAGCGAAGCGCTC 1274
SCMV-Rw145 TTTTGTGGCAAGTTGCAGAAACAATAGGACACTACACTGATAATCAGTCAAAGCAAATAATGGATATCAGCGAAGCGCTC 1276
1200 1210 1220 1230 1240 1250 1260 1270
9640 bases
13 features
120 125 130 135 140
I L W Q V A E T I G H Y T D N Q S K Q I M D I S E A L
HC-Pro

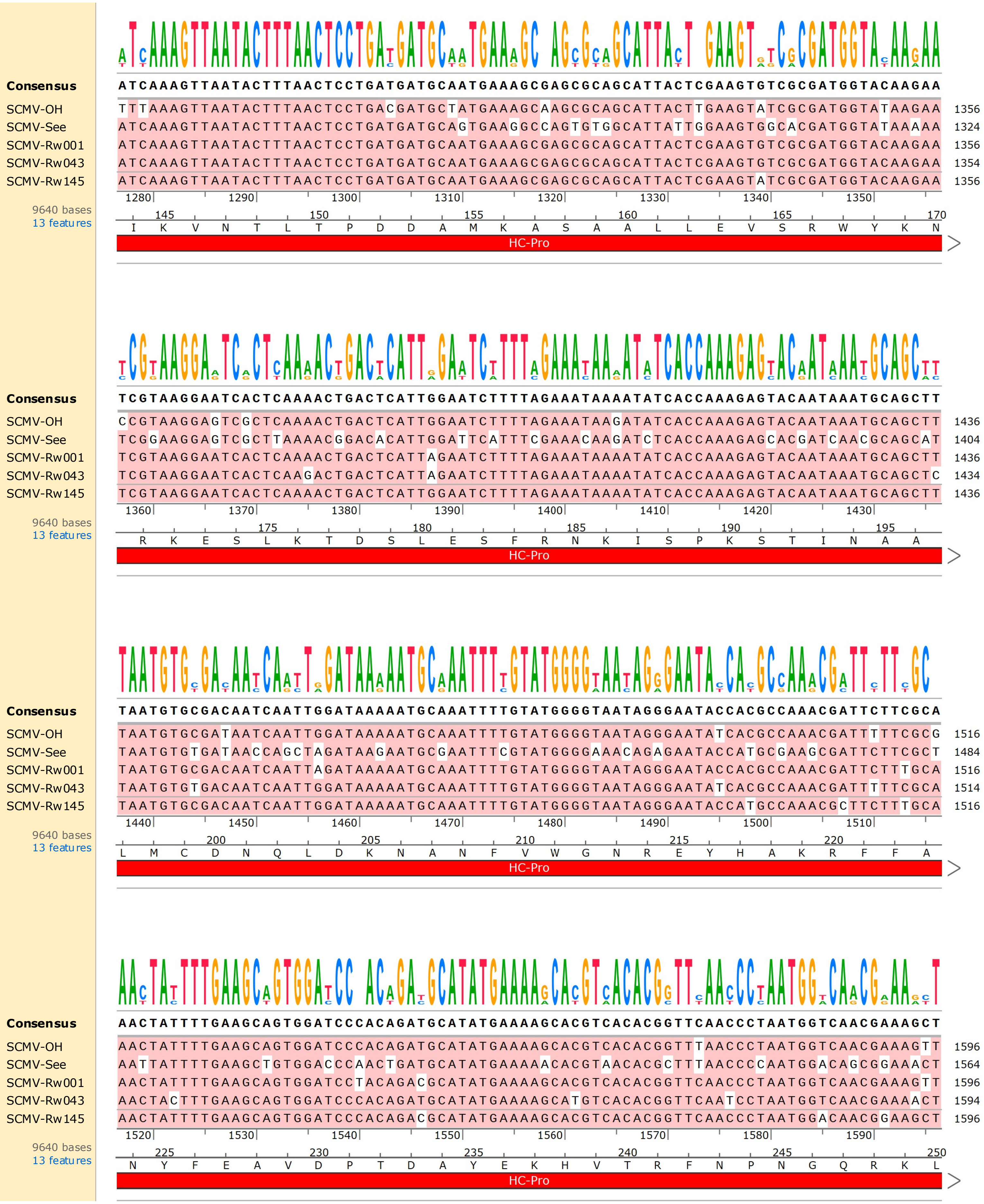
Consensus ATCAAAGTTAATACTTTAACTCCTGATGATGCAATGAAAGCGAGCGCAGCATTACTCGAAGTGTCGCGATGGTACAAGAA
SCMV-OH TTTAAAGTTAATACTTTAACTCCTGACGATGCTATGAAAGCAAGCGCAGCATTACTTGAAGTATCGCGATGGTATAAGAA 1356
SCMV-See ATCAAAGTTAATACTTTAACTCCTGATGATGCAGTGAAGGCCAGTGTGGCATTATTGGAAGTGGCACGATGGTATAAAAA 1324
SCMV-Rw001 ATCAAAGTTAATACTTTAACTCCTGATGATGCAATGAAAGCGAGCGCAGCATTACTCGAAGTGTCGCGATGGTACAAGAA 1356
SCMV-Rw043 ATCAAAGTTAATACTTTAACTCCTGATGATGCAATGAAAGCGAGCGCAGCATTACTCGAAGTGTCGCGATGGTACAAGAA 1354
SCMV-Rw145 ATCAAAGTTAATACTTTAACTCCTGATGATGCAATGAAAGCGAGCGCAGCATTACTCGAAGTATCGCGATGGTACAAGAA 1356
1280 1290 1300 1310 1320 1330 1340 1350
9640 bases
13 features
145 150 155 160 165 170
I K V N T L T P D D A M K A S A A L L E V S R W Y K N
HC-Pro
Consensus TCGTAAGGAATCACTCAAAACTGACTCATTGGAATCTTTTAGAAATAAAATATCACCAAAGAGTACAATAAATGCAGCTT
SCMV-OH CCGTAAGGAGTCGCTCAAAACTGACTCATTGGAATCTTTTAGAAATAAGATATCACCAAAGAGTACAATAAATGCAGCTT 1436
SCMV-See TCGGAAGGAGTCGCTTAAAACGGACACATTGGATTCATTTCGAAACAAGATCTCACCAAAGAGCACGATCAACGCAGCAT 1404
SCMV-Rw001 TCGTAAGGAATCACTCAAAACTGACTCATTAGAATCTTTTAGAAATAAAATATCACCAAAGAGTACAATAAATGCAGCTT 1436
SCMV-Rw043 TCGTAAGGAATCACTCAAGACTGACTCATTAGAATCTTTTAGAAATAAAATATCACCAAAGAGTACAATAAATGCAGCTC 1434
SCMV-Rw145 TCGTAAGGAATCACTCAAAACTGACTCATTGGAATCTTTTAGAAATAAAATATCACCAAAGAGTACAATAAATGCAGCTT 1436
1360 1370 1380 1390 1400 1410 1420 1430
9640 bases
13 features
175 180 185 190 195
R K E S L K T D S L E S F R N K I S P K S T I N A A
HC-Pro
Consensus TAATGTGCGACAATCAATTGGATAAAAATGCAAATTTTGTATGGGGTAATAGGGAATACCACGCCAAACGATTCTTCGCA
SCMV-OH TAATGTGCGATAATCAATTGGATAAAAATGCAAATTTTGTATGGGGTAATAGGGAATATCACGCCAAACGATTTTTCGCG 1516
SCMV-See TAATGTGTGATAACCAGCTAGATAAGAATGCGAATTTCGTATGGGGAAACAGAGAATACCATGCGAAGCGATTCTTCGCT 1484
SCMV-Rw001 TAATGTGCGACAATCAATTAGATAAAAATGCAAATTTTGTATGGGGTAATAGGGAATACCACGCCAAACGATTCTTTGCA 1516
SCMV-Rw043 TAATGTGTGACAATCAATTGGATAAAAATGCAAATTTTGTATGGGGTAATAGGGAATATCACGCCAAACGATTTTTCGCA 1514
SCMV-Rw145 TAATGTGCGACAATCAATTGGATAAAAATGCAAATTTTGTATGGGGTAATAGGGAATACCATGCCAAACGCTTCTTTGCA 1516
1440 1450 1460 1470 1480 1490 1500 1510
9640 bases
13 features
200 205 210 215 220
L M C D N Q L D K N A N F V W G N R E Y H A K R F F A
HC-Pro
Consensus AACTATTTTGAAGCAGTGGATCCCACAGATGCATATGAAAAGCACGTCACACGGTTCAACCCTAATGGTCAACGAAAGCT
SCMV-OH AACTATTTTGAAGCAGTGGATCCCACAGATGCATATGAAAAGCACGTCACACGGTTTAACCCTAATGGTCAACGAAAGTT 1596
SCMV-See AATTATTTTGAAGCTGTGGACCCAACTGATGCATATGAAAAACACGTAACACGCTTTAACCCCAATGGACAGCGGAAACT 1564
SCMV-Rw001 AACTATTTTGAAGCAGTGGATCCTACAGACGCATATGAAAAGCACGTCACACGGTTCAACCCTAATGGTCAACGAAAGTT 1596
SCMV-Rw043 AACTACTTTGAAGCAGTGGATCCCACAGATGCATATGAAAAGCATGTCACACGGTTCAATCCTAATGGTCAACGAAAACT 1594
SCMV-Rw145 AACTATTTTGAAGCAGTGGATCCCACAGACGCATATGAAAAGCACGTCACACGGTTCAACCCTAATGGACAACGGAAGCT 1596
1520 1530 1540 1550 1560 1570 1580 1590
9640 bases
13 features
225 230 235 240 245 250
N Y F E A V D P T D A Y E K H V T R F N P N G Q R K L
HC-Pro

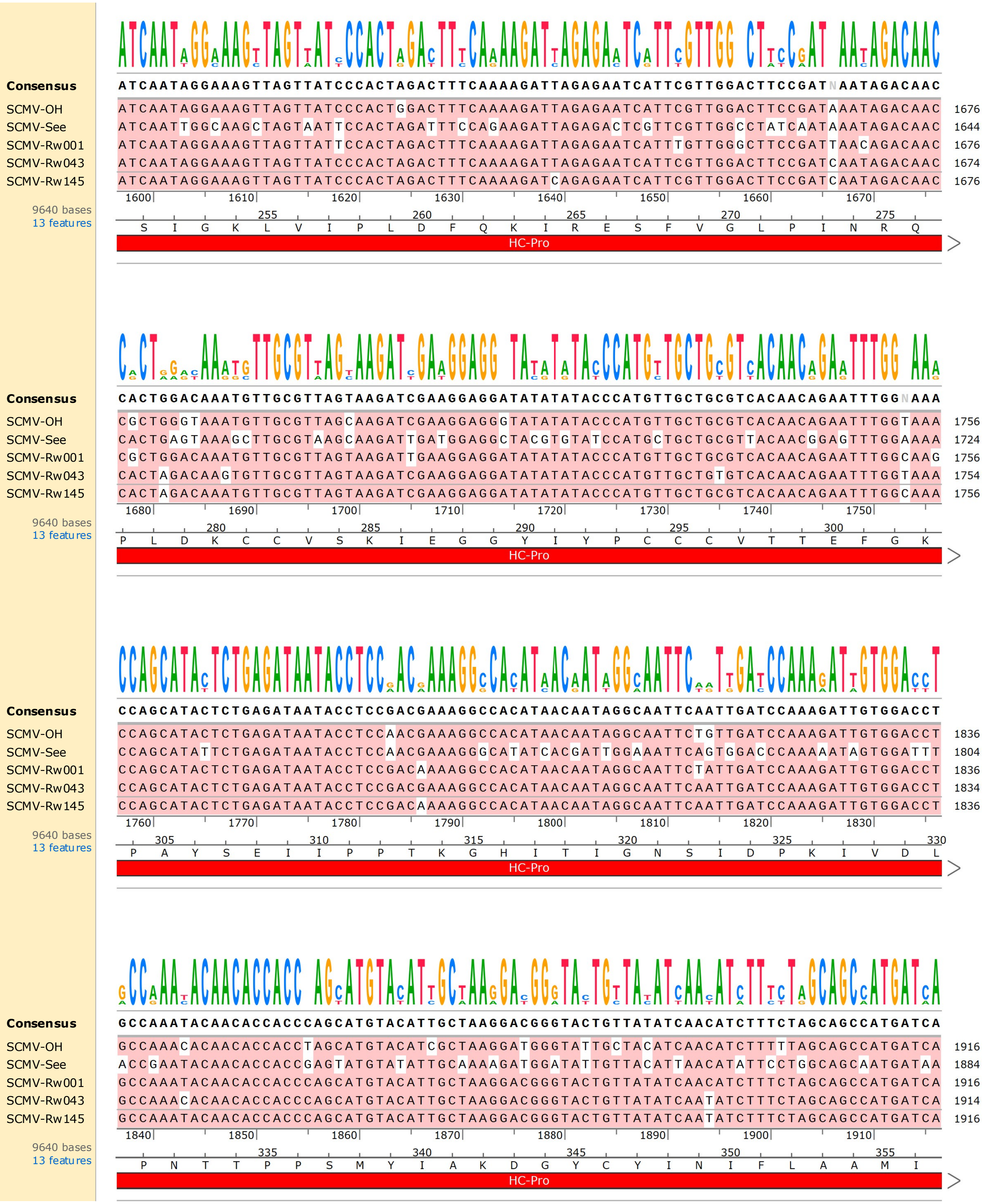
Consensus ATCAATAGGAAAGTTAGTTATCCCACTAGACTTTCAAAAGATTAGAGAATCATTCGTTGGACTTCCGATNAATAGACAAC
SCMV-OH ATCAATAGGAAAGTTAGTTATCCCACTGGACTTTCAAAAGATTAGAGAATCATTCGTTGGACTTCCGATAAATAGACAAC 1676
SCMV-See ATCAATTGGCAAGCTAGTAATTCCACTAGATTTCCAGAAGATTAGAGACTCGTTCGTTGGCCTATCAATAAATAGACAAC 1644
SCMV-Rw001 ATCAATAGGAAAGTTAGTTATTCCACTAGACTTTCAAAAGATTAGAGAATCATTTGTTGGGCTTCCGATTAACAGACAAC 1676
SCMV-Rw043 ATCAATAGGAAAGTTAGTTATCCCACTAGACTTTCAAAAGATTAGAGAATCATTCGTTGGACTTCCGATCAATAGACAAC 1674
SCMV-Rw145 ATCAATAGGAAAGTTAGTTATCCCACTAGACTTTCAAAAGATCAGAGAATCATTCGTTGGACTTCCGATCAATAGACAAC 1676
1600 1610 1620 1630 1640 1650 1660 1670
9640 bases
13 features
255 260 265 270 275
S I G K L V I P L D F Q K I R E S F V G L P I N R Q
HC-Pro
Consensus CACTGGACAAATGTTGCGTTAGTAAGATCGAAGGAGGATATATATACCCATGTTGCTGCGTCACAACAGAATTTGGNAAA
SCMV-OH CGCTGGGTAAATGTTGCGTTAGCAAGATCGAAGGAGGGTATATATACCCATGTTGCTGCGTCACAACAGAATTTGGTAAA 1756
SCMV-See CACTGAGTAAAGCTTGCGTAAGCAAGATTGATGGAGGCTACGTGTATCCATGCTGCTGCGTTACAACGGAGTTTGGAAAA 1724
SCMV-Rw001 CGCTGGACAAATGTTGCGTTAGTAAGATTGAAGGAGGATATATATACCCATGTTGCTGCGTCACAACAGAATTTGGCAAG 1756
SCMV-Rw043 CACTAGACAAGTGTTGCGTTAGTAAGATCGAAGGAGGATATATATACCCATGTTGCTGTGTCACAACAGAATTTGGTAAA 1754
SCMV-Rw145 CACTAGACAAATGTTGCGTTAGTAAGATCGAAGGAGGATATATATACCCATGTTGCTGCGTCACAACAGAATTTGGCAAA 1756
1680 1690 1700 1710 1720 1730 1740 1750
9640 bases
13 features
280 285 290 295 300
P L D K C C V S K I E G G Y I Y P C C V T T E F G K
HC-Pro
Consensus CCAGCATACTCTGAGATAATACCTCCGACGAAAGGCCACATAACAATAGGCAATTCAATTGATCCAAAGATTGTGGACCT
SCMV-OH CCAGCATACTCTGAGATAATACCTCCAACGAAAGGCCACATAACAATAGGCAATTCTGTTGATCCAAAGATTGTGGACCT 1836
SCMV-See CCAGCATATTCTGAGATAATACCTCCAACGAAAGGGCATATCACGATTGGAAATTCAGTGGACCCAAAAATAGTGGATTT 1804
SCMV-Rw001 CCAGCATACTCTGAGATAATACCTCCGACAAAAGGCCACATAACAATAGGCAATTCTATTGATCCAAAGATTGTGGACCT 1836
SCMV-Rw043 CCAGCATACTCTGAGATAATACCTCCGACGAAAGGCCACATAACAATAGGCAATTCAATTGATCCAAAGATTGTGGACCT 1834
SCMV-Rw145 CCAGCATACTCTGAGATAATACCTCCGACAAAAGGCCACATAACAATAGGCAATTCAATTGATCCAAAGATTGTGGACCT 1836
1760 1770 1780 1790 1800 1810 1820 1830
9640 bases
13 features
305 310 315 320 325 330
P A Y S E I I P P T K G H I T I G N S I D P K I V D L
HC-Pro
Consensus GCCAAATACAACACCACCCAGCATGTACATTGCTAAGGACGGGTACTGTTATATCAACATCTTTCTAGCAGCCATGATCA
SCMV-OH GCCAAACACAACACCACCTAGCATGTACATCGCTAAGGATGGGTATTGCTACATCAACATCTTTTTAGCAGCCATGATCA 1916
SCMV-See ACCGAATACAACACCACCGAGTATGTATATTGCAAAAGATGGATATTGTTACATTAACATATTCCTGGCAGCAATGATAA 1884
SCMV-Rw001 GCCAAATACAACACCACCCAGCATGTACATTGCTAAGGACGGGTACTGTTATATCAACATCTTTCTAGCAGCCATGATCA 1916
SCMV-Rw043 GCCAAACACAACACCACCCAGCATGTACATTGCTAAGGACGGGTACTGTTATATCAATATCTTTCTAGCAGCCATGATCA 1914
SCMV-Rw145 GCCAAATACAACACCACCCAGCATGTACATTGCTAAGGACGGGTACTGTTATATCAATATCTTTCTAGCAGCCATGATCA 1916
1840 1850 1860 1870 1880 1890 1900 1910
9640 bases
13 features
335 340 345 350 355
P N T T P P S M Y I A K D G Y C Y I N I F L A A M I
HC-Pro

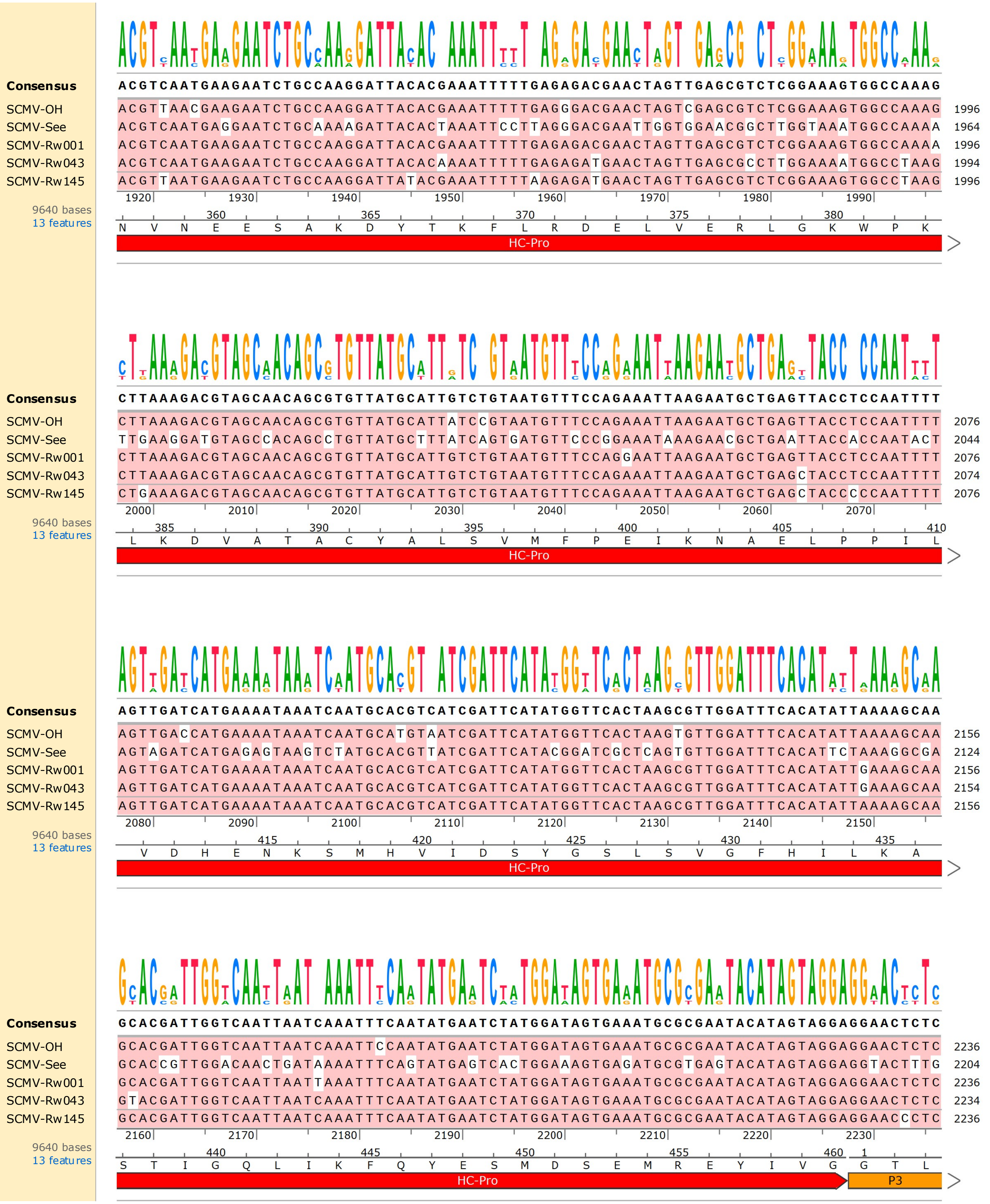
Consensus ACGTCAATGAAGAATCTGCCAAGGATTACACGAAATTTTTGAGAGACGAACTAGTTGAGCGTCTCGGAAAGTGGCCAAAG
SCMV-OH ACGTTAACGAAGAATCTGCCAAGGATTACACGAAATTTTTGAGGGACGAACTAGTCGAGCGTCTCGGAAAGTGGCCAAAG 1996
SCMV-See ACGTCAATGAGGAATCTGCAAAAGATTACACTAAATTCCTTAGGGACGAATTGGTGGAACGGCTTGGTAAATGGCCAAAA 1964
SCMV-Rw001 ACGTCAATGAAGAATCTGCCAAGGATTACACGAAATTTTTGAGAGACGAACTAGTTGAGCGTCTCGGAAAGTGGCCAAAA 1996
SCMV-Rw043 ACGTCAATGAAGAATCTGCCAAGGATTACACAAAATTTTTGAGAGATGAACTAGTTGAGCGCCTTGGAAAATGGCCTAAG 1994
SCMV-Rw145 ACGTTAATGAAGAATCTGCCAAGGATTATACGAAATTTTTAAGAGATGAACTAGTTGAGCGTCTCGGAAAGTGGCCTAAG 1996
1920 1930 1940 1950 1960 1970 1980 1990
9640 bases
13 features
360 365 370 375 380
N V N E E S A K D Y T K F L R D E L V E R L G K W P K
HC-Pro
Consensus CTTAAAGACGTAGCAACAGCGTGTTATGCATTGTCTGTAATGTTTCCAGAAATTAAGAATGCTGAGTTACCTCCAATTTT
SCMV-OH CTTAAAGACGTAGCAACAGCGTGTTATGCATTATCCGTAATGTTTCCAGAAATTAAGAATGCTGAGTTACCTCCAATTTT 2076
SCMV-See TTGAAGGATGTAGCCACAGCCTGTTATGCTTTATCAGTGATGTTCCCGGAAATAAAGAACGCTGAATTACCACCAATACT 2044
SCMV-Rw001 CTTAAAGACGTAGCAACAGCGTGTTATGCATTGTCTGTAATGTTTCCAGGAATTAAGAATGCTGAGTTACCTCCAATTTT 2076
SCMV-Rw043 CTTAAAGACGTAGCAACAGCGTGTTATGCATTGTCTGTAATGTTTCCAGAAATTAAGAATGCTGAGCTACCTCCAATTTT 2074
SCMV-Rw145 CTGAAAGACGTAGCAACAGCGTGTTATGCATTGTCTGTAATGTTTCCAGAAATTAAGAATGCTGAGCTACCCCCAATTTT 2076
2000 2010 2020 2030 2040 2050 2060 2070
9640 bases
13 features
385 390 395 400 405 410
L K D V A T A C Y A L S V M F P E I K N A E L P P I L
HC-Pro
Consensus AGTTGATCATGAAAATAAATCAATGCACGTCATCGATTCATATGGTTCACTAAGCGTTGGATTTCACATATTAAAAGCAA
SCMV-OH AGTTGACCATGAAAATAAATCAATGCATGTAATCGATTCATATGGTTCACTAAGTGTTGGATTTCACATATTAAAAGCAA 2156
SCMV-See AGTAGATCATGAGAGTAAGTCTATGCACGTTATCGATTCATACGGATCGCTCAGTGTTGGATTTCACATTCTAAAGGCGA 2124
SCMV-Rw001 AGTTGATCATGAAAATAAATCAATGCACGTCATCGATTCATATGGTTCACTAAGCGTTGGATTTCACATATTGAAAGCAA 2156
SCMV-Rw043 AGTTGATCATGAAAATAAATCAATGCACGTCATCGATTCATATGGTTCACTAAGCGTTGGATTTCACATATTGAAAGCAA 2154
SCMV-Rw145 AGTTGATCATGAAAATAAATCAATGCACGTCATCGATTCATATGGTTCACTAAGCGTTGGATTTCACATATTAAAAGCAA 2156
2080 2090 2100 2110 2120 2130 2140 2150
9640 bases
13 features
415 420 425 430 435
V D H E N K S M H V I D S Y G S L S V G F H I L K A
HC-Pro
Consensus GCACGATTGGTCAATTAATCAAATTTCAATATGAATCTATGGATAGTGAAATGCGCGAATACATAGTAGGAGGAACTCTC
SCMV-OH GCACGATTGGTCAATTAATCAAATTCCAATATGAATCTATGGATAGTGAAATGCGCGAATACATAGTAGGAGGAACTCTC 2236
SCMV-See GCACCGTTGGACAACTGATAAAATTTCAGTATGAGTCACTGGAAAGTGAGATGCGTGAGTACATAGTAGGAGGTACTTTG 2204
SCMV-Rw001 GCACGATTGGTCAATTAATTAAATTTCAATATGAATCTATGGATAGTGAAATGCGCGAATACATAGTAGGAGGAACTCTC 2236
SCMV-Rw043 GTACGATTGGTCAATTAATCAAATTTCAATATGAATCTATGGATAGTGAAATGCGCGAATACATAGTAGGAGGAACTCTC 2234
SCMV-Rw145 GCACGATTGGTCAATTAATCAAATTTCAATATGAATCTATGGATAGTGAAATGCGCGAATACATAGTAGGAGGAACCCTC 2236
2160 2170 2180 2190 2200 2210 2220 2230
9640 bases
13 features
440 445 450 455 460 1
S T I G Q L I K F Q Y E S M D S E M R E Y I V G G T L
HC-Pro
P3

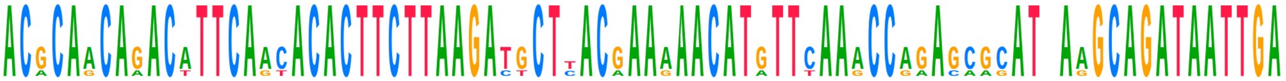

```
Consensus    ACGCAACAGACATTCAACACACTTCTTAAGATGCTTACGAAAAACATGTTCAAACCAGAGCGCATTAAGCAGATAATTGA
SCMV-OH      ACGCAACAGACATTCAACACACTTCTTAAGATGCTTACGAAAAACATGTTCAAACCAGAGCGCATCAAGCAGATAATTGA  2316
SCMV-See     ACACAGCAAACTTTCAGTACACTTCTTAAGACTCTCACAAAGAACATGTTTAAGCCGAACAAGATAAGGCAGATAATTGA  2284
SCMV-Rw001   ACGCAACAGACATTCAACACACTTCTTAAGATGCTCACGAAAAACATGTTCAAACCAGAGCGCATTAAGCAGATAATTGA  2316
SCMV-Rw043   ACGCAACAGACATTCAACACACTTCTTAAGATGCTTACGAAAAACATATTCAAACCAGAGCGCATTAAGCAGATAATTGA  2314
SCMV-Rw145   ACGCAACAGACATTCAACACACTTCTTAAGATGCTTACGAAAAACATGTTCAAACCAGAGCGCATTAAGCAGATAATTGA  2316
             2240  2250  2260  2270  2280  2290  2300  2310
9640 bases   5  10  15  20  25  30
13 features  T Q Q T F N T L L K M L T K N M F K P E R I K Q I I E
             P3
```

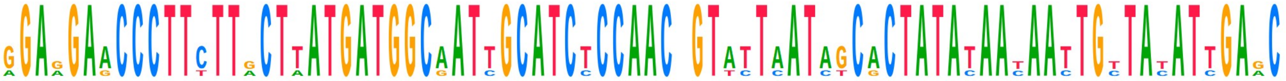

```
Consensus    GGAGGAACCCTTCTTGCTTATGATGGCAATTGCATCTCCAACGGTATTAATAGCACTATATAATAATTGTTATATTGAGC
SCMV-OH      AGAGGAACCCTTTTTACTTATGATGGCGATTGCATCTCCAACGGTATTAATAGCACTATATAATAATTGTTATATTGAGC  2396
SCMV-See     GGAAGAGCCCTTCTTACTAATGATGGCAATTGCATCTCCAACTGTACTCATCTCGCTATACAACAACTGCTACATCGAAC  2364
SCMV-Rw001   GGAGGAACCCTTCTTGCTTATGATGGCAATCGCATCCCCAACGGTATTAATAGCACTATATAACAATTGCTACATTGAGC  2396
SCMV-Rw043   GGAAGAACCCTTCTTGCTTATGATGGCAATTGCATCTCCAACAGTATTAATAGCACTATATAATAATTGTTATATTGAAC  2394
SCMV-Rw145   GGAGGAACCCTTCTTGCTTATGATGGCAATTGCATCTCCAACGGTTTTAATAGCACTATATAATAATTGTTATATTGAGC  2396
             2320  2330  2340  2350  2360  2370  2380  2390
9640 bases   35  40  45  50  55
13 features  E E P F L L M M A I A S P T V L I A L Y N N C Y I E
             P3
```

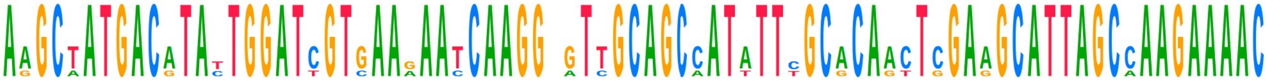

```
Consensus    AAGCTATGACATATTGGATCGTGAAGAATCAAGGAGTTGCAGCCATATTCGCACAACTCGAAGCATTAGCCAAGAAAACA
SCMV-OH      AAGCTATGACATACTGGATCGTGAAGAATCAAGGAGTTGCAGCCATATTCGCACAACTCGAAGCATTAGCCAAGAAAACG  2476
SCMV-See     AGGCAATGACGTATTGGATTGTCAAGAACCAAGGCATCGCAGCAATTTTTGCGCAGTTGGAGGCATTAGCAAAGAAAACT  2444
SCMV-Rw001   AAGCTATGACATACTGGATCGTGAAGAATCAAGGTGTTGCAGCCATATTTGCACAACTCGAAGCATTAGCCAAGAAAACA  2476
SCMV-Rw043   AAGCTATGACATATTGGATCGTGAAAAATCAAGGAGTTGCAGCCATATTCGCACAACTCGAAGCATTAGCCAAGAAAACA  2474
SCMV-Rw145   AAGCTATGACATATTGGATCGTGAAAAATCAAGGAGTTGCAGCCATATTCGCACAACTCGAAGCATTAGCCAAGAAAACA  2476
             2400  2410  2420  2430  2440  2450  2460  2470
9640 bases   60  65  70  75  80
13 features  Q A M T Y W I V K N Q G V A A I F A Q L E A L A K K T
             P3
```

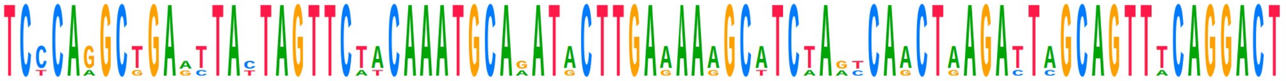

```
Consensus    TCCCAGGCTGAATTACTAGTTCTACAAATGCAGATACTTGAAAAAGCATCTAGTCAACTAAGATTAGCAGTTTCAGGACT
SCMV-OH      TCCCAGGCTGAGTTATTAGTTCTACAAATGCAAATACTTGAAAAGGCATCTAACCAACTAAGATTAGCAGTTTCAGGACT  2556
SCMV-See     TCTCAAGCGGAACTACTAGTTCTTCAAATGCAAATACTTGAGAAAGCTTCAAACCAACTGAGACTCGCAGTTACAGGACT  2524
SCMV-Rw001   TCCCAGGCTGAATTACTAGTTCTACAAATGCAGATACTTGAAAAAGCATCTAGTCAACTAAGATTAGCAGTTTCAGGACT  2556
SCMV-Rw043   TCCCAGGCTGAGTTATTAGTTCAACAAATGCAGATACTTGAAAAAGCATCTAGTCAACTAAGATTAGCAGTTTCAGGACT  2554
SCMV-Rw145   TCCCAGGCTGAATTACTAGTTCTACAAATGCAGATGCTTGAAAAAGCATCTAGTCAGCTAAGATTAGCAGTTTCAGGACT  2556
             2480  2490  2500  2510  2520  2530  2540  2550
9640 bases   85  90  95  100  105  110
13 features  S Q A E L L V L Q M Q M L E K A S S Q L R L A V S G L
             P3
```

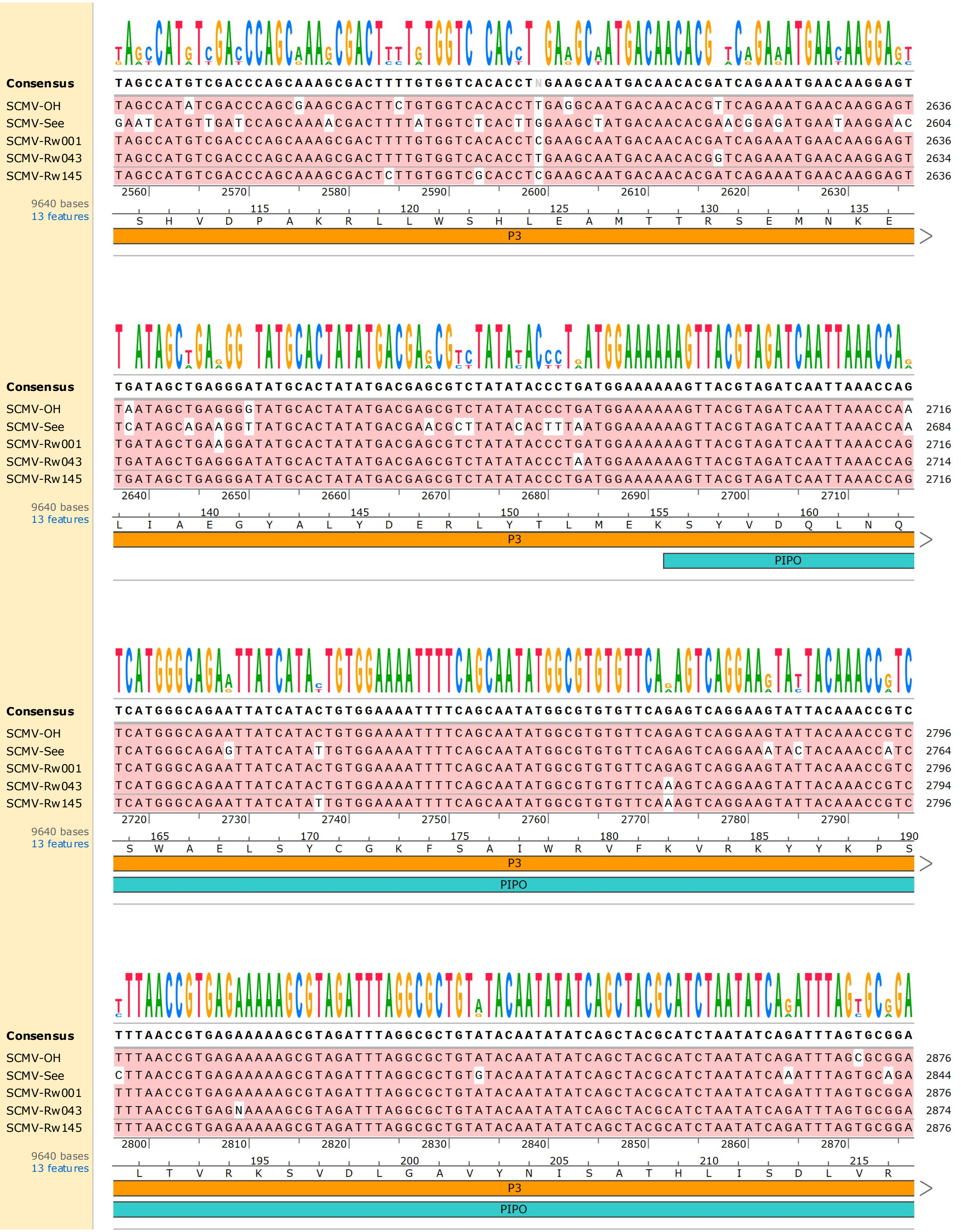
Consensus TAGCCATGTCGACCCAGCAAAGCGACTTTTGTGGTCACACCTNGAAGCAATGACAACACGATCAGAAATGAACAAGGAGT
SCMV-OH TAGCCATATCGACCCAGCGAAGCGACTTCTGTGGTCACACCTTGAGGCAATGACAACACGTTCAGAAATGAACAAGGAGT 2636
SCMV-See GAATCATGTTGATCCAGCAAAACGACTTTTATGGTCTCACTTGGAAGCTATGACAACACGAACGGAGATGAATAAGGAAC 2604
SCMV-Rw001 TAGCCATGTCGACCCAGCAAAGCGACTTTTGTGGTCACACCTCGAAGCAATGACAACACGATCAGAAATGAACAAGGAGT 2636
SCMV-Rw043 TAGCCATGTCGACCCAGCAAAGCGACTTTTGTGGTCACACCTTGAAGCAATGACAACACGGTCAGAAATGAACAAGGAGT 2634
SCMV-Rw145 TAGCCATGTCGACCCAGCAAAGCGACTCTTGTGGTCGCACCTCGAAGCAATGACAACACGATCAGAAATGAACAAGGAGT 2636
2560 2570 2580 2590 2600 2610 2620 2630
9640 bases
13 features
115 120 125 130 135
S H V D P A K R L L W S H L E A M T T R S E M N K E
P3
Consensus TGATAGCTGAGGGATATGCACTATATGACGAGCGTCTATATACCCTGATGGAAAAAAGTTACGTAGATCAATTAAACCAG
SCMV-OH TAATAGCTGAGGGGTATGCACTATATGACGAGCGTCTATATACCCTGATGGAAAAAAGTTACGTAGATCAATTAAACCAA 2716
SCMV-See TCATAGCAGAAGGTTATGCACTATATGACGAACGCTTATACACTTTAATGGAAAAAAGTTACGTAGATCAATTAAACCAA 2684
SCMV-Rw001 TGATAGCTGAAGGATATGCACTATATGACGAGCGTCTATATACCCTGATGGAAAAAAGTTACGTAGATCAATTAAACCAG 2716
SCMV-Rw043 TGATAGCTGAGGGATATGCACTATATGACGAGCGTCTATATACCCTAATGGAAAAAAGTTACGTAGATCAATTAAACCAG 2714
SCMV-Rw145 TGATAGCTGAGGGATATGCACTATATGACGAGCGTCTATATACCCTGATGGAAAAAAGTTACGTAGATCAATTAAACCAG 2716
2640 2650 2660 2670 2680 2690 2700 2710
9640 bases
13 features
140 145 150 155 160
L I A E G Y A L Y D E R L Y T L M E K S Y V D Q L N Q
P3
PIPO
Consensus TCATGGGCAGAATTATCATACTGTGGAAAATTTTCAGCAATATGGCGTGTGTTCAGAGTCAGGAAGTATTACAAACCGTC
SCMV-OH TCATGGGCAGAATTATCATACTGTGGAAAATTTTCAGCAATATGGCGTGTGTTCAGAGTCAGGAAGTATTACAAACCGTC 2796
SCMV-See TCATGGGCAGAGTTATCATATTGTGGAAAATTTTCAGCAATATGGCGTGTGTTCAGAGTCAGGAAATACTACAAACCATC 2764
SCMV-Rw001 TCATGGGCAGAATTATCATACTGTGGAAAATTTTCAGCAATATGGCGTGTGTTCAGAGTCAGGAAGTATTACAAACCGTC 2796
SCMV-Rw043 TCATGGGCAGAATTATCATACTGTGGAAAATTTTCAGCAATATGGCGTGTGTTCAAAGTCAGGAAGTATTACAAACCGTC 2794
SCMV-Rw145 TCATGGGCAGAATTATCATATTGTGGAAAATTTTCAGCAATATGGCGTGTGTTCAAAGTCAGGAAGTATTACAAACCGTC 2796
2720 2730 2740 2750 2760 2770 2780 2790
9640 bases
13 features
165 170 175 180 185 190
S W A E L S Y C G K F S A I W R V F K V R K Y Y K P S
P3
PIPO
Consensus TTTAACCGTGAGAAAAAGCGTAGATTTAGGCGCTGTATACAATATATCAGCTACGCATCTAATATCAGATTTAGTGCGGA
SCMV-OH TTTAACCGTGAGAAAAAGCGTAGATTTAGGCGCTGTATACAATATATCAGCTACGCATCTAATATCAGATTTAGCGCGGA 2876
SCMV-See CTTAACCGTGAGAAAAAGCGTAGATTTAGGCGCTGTGTACAATATATCAGCTACGCATCTAATATCAAATTTAGTGCAGA 2844
SCMV-Rw001 TTTAACCGTGAGAAAAAGCGTAGATTTAGGCGCTGTATACAATATATCAGCTACGCATCTAATATCAGATTTAGTGCGGA 2876
SCMV-Rw043 TTTAACCGTGAGNAAAAGCGTAGATTTAGGCGCTGTATACAATATATCAGCTACGCATCTAATATCAGATTTAGTGCGGA 2874
SCMV-Rw145 TTTAACCGTGAGAAAAAGCGTAGATTTAGGCGCTGTATACAATATATCAGCTACGCATCTAATATCAGATTTAGTGCGGA 2876
2800 2810 2820 2830 2840 2850 2860 2870
9640 bases
13 features
195 200 205 210 215
L T V R K S V D L G A V Y N I S A T H L I S D L V R
P3
PIPO

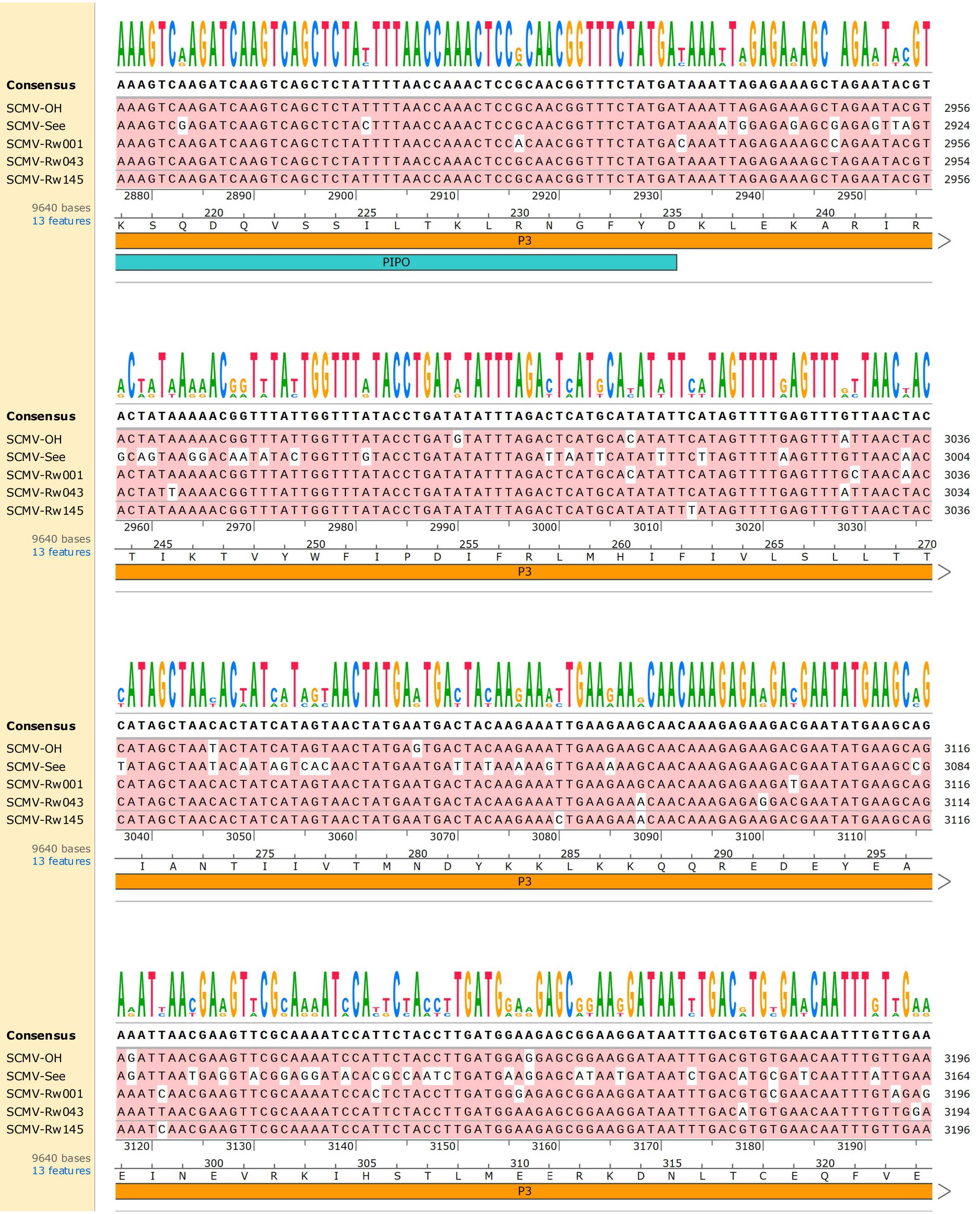
Consensus
SCMV-OH
SCMV-See
SCMV-Rw001
SCMV-Rw043
SCMV-Rw145
9640 bases
13 features
P3
PIPO

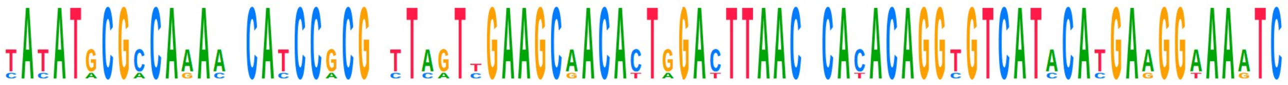

```
Consensus     TATATGCGCCAAAANCATCCGCGGTTAGTTGAAGCAACACTGGACTTAACCCACACAGGTGTCATACATGAAGGAAAATC
SCMV-OH       TATATGCGCCAAAATCATCCGCGGTTAGTTGAAGCGACACTGGACTTAACTCACACAGGCGTCATACACGAAGGAAAATC  3276
SCMV-See      CACATACGACAGACACATCCACGCCTCATCGAAGCAACATTGGATTTAACACATACAGGTGTCATCCATGAGGGTAAGTC  3244
SCMV-Rw001    TATATGCGCCAAAATCACCCGCGATTAGTTGAAGCAACACTGGACTTAACCCACACAGGTGTCATACATGAAGGAAAATC  3276
SCMV-Rw043    TATATGCGCCAAAACCATCCGCGGTTAGTTGAAGCAACACTAGACTTAACCCACACAGGTGTCATACATGAAGGAAAATC  3274
SCMV-Rw145    TATATGCGCCAAAACCATCCGCGGTTAGTCGAAGCAACACTGGACTTAACCCACACAGGTGTCATACATGAAGGAAAATC  3276
              3200      3210      3220      3230      3240      3250      3260      3270
9640 bases     325        330        335        340        345     1
13 features    Y  M  R  Q  N  H  P  R  L  V  E  A  T  L  D  L  T  H  T  G  V  I  H  E  G  K  S
               P3                                                                      6K1
```

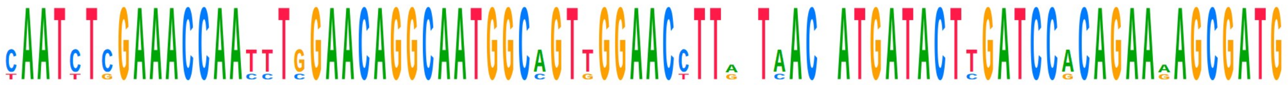

```
Consensus     CAATCTCGAAACCAATTTGGAACAGGCAATGGCAGTTGGAACCTTAATAACCATGATACTTGATCCACAGAAAAGCGATG
SCMV-OH       TAATCTCGAAACCAATTTGGAACAGGCAATGGCAGTTGGAACCTTGATAACAATGATACTTGATCCACAGAAAAGCGATG  3356
SCMV-See      CAATTTGGAAACCAACCTCGAACAGGCAATGGCCGTGGGAACTTTACTCACTATGATACTCGATCCACAGAAGAGCGATG  3324
SCMV-Rw001    CAATCTCGAAACCAATTTGGAACAGGCAATGGCAGTTGGAACCTTATTAACCATGATACTTGATCCGCAGAAGAGCGATG  3356
SCMV-Rw043    CAATCTCGAAACCAATTTGGAACAGGCAATGGCAGTTGGAACCTTGATAACCATGATACTTGATCCACAGAAAAGCGATG  3354
SCMV-Rw145    CAATCTCGAAACCAATTTGGAACAGGCAATGGCAGTTGGAACCTTAATAACCATGATACTTGATCCACAGAAAAGCGATG  3356
              3280      3290      3300      3310      3320      3330      3340      3350
9640 bases       5          10         15         20         25
13 features    N  L  E  T  N  L  E  Q  A  M  A  V  G  T  L  I  T  M  I  L  D  P  Q  K  S  D
               6K1
```

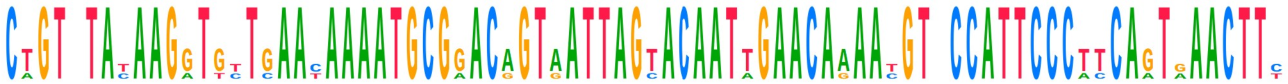

```
Consensus     CTGTCTATAAGGTGTTGAACAAAATGCGGACAGTAATTAGTACAATTGAACAAAATGTCCCATTCCCTTCAGTGAACTTC
SCMV-OH       CTGTCTATAAGGTGTTGAACAAAATGCGGACAGTAATTAGTACAATTGAACAAAATGTCCCATTCCCTTCAGTGAACTTC  3436
SCMV-See      CAGTTTATAAGATTCTCAATAAAATGCGAACGGTGATTAGCACAATAGAACAAAATGTGCCATTCCCATCAGTGAACTTC  3404
SCMV-Rw001    CTGTATATAAGGTGTTGAACAAAATGCGGACAGTAATTAGTACAATTGAACAAAACGTCCCATTCCCTTCAATAAACTTT  3436
SCMV-Rw043    CTGTCTACAAGGTGTTGAACAAAATGCGGACAGTAATTAGTACAATTGAACAAAACGTCCCATTCCCTTCAGTAAACTTT  3434
SCMV-Rw145    CTGTCTACAAGGTGCTGAATAAAATGCGGACAGTAATTAGTACAATTGAACAGAATGTTCCATTCCCTCCAGTGAACTTC  3436
              3360      3370      3380      3390      3400      3410      3420      3430
9640 bases    30        35         40         45         50         55
13 features    A  V  Y  K  V  L  N  K  M  R  T  V  I  S  T  I  E  Q  N  V  P  F  P  P  V  N  F
               6K1
```

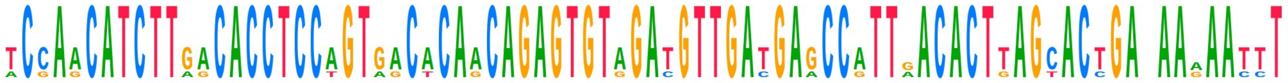

```
Consensus     TCCAACATCTTGACACCTCCAGTGACACAACAGAGTGTAGATGTTGATGAGCCATTGACACTTAGCACTGANAAAAATTT
SCMV-OH       TCCAACATCTTGACACCTCCAGTGACACAACAGAGTGTAGATGTTGATGAGCCATTGACACTTAGCACTGATAAAAACCT  3516
SCMV-See      ACGAGCATCTTGGCACCTCCTGTAGCTCAGCAGAGTGTAGATGTTGACGAACCGTTAACACTGAGTACCGACAAGAATTT  3484
SCMV-Rw001    TCCAACATCTTGACACCTCCAGTGACACAACAGAGTGTAGATGTTGATGAGCCATTGACACTTAGCACTGAGAAGAATTT  3516
SCMV-Rw043    TCCAACATCTTGACACCTCCAGTGACACAACAGAGTGTAGACGTTGATGAGCCATTGACACTTAGCACTGAGAAAAATTT  3514
SCMV-Rw145    TCCAACATCTTAACACCTCCAGTGACACAACAGAGTGTGGATGTTGATGAGCCATTAACACTTAGCACTGATAAAAATTT  3516
              3440      3450      3460      3470      3480      3490      3500      3510
9640 bases          60         65       1         5          10         15
13 features    S  N  I  L  T  P  P  V  T  Q  Q  S  V  D  V  D  E  P  L  T  L  S  T  D  K  N  L
               6K1                            CIP
```

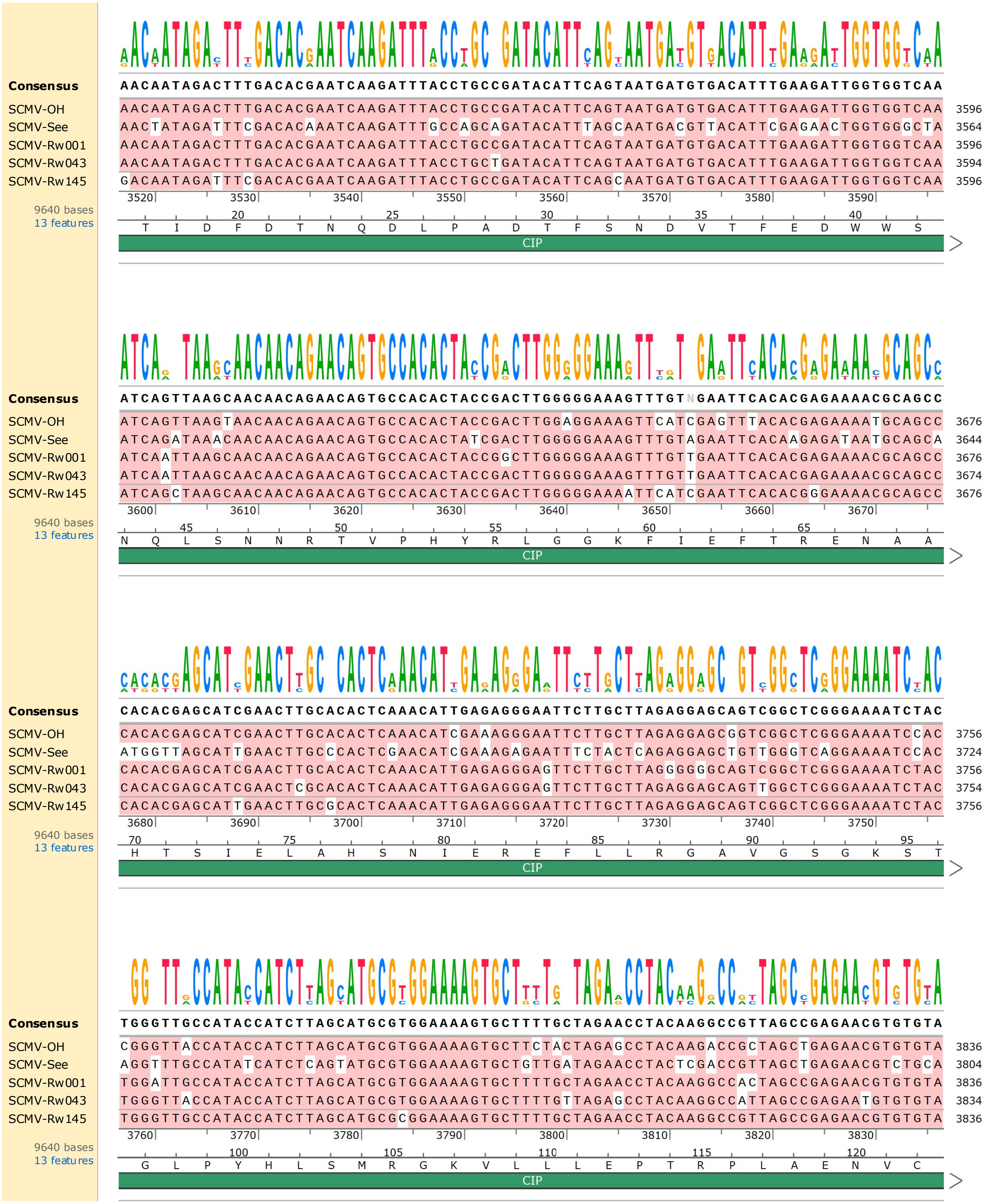

Consensus AACAATAGACTTTGACACGAATCAAGATTTACCTGCCGATACATTCAGTAATGATGTGACATTTGAAGATTGGTGGTCAA
SCMV-OH AACAATAGACTTTGACACGAATCAAGATTTACCTGCCGATACATTCAGTAATGATGTGACATTTGAAGATTGGTGGTCAA 3596
SCMV-See AACTATAGATTTCGACACAAATCAAGATTTGCCAGCAGATACATTTAGCAATGACGTTACATTCGAGAACTGGTGGGCTA 3564
SCMV-Rw001 AACAATAGACTTTGACACGAATCAAGATTTACCTGCCGATACATTCAGTAATGATGTGACATTTGAAGATTGGTGGTCAA 3596
SCMV-Rw043 AACAATAGACTTTGACACGAATCAAGATTTACCTGCTGATACATTCAGTAATGATGTGACATTTGAAGATTGGTGGTCAA 3594
SCMV-Rw145 GACAATAGATTTCGACACGAATCAAGATTTACCTGCCGATACATTCAGCAATGATGTGACATTTGAAGATTGGTGGTCAA 3596
3520 3530 3540 3550 3560 3570 3580 3590
9640 bases
13 features
20 25 30 35 40
T I D F D T N Q D L P A D T F S N D V T F E D W W S
CIP
Consensus ATCAGTTAAGCAACAACAGAACAGTGCCACACTACCGACTTGGGGGAAAGTTTGTNGAATTCACACGAGAAAACGCAGCC
SCMV-OH ATCAGTTAAGTAACAACAGAACAGTGCCACACTACCGACTTGGAGGAAAGTTCATCGAGTTTACACGAGAAAATGCAGCC 3676
SCMV-See ATCAGATAAACAACAACAGAACAGTGCCACACTATCGACTTGGGGGAAAGTTTGTAGAATTCACAAGAGATAATGCAGCA 3644
SCMV-Rw001 ATCAATTAAGCAACAACAGAACAGTGCCACACTACCGGCTTGGGGGAAAGTTTGTTGAATTCACACGAGAAAACGCAGCC 3676
SCMV-Rw043 ATCAATTAAGCAACAACAGAACAGTGCCACACTACCGACTTGGGGGAAAGTTTGTTGAATTCACACGAGAAAACGCAGCC 3674
SCMV-Rw145 ATCAGCTAAGCAACAACAGAACAGTGCCACACTACCGACTTGGGGGAAAATTCATCGAATTCACACGGGAAAACGCAGCC 3676
3600 3610 3620 3630 3640 3650 3660 3670
9640 bases
13 features
45 50 55 60 65
N Q L S N N R T V P H Y R L G G K F I E F T R E N A A
CIP
Consensus CACACGAGCATCGAACTTGCACACTCAAACATTGAGAGGGAATTCTTGCTTAGAGGAGCAGTCGGCTCGGGAAAATCTAC
SCMV-OH CACACGAGCATCGAACTTGCACACTCAAACATCGAAAGGGAATTCTTGCTTAGAGGAGCGGTCGGCTCGGGAAAATCCAC 3756
SCMV-See ATGGTTAGCATTGAACTTGCCCACTCGAACATCGAAAGAGAATTTCTACTCAGAGGAGCTGTTGGGTCAGGAAAATCCAC 3724
SCMV-Rw001 CACACGAGCATCGAACTTGCACACTCAAACATTGAGAGGGAGTTCTTGCTTAGGGGGCAGTCGGCTCGGGAAAATCTAC 3756
SCMV-Rw043 CACACGAGCATCGAACTCGCACACTCAAACATTGAGAGGGAGTTCTTGCTTAGAGGAGCAGTTGGCTCGGGAAAATCTAC 3754
SCMV-Rw145 CACACGAGCATTGAACTTGCGCACTCAAACATTGAGAGGGAATTCTTGCTTAGAGGAGCAGTCGGCTCGGGAAAATCTAC 3756
3680 3690 3700 3710 3720 3730 3740 3750
9640 bases
13 features
70 75 80 85 90 95
H T S I E L A H S N I E R E F L L R G A V G S G K S T
CIP
Consensus TGGGTTGCCATACCATCTTAGCATGCGTGGAAAAGTGCTTTTGCTAGAACCTACAAGGCCGTTAGCCGAGAACGTGTGTA
SCMV-OH CGGGTTACCATACCATCTTAGCATGCGTGGAAAAGTGCTTCTACTAGAGCCTACAAGACCGCTAGCTGAGAACGTGTGTA 3836
SCMV-See AGGTTTGCCATATCATCTCAGTATGCGTGGAAAAGTGCTGTTGATAGAACCTACTCGACCGTTAGCTGAGAACGTCTGCA 3804
SCMV-Rw001 TGGATTGCCATACCATCTTAGCATGCGTGGAAAAGTGCTTTTGCTAGAACCTACAAGGCCACTAGCCGAGAACGTGTGTA 3836
SCMV-Rw043 TGGGTTACCATACCATCTTAGCATGCGTGGAAAAGTGCTTTTGTTAGAGCCTACAAGGCCATTAGCCGAGAATGTGTGTA 3834
SCMV-Rw145 TGGGTTGCCATACCATCTTAGCATGCGCGGAAAAGTGCTTTTGCTAGAACCTACAAGGCCGTTAGCCGAGAACGTGTGTA 3836
3760 3770 3780 3790 3800 3810 3820 3830
9640 bases
13 features
100 105 110 115 120
G L P Y H L S M R G K V L L L E P T R P L A E N V C
CIP

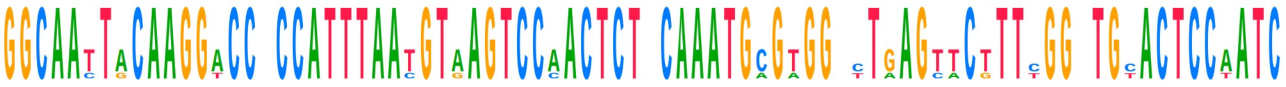

| Sequence | Alignment | End |
| --- | --- | --- |
| Consensus | GGCAATTACAAGGACCACCATTTAATGTAAGTCCAACTCTCCAAATGCGTGGACTGAGTTCTTTCGGGTGCACTCCAATC | |
| SCMV-OH | GGCAATTACAAGGACCGCCATTTAACGTAAGTCCAACTCTTCAAATGCGTGGACTAAGTTCTTTTGGATGCACTCCAATC | 3916 |
| SCMV-See | GGCAACTGCAAGGTCCTCCATTTAATGTGAGTCCCACTCTACAAATGAGAGGTTTGAGCACGTTTGGCTGCACTCCTATC | 3884 |
| SCMV-Rw001 | GGCAATTACAAGGACCACCATTTAATGTAAGTCCAACTCTCCAAATGCGTGGGCTGAGTTCTTTCGGGTGTACTCCAATC | 3916 |
| SCMV-Rw043 | GGCAATTACAAGGACCACCATTTAATGTAAGTCCAACTCTCCAAATGCGTGGATTGAGTTCTTTCGGGTGCACTCCAATC | 3914 |
| SCMV-Rw145 | GGCAATTACAAGGACCACCATTTAATGTAAGTCCAACTCTCCAAATGCGTGGACTGAGTTCTTTCGGGTGTACTCCAATC | 3916 |

3840 3850 3860 3870 3880 3890 3900 3910

9640 bases
13 features

125 130 135 140 145
R Q L Q G P P F N V S P T L Q M R G L S S F G C T P I
CIP

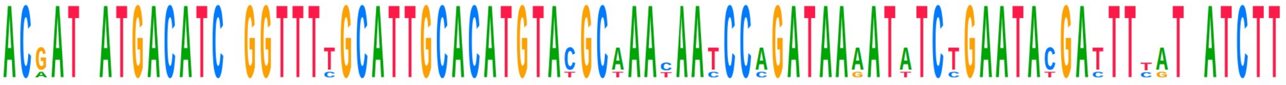

| Sequence | Alignment | End |
| --- | --- | --- |
| Consensus | ACGATCATGACATCAGGTTTTGCATTGCACATGTACGCAAACAATCCAGATAAAATATCTGAATACGATTTTATAATCTT | |
| SCMV-OH | ACAATCATGACATCCGGTTTTGCATTGCACATGTACGCAAACAATCCAGATAAAATATCCGAATACGATTTTATTATCTT | 3996 |
| SCMV-See | ACGATAATGACATCTGGTTTCGCATTGCACATGTATGCTAATAACCCCGATAAGATTTCTGAATACGACTTCATCATCTT | 3964 |
| SCMV-Rw001 | ACGATCATGACATCAGGTTTTGCATTGCACATGTACGCAAACAATCCAGATAAAATATCTGAATATGATTTTGTAATCTT | 3996 |
| SCMV-Rw043 | ACGATCATGACATCAGGTTTTGCATTGCACATGTACGCAAATAATCCAGATAAAATATCTGAATACGATTTTATAATCTT | 3994 |
| SCMV-Rw145 | ACGATTATGACATCAGGTTTTGCATTGCACATGTACGCAAACAATCCAGATAAAATATCTGAATACGATTTCATAATCTT | 3996 |

3920 3930 3940 3950 3960 3970 3980 3990

9640 bases
13 features

150 155 160 165 170 175
T I M T S G F A L H M Y A N N P D K I S E Y D F I I F
CIP

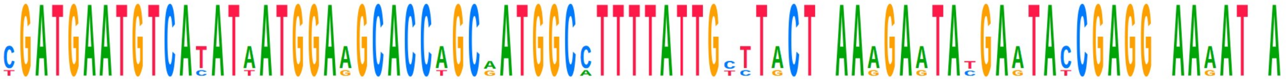

| Sequence | Alignment | End |
| --- | --- | --- |
| Consensus | CGATGAATGTCATATAATGGAAGCACCAGCAATGGCCTTTTATTGCTTACTCAAAGAATATGAATACCGAGGGAAAATCA | |
| SCMV-OH | CGATGAATGTCATATAATGGAAGCACCAGCGATGGCCTTTTATTGCTTACTCAAAGAATATGAGTACCGAGGGAAAATTA | 4076 |
| SCMV-See | TGATGAATGTCACATTATGGAAGCACCTGCGATGGCATTTTATTGTTTGCTTAAGGAGTATGAATACCGAGGCAAGATAA | 4044 |
| SCMV-Rw001 | CGATGAATGTCATATAATGGAGGCACCAGCAATGGCCTTTTATTGTCTACTCAAAGAATACGAATACCGAGGAAAAATCA | 4076 |
| SCMV-Rw043 | CGATGAATGTCATATAATGGAAGCACCAGCAATGGCCTTTTATTGCTTACTGAAAGAATACGAATACCGAGGGAAAATCA | 4074 |
| SCMV-Rw145 | CGATGAATGTCATATAATGGAAGCACCAGCAATGGCCTTTTATTGCTTACTCAAAGAATATGAATATCGAGGGAAAATCA | 4076 |

4000 4010 4020 4030 4040 4050 4060 4070

9640 bases
13 features

180 185 190 195 200
D E C H I M E A P A M A F Y C L L K E Y E Y R G K I
CIP

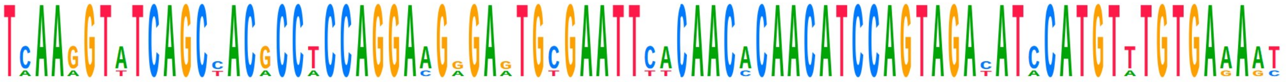

| Sequence | Alignment | End |
| --- | --- | --- |
| Consensus | TCAAGGTATCAGCCACGCCTCCAGGAAGGGAGTGCGAATTCACAACACAACATCCAGTAGACATCCATGTTTGTGAAAAT | |
| SCMV-OH | TCAAGGTATCAGCTACGCCTCCAGGAAGAGAGTGTGAATTCACAACACAACATCCAGTAGACATCCATGTTTGTGAGAAT | 4156 |
| SCMV-See | TAAAAGTTTCAGCTACACCACCAGGACGAGAATGCGAATTTTCAACCCAACATCCAGTAGATATACATGTATGTGAAAGC | 4124 |
| SCMV-Rw001 | TCAAGGTATCAGCCACGCCTCCAGGAAGGGAGTGCGAATTCACAACACAACATCCAGTAGATATCCATGTTTGTGAAAAT | 4156 |
| SCMV-Rw043 | TCAAGGTATCAGCCACGCCTCCAGGAAGGGAATGCGAATTCACAACACAACATCCAGTAGACATCCATGTTTGTGAAAAT | 4154 |
| SCMV-Rw145 | TCAAGGTATCAGCCACGCCTCCAGGAAGGGAGTGCGAATTCACAACACAACATCCAGTAGACATCCATGTTTGTGAAAAT | 4156 |

4080 4090 4100 4110 4120 4130 4140 4150

9640 bases
13 features

205 210 215 220 225
I K V S A T P P G R E C E F T T Q H P V D I H V C E N
CIP

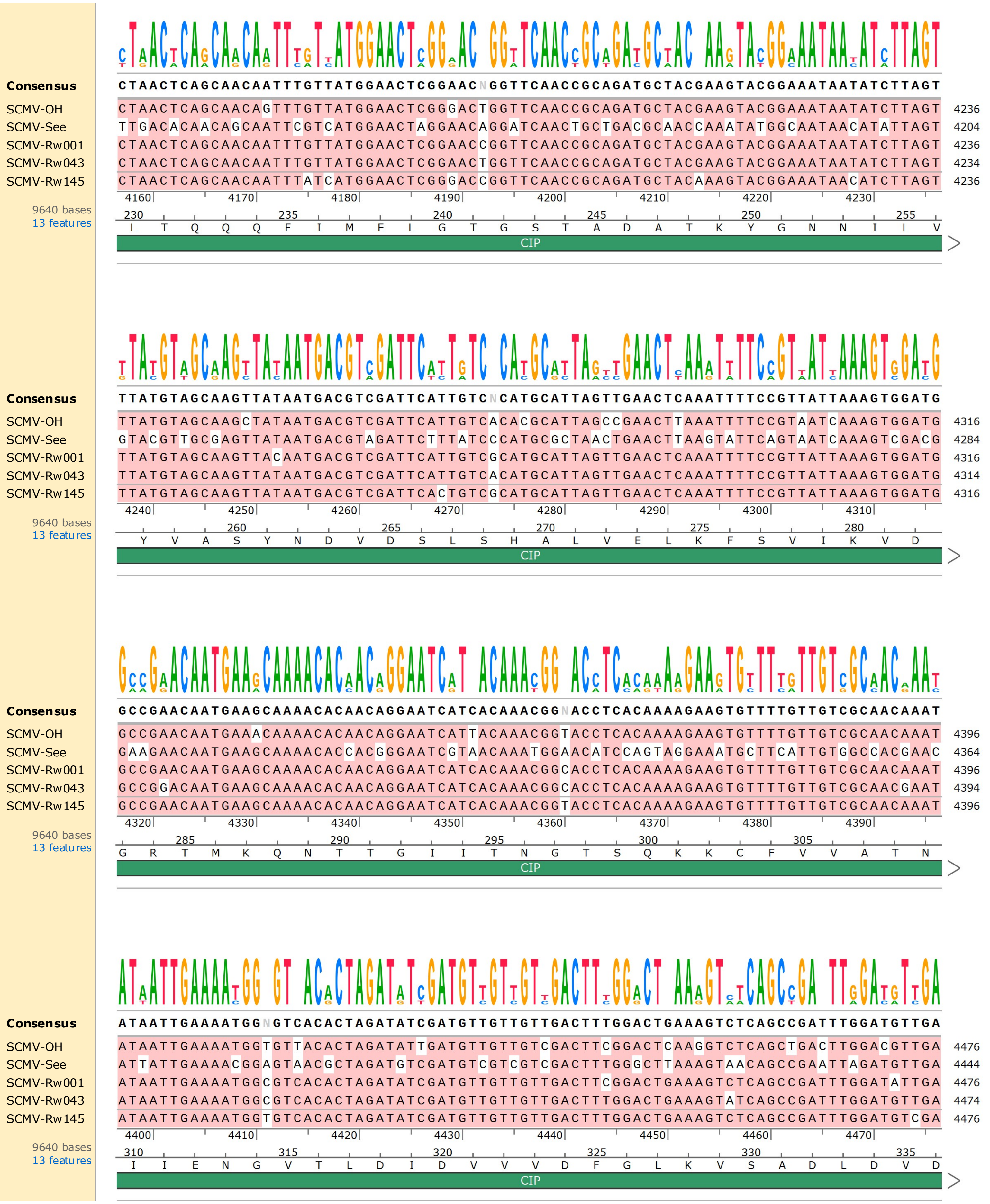

Consensus
CTAACTCAGCAACAATTTGTTATGGAACTCGGAACNGGTTCAACCGCAGATGCTACGAAGTACGGAAATAATATCTTAGT
SCMV-OH CTAACTCAGCAACAGTTTGTTATGGAACTCGGGACTGGTTCAACCGCAGATGCTACGAAGTACGGAAATAATATCTTAGT 4236
SCMV-See TTGACACAACAGCAATTCGTCATGGAACTAGGAACAGGATCAACTGCTGACGCAACCAAATATGGCAATAACATATTAGT 4204
SCMV-Rw001 CTAACTCAGCAACAATTTGTTATGGAACTCGGAACCGGTTCAACCGCAGATGCTACGAAGTACGGAAATAATATCTTAGT 4236
SCMV-Rw043 CTAACTCAGCAACAATTTGTTATGGAACTCGGAACTGGTTCAACCGCAGATGCTACGAAGTACGGAAATAATATCTTAGT 4234
SCMV-Rw145 CTAACTCAGCAACAATTTATCATGGAACTCGGGACCGGTTCAACCGCAGATGCTACAAAGTACGGAAATAACATCTTAGT 4236
4160 4170 4180 4190 4200 4210 4220 4230
9640 bases
13 features
230 235 240 245 250 255
L T Q Q Q F I M E L G T G S T A D A T K Y G N N I L V
CIP
Consensus
TTATGTAGCAAGTTATAATGACGTCGATTCATTGTCNCATGCATTAGTTGAACTCAAATTTTCCGTTATTAAAGTGGATG
SCMV-OH TTATGTAGCAAGCTATAATGACGTCGATTCATTGTCACACGCATTAGCCGAACTTAAATTTTCCGTAATCAAAGTGGATG 4316
SCMV-See GTACGTTGCGAGTTATAATGACGTAGATTCTTTATCCCATGCGCTAACTGAACTTAAGTATTCAGTAATCAAAGTCGACG 4284
SCMV-Rw001 TTATGTAGCAAGTTACAATGACGTCGATTCATTGTCGCATGCATTAGTTGAACTCAAATTTTCCGTTATTAAAGTGGATG 4316
SCMV-Rw043 TTATGTAGCAAGTTATAATGACGTCGATTCATTGTCACATGCATTAGTTGAACTCAAATTTTCCGTTATTAAAGTGGATG 4314
SCMV-Rw145 TTATGTAGCAAGTTATAATGACGTCGATTCACTGTCGCATGCATTAGTTGAACTCAAATTTTCCGTTATTAAAGTGGATG 4316
4240 4250 4260 4270 4280 4290 4300 4310
9640 bases
13 features
260 265 270 275 280
Y V A S Y N D V D S L S H A L V E L K F S V I K V D
CIP
Consensus
GCCGAACAATGAAGCAAAACACAACAGGAATCATCACAAACGGNACCTCACAAAAGAAGTGTTTTGTTGTCGCAACAAAT
SCMV-OH GCCGAACAATGAAACAAAACACAACAGGAATCATTACAAACGGTACCTCACAAAAGAAGTGTTTTGTTGTCGCAACAAAT 4396
SCMV-See GAAGAACAATGAAGCAAAACACCACGGGAATCGTAACAAATGGAACATCCAGTAGGAAATGCTTCATTGTGGCCACGAAC 4364
SCMV-Rw001 GCCGAACAATGAAGCAAAACACAACAGGAATCATCACAAACGGCACCTCACAAAAGAAGTGTTTTGTTGTCGCAACAAAT 4396
SCMV-Rw043 GCCGGACAATGAAGCAAAACACAACAGGAATCATCACAAACGGCACCTCACAAAAGAAGTGTTTTGTTGTCGCAACGAAT 4394
SCMV-Rw145 GCCGAACAATGAAGCAAAACACAACAGGAATCATCACAAACGGTACCTCACAAAAGAAGTGTTTTGTTGTCGCAACAAAT 4396
4320 4330 4340 4350 4360 4370 4380 4390
9640 bases
13 features
285 290 295 300 305
G R T M K Q N T T G I I T N G T S Q K K C F V V A T N
CIP
Consensus
ATAATTGAAAATGGNGTCACACTAGATATCGATGTTGTTGTTGACTTTGGACTGAAAGTCTCAGCCGATTTGGATGTTGA
SCMV-OH ATAATTGAAAATGGTGTTACACTAGATATTGATGTTGTTGTCGACTTCGGACTCAAGGTCTCAGCTGACTTGGACGTTGA 4476
SCMV-See ATTATTGAAAACGGAGTAACGCTAGATGTCGATGTCGTCGTCGACTTTGGGCTTAAAGTAACAGCCGAATTAGATGTTGA 4444
SCMV-Rw001 ATAATTGAAAATGGCGTCACACTAGATATCGATGTTGTTGTTGACTTCGGACTGAAAGTCTCAGCCGATTTGGATATTGA 4476
SCMV-Rw043 ATAATTGAAAATGGCGTCACACTAGATATCGATGTTGTTGTTGACTTTGGACTGAAAGTATCAGCCGATTTGGATGTTGA 4474
SCMV-Rw145 ATAATTGAAAATGGTGTCACACTAGATATCGATGTTGTTGTTGACTTTGGACTGAAAGTCTCAGCCGATTTGGATGTCGA 4476
4400 4410 4420 4430 4440 4450 4460 4470
9640 bases
13 features
310 315 320 325 330 335
I I E N G V T L D I D V V V D F G L K V S A D L D V D
CIP

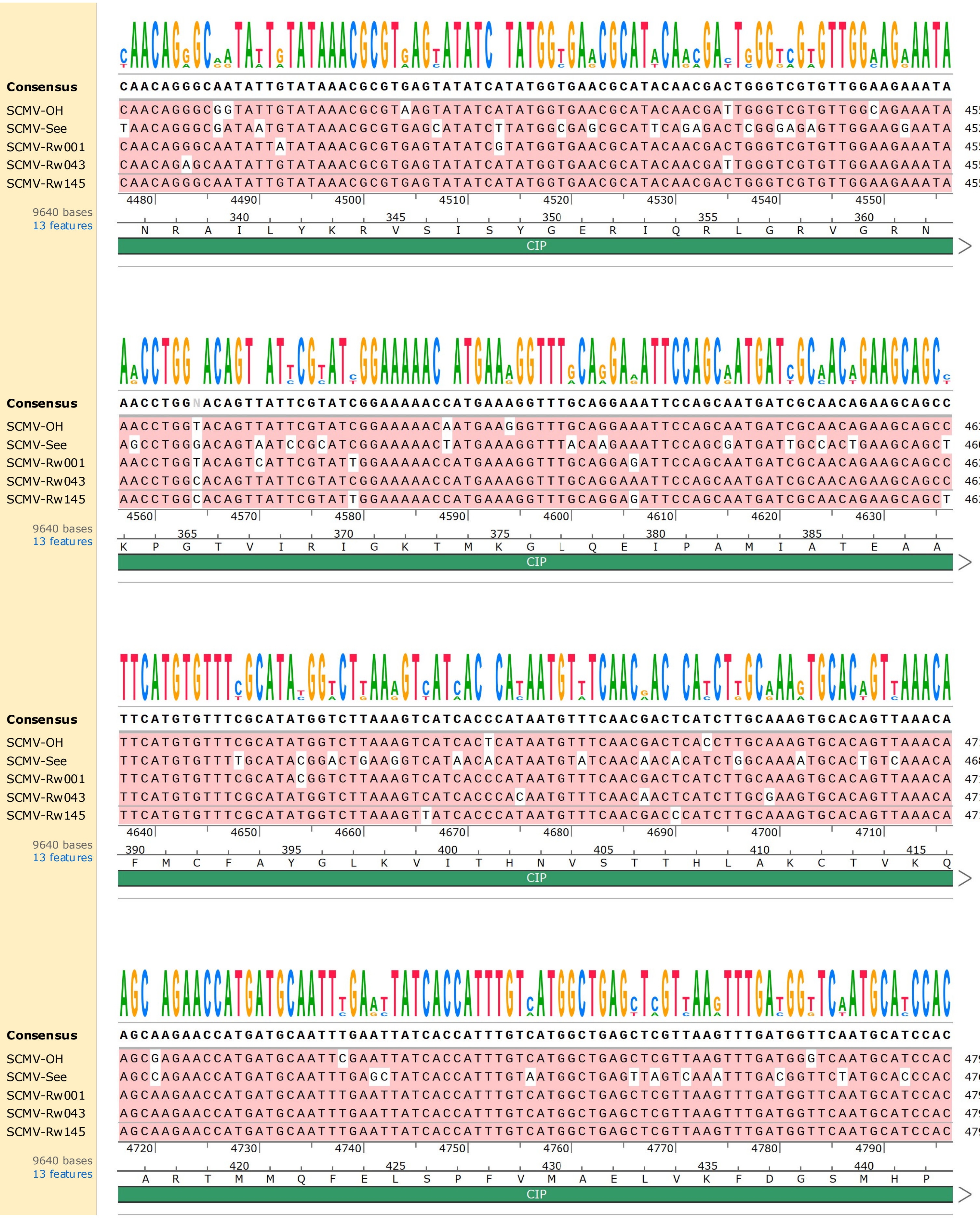

Consensus CAACAGGGCAATATTGTATAAACGCGTGAGTATATCATATGGTGAACGCATACAACGACTGGGTCGTGTTGGAAGAAATA
SCMV-OH CAACAGGGCGGTATTGTATAAACGCGTAAGTATATCATATGGTGAACGCATACAACGATTGGGTCGTGTTGGCAGAAATA 4556
SCMV-See TAACAGGGCGATAATGTATAAACGCGTGAGCATATCTTATGGCGAGCGCATTCAGAGACTCGGGAGAGTTGGAAGGAATA 4524
SCMV-Rw001 CAACAGGGCAATATTATATAAACGCGTGAGTATATCGTATGGTGAACGCATACAACGACTGGGTCGTGTTGGAAGAAATA 4556
SCMV-Rw043 CAACAGAGCAATATTGTATAAACGCGTGAGTATATCATATGGTGAACGCATACAACGATTGGGTCGTGTTGGAAGAAATA 4554
SCMV-Rw145 CAACAGGGCAATATTGTATAAACGCGTGAGTATATCATATGGTGAACGCATACAACGACTGGGTCGTGTTGGAAGAAATA 4556
4480 4490 4500 4510 4520 4530 4540 4550
9640 bases
13 features
340 345 350 355 360
N R A I L Y K R V S I S Y G E R I Q R L G R V G R N
CIP
Consensus AACCTGGNACAGTTATTCGTATCGGAAAAACCATGAAAGGTTTGCAGGAAATTCCAGCAATGATCGCAACAGAAGCAGCC
SCMV-OH AACCTGGTACAGTTATTCGTATCGGAAAAACAATGAAGGGTTTGCAGGAAATTCCAGCAATGATCGCAACAGAAGCAGCC 4636
SCMV-See AGCCTGGGACAGTAATCCGCATCGGAAAAACTATGAAAGGTTTACAAGAAATTCCAGCGATGATTGCCACTGAAGCAGCT 4604
SCMV-Rw001 AACCTGGTACAGTCATTCGTATTGGAAAAACCATGAAAGGTTTGCAGGAGATTCCAGCAATGATCGCAACAGAAGCAGCC 4636
SCMV-Rw043 AACCTGGCACAGTTATTCGTATCGGAAAAACCATGAAAGGTTTGCAGGAAATTCCAGCAATGATCGCAACAGAAGCAGCC 4634
SCMV-Rw145 AACCTGGCACAGTTATTCGTATTGGAAAAACCATGAAAGGTTTGCAGGAGATTCCAGCAATGATCGCAACAGAAGCAGCT 4636
4560 4570 4580 4590 4600 4610 4620 4630
9640 bases
13 features
365 370 375 380 385
K P G T V I R I G K T M K G L Q E I P A M I A T E A A
CIP
Consensus TTCATGTGTTTCGCATATGGTCTTAAAGTCATCACCCATAATGTTTCAACGACTCATCTTGCAAAGTGCACAGTTAAACA
SCMV-OH TTCATGTGTTTCGCATATGGTCTTAAAGTCATCACTCATAATGTTTCAACGACTCACCTTGCAAAGTGCACAGTTAAACA 4716
SCMV-See TTCATGTGTTTTGCATACGGACTGAAGGTCATAACACATAATGTATCAACAACACATCTGGCAAAATGCACTGTCAAACA 4684
SCMV-Rw001 TTCATGTGTTTCGCATACGGTCTTAAAGTCATCACCCATAATGTTTCAACGACTCATCTTGCAAAGTGCACAGTTAAACA 4716
SCMV-Rw043 TTCATGTGTTTCGCATATGGTCTTAAAGTCATCACCCACAATGTTTCAACAACTCATCTTGCGAAGTGCACAGTTAAACA 4714
SCMV-Rw145 TTCATGTGTTTCGCATATGGTCTTAAAGTTATCACCCATAATGTTTCAACGACCCATCTTGCAAAGTGCACAGTTAAACA 4716
4640 4650 4660 4670 4680 4690 4700 4710
9640 bases
13 features
390 395 400 405 410 415
F M C F A Y G L K V I T H N V S T T H L A K C T V K Q
CIP
Consensus AGCAAGAACCATGATGCAATTTGAATTATCACCATTTGTCATGGCTGAGCTCGTTAAGTTTGATGGTTCAATGCATCCAC
SCMV-OH AGCGAGAACCATGATGCAATTCGAATTATCACCATTTGTCATGGCTGAGCTCGTTAAGTTTGATGGGTCAATGCATCCAC 4796
SCMV-See AGCCAGAACCATGATGCAATTTGAGCTATCACCATTTGTAATGGCTGAGTTAGTCAAATTTGACGGTTCTATGCACCCAC 4764
SCMV-Rw001 AGCAAGAACCATGATGCAATTTGAATTATCACCATTTGTCATGGCTGAGCTCGTTAAGTTTGATGGTTCAATGCATCCAC 4796
SCMV-Rw043 AGCAAGAACCATGATGCAATTTGAATTATCACCATTTGTCATGGCTGAGCTCGTTAAGTTTGATGGTTCAATGCATCCAC 4794
SCMV-Rw145 AGCAAGAACCATGATGCAATTTGAATTATCACCATTTGTCATGGCTGAGCTCGTTAAGTTTGATGGTTCAATGCATCCAC 4796
4720 4730 4740 4750 4760 4770 4780 4790
9640 bases
13 features
420 425 430 435 440
A R T M M Q F E L S P F V M A E L V K F D G S M H P
CIP

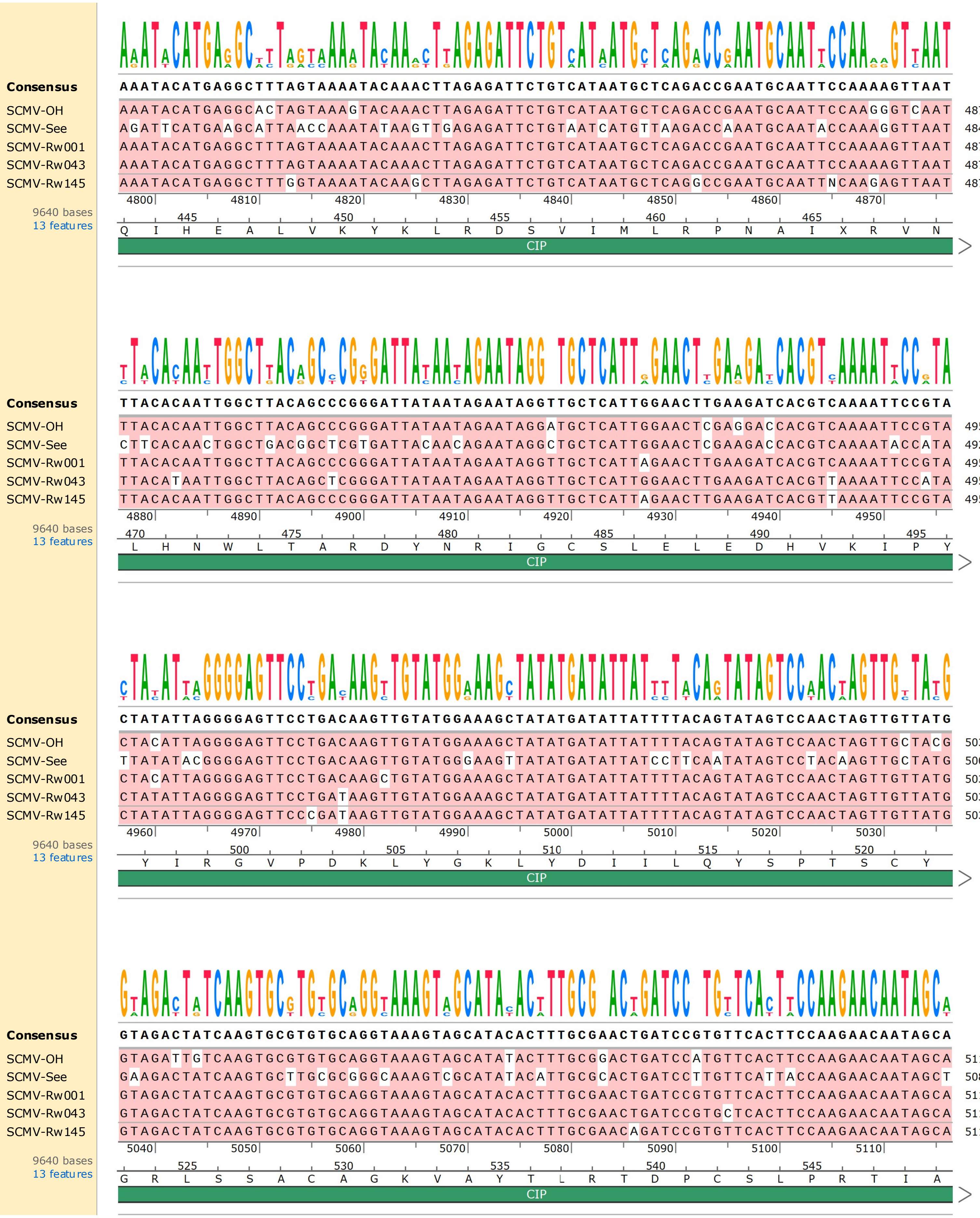
Consensus AAATACATGAGGCTTTAGTAAAATACAAACTTAGAGATTCTGTCATAATGCTCAGACCGAATGCAATTCCAAAAGTTAAT
SCMV-OH AAATACATGAGGCACTAGTAAAGTACAAACTTAGAGATTCTGTCATAATGCTCAGACCGAATGCAATTCCAAGGGTCAAT 4876
SCMV-See AGATTCATGAAGCATTAACCAAATATAAGTTGAGAGATTCTGTAATCATGTTAAGACCAAATGCAATACCAAAGGTTAAT 4844
SCMV-Rw001 AAATACATGAGGCTTTAGTAAAATACAAACTTAGAGATTCTGTCATAATGCTCAGACCGAATGCAATTCCAAAAGTTAAT 4876
SCMV-Rw043 AAATACATGAGGCTTTAGTAAAATACAAACTTAGAGATTCTGTCATAATGCTCAGACCGAATGCAATTCCAAAAGTTAAT 4874
SCMV-Rw145 AAATACATGAGGCTTTGGTAAAATACAAGCTTAGAGATTCTGTCATAATGCTCAGGCCGAATGCAATTNCAAGAGTTAAT 4876
4800 4810 4820 4830 4840 4850 4860 4870
9640 bases
13 features
445 450 455 460 465
Q I H E A L V K Y K L R D S V I M L R P N A I X R V N
CIP
Consensus TTACACAATTGGCTTACAGCCCGGGATTATAATAGAATAGGTTGCTCATTGGAACTTGAAGATCACGTCAAAATTCCGTA
SCMV-OH TTACACAATTGGCTTACAGCCCGGGATTATAATAGAATAGGATGCTCATTGGAACTCGAGGACCACGTCAAAATTCCGTA 4956
SCMV-See CTTCACAACTGGCTGACGGCTCGTGATTACAACAGAATAGGCTGCTCATTGGAACTCGAAGACCACGTCAAAATACCATA 4924
SCMV-Rw001 TTACACAATTGGCTTACAGCCCGGGATTATAATAGAATAGGTTGCTCATTAGAACTTGAAGATCACGTCAAAATTCCGTA 4956
SCMV-Rw043 TTACATAATTGGCTTACAGCTCGGGATTATAATAGAATAGGTTGCTCATTGGAACTTGAAGATCACGTTAAAATTCCATA 4954
SCMV-Rw145 TTACACAATTGGCTTACAGCCCGGGATTATAATAGAATAGGTTGCTCATTAGAACTTGAAGATCACGTTAAAATTCCGTA 4956
4880 4890 4900 4910 4920 4930 4940 4950
9640 bases
13 features
470 475 480 485 490 495
L H N W L T A R D Y N R I G C S L E L E D H V K I P Y
CIP
Consensus CTATATTAGGGGAGTTCCTGACAAGTTGTATGGAAAGCTATATGATATTATTTTACAGTATAGTCCAACTAGTTGTTATG
SCMV-OH CTACATTAGGGGAGTTCCTGACAAGTTGTATGGAAAGCTATATGATATTATTTTACAGTATAGTCCAACTAGTTGCTACG 5036
SCMV-See TTATATACGGGGAGTTCCTGACAAGTTGTATGGGAAGTTATATGATATTATCCTTCAATATAGTCCTACAAGTTGCTATG 5004
SCMV-Rw001 CTACATTAGGGGAGTTCCTGACAAGCTGTATGGAAAGCTATATGATATTATTTTACAGTATAGTCCAACTAGTTGTTATG 5036
SCMV-Rw043 CTATATTAGGGGAGTTCCTGATAAGTTGTATGGAAAGCTATATGATATTATTTTACAGTATAGTCCAACTAGTTGTTATG 5034
SCMV-Rw145 CTATATTAGGGGAGTTCCCGATAAGTTGTATGGAAAGCTATATGATATTATTTTACAGTATAGTCCAACTAGTTGTTATG 5036
4960 4970 4980 4990 5000 5010 5020 5030
9640 bases
13 features
500 505 510 515 520
Y I R G V P D K L Y G K L Y D I I L Q Y S P T S C Y
CIP
Consensus GTAGACTATCAAGTGCGTGTGCAGGTAAAGTAGCATACACTTTGCGAACTGATCCGTGTTCACTTCCAAGAACAATAGCA
SCMV-OH GTAGATTGTCAAGTGCGTGTGCAGGTAAAGTAGCATATACTTTGCGGACTGATCCATGTTCACTTCCAAGAACAATAGCA 5116
SCMV-See GAAGACTATCAAGTGCTTGCGCGGGCAAAGTCGCATATACATTGCGCACTGATCCTTGTTCATTACCAAGAACAATAGCT 5084
SCMV-Rw001 GTAGACTATCAAGTGCGTGTGCAGGTAAAGTAGCATACACTTTGCGAACTGATCCGTGTTCACTTCCAAGAACAATAGCA 5116
SCMV-Rw043 GTAGACTATCAAGTGCGTGTGCAGGTAAAGTAGCATACACTTTGCGAACTGATCCGTGCTCACTTCCAAGAACAATAGCA 5114
SCMV-Rw145 GTAGACTATCAAGTGCGTGTGCAGGTAAAGTAGCATACACTTTGCGAACAGATCCGTGTTCACTTCCAAGAACAATAGCA 5116
5040 5050 5060 5070 5080 5090 5100 5110
9640 bases
13 features
525 530 535 540 545
G R L S S A C A G K V A Y T L R T D P C S L P R T I A
CIP

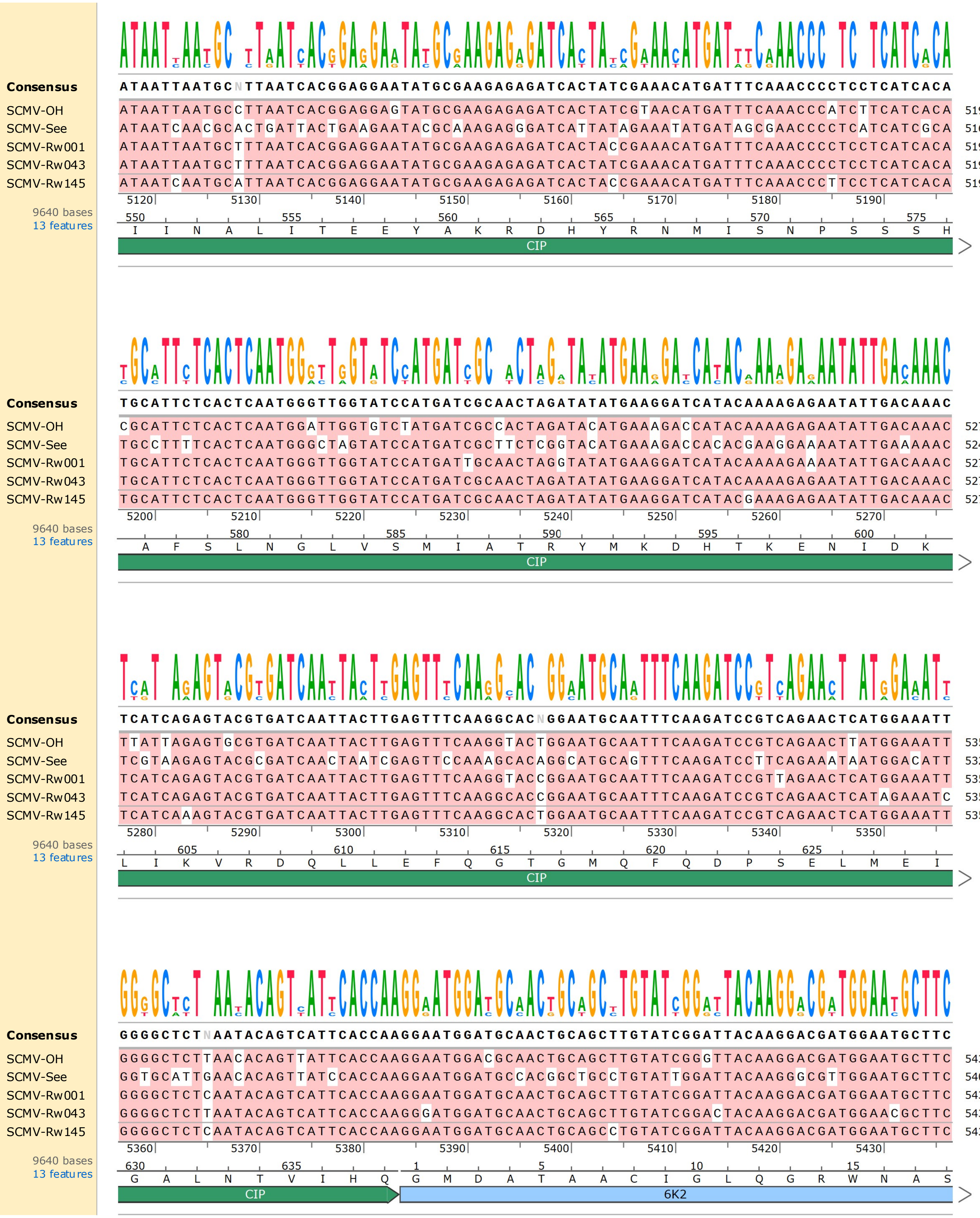
Consensus ATAATTAATGCNTTAATCACGGAGGAATATGCGAAGAGAGATCACTATCGAAACATGATTTCAAACCCCTCCTCATCACA
SCMV-OH ATAATTAATGCCTTAATCACGGAGGAGTATGCGAAGAGAGATCACTATCGTAACATGATTTCAAACCCATCTTCATCACA 5196
SCMV-See ATAATCAACGCACTGATTACTGAAGAATACGCAAAGAGGGATCATTATAGAAATATGATAGCGAACCCCTCATCATCGCA 5164
SCMV-Rw001 ATAATTAATGCTTTAATCACGGAGGAATATGCGAAGAGAGATCACTACCGAAACATGATTTCAAACCCCTCCTCATCACA 5196
SCMV-Rw043 ATAATTAATGCTTTAATCACGGAGGAATATGCGAAGAGAGATCACTATCGAAACATGATTTCAAACCCCTCCTCATCACA 5194
SCMV-Rw145 ATAATCAATGCATTAATCACGGAGGAATATGCGAAGAGAGATCACTACCGAAACATGATTTCAAACCCTTCCTCATCACA 5196
5120 5130 5140 5150 5160 5170 5180 5190
9640 bases
13 features
550 555 560 565 570 575
I I N A L I T E E Y A K R D H Y R N M I S N P S S S H
CIP
Consensus TGCATTCTCACTCAATGGGTTGGTATCCATGATCGCAACTAGATATATGAAGGATCATACAAAAGAGAATATTGACAAAC
SCMV-OH CGCATTCTCACTCAATGGATTGGTGTCTATGATCGCCACTAGATACATGAAAGACCATACAAAAGAGAATATTGACAAAC 5276
SCMV-See TGCCTTTTCACTCAATGGGCTAGTATCCATGATCGCTTCTCGGTACATGAAAGACCACACGAAGGAAAATATTGAAAAAC 5244
SCMV-Rw001 TGCATTCTCACTCAATGGGTTGGTATCCATGATTGCAACTAGGTATATGAAGGATCATACAAAAGAAAATATTGACAAAC 5276
SCMV-Rw043 TGCATTCTCACTCAATGGGTTGGTATCCATGATCGCAACTAGATATATGAAGGATCATACAAAAGAGAATATTGACAAAC 5274
SCMV-Rw145 TGCATTCTCACTCAATGGGTTGGTATCCATGATCGCAACTAGATATATGAAGGATCATACGAAAGAGAATATTGACAAAC 5276
5200 5210 5220 5230 5240 5250 5260 5270
9640 bases
13 features
580 585 590 595 600
A F S L N G L V S M I A T R Y M K D H T K E N I D K
CIP
Consensus TCATCAGAGTACGTGATCAATTACTTGAGTTTCAAGGCACNGGAATGCAATTTCAAGATCCGTCAGAACTCATGGAAATT
SCMV-OH TTATTAGAGTGCGTGATCAATTACTTGAGTTTCAAGGTACTGGAATGCAATTTCAAGATCCGTCAGAACTTATGGAAATT 5356
SCMV-See TCGTAAGAGTACGCGATCAACTAATCGAGTTCCAAAGCACAGGCATGCAGTTTCAAGATCCTTCAGAAATAATGGACATT 5324
SCMV-Rw001 TCATCAGAGTACGTGATCAATTACTTGAGTTTCAAGGTACCGGAATGCAATTTCAAGATCCGTTAGAACTCATGGAAATT 5356
SCMV-Rw043 TCATCAGAGTACGTGATCAATTACTTGAGTTTCAAGGCACCGGAATGCAATTTCAAGATCCGTCAGAACTCATAGAAATC 5354
SCMV-Rw145 TCATCAAAGTACGTGATCAATTACTTGAGTTTCAAGGCACTGGAATGCAATTTCAAGATCCGTCAGAACTCATGGAAATT 5356
5280 5290 5300 5310 5320 5330 5340 5350
9640 bases
13 features
605 610 615 620 625
L I K V R D Q L L E F Q G T G M Q F Q D P S E L M E I
CIP
Consensus GGGGCTCTNAATACAGTCATTCACCAAGGAATGGATGCAACTGCAGCTTGTATCGGATTACAAGGACGATGGAATGCTTC
SCMV-OH GGGGCTCTTAACACAGTTATTCACCAAGGAATGGACGCAACTGCAGCTTGTATCGGGTTACAAGGACGATGGAATGCTTC 5436
SCMV-See GGTGCATTGAACACAGTTATCCACCAAGGAATGGATGCCACGGCTGCCTGTATTGGATTACAAGGGCGTTGGAATGCTTC 5404
SCMV-Rw001 GGGGCTCTCAATACAGTCATTCACCAAGGAATGGATGCAACTGCAGCTTGTATCGGATTACAAGGACGATGGAATGCTTC 5436
SCMV-Rw043 GGGGCTCTTAATACAGTCATTCACCAAGGGATGGATGCAACTGCAGCTTGTATCGGACTACAAGGACGATGGAACGCTTC 5434
SCMV-Rw145 GGGGCTCTCAATACAGTCATTCACCAAGGAATGGATGCAACTGCAGCCTGTATCGGATTACAAGGACGATGGAATGCTTC 5436
5360 5370 5380 5390 5400 5410 5420 5430
9640 bases
13 features
630 635 1 5 10 15
G A L N T V I H Q G M D A T A A C I G L Q G R W N A S
CIP
6K2

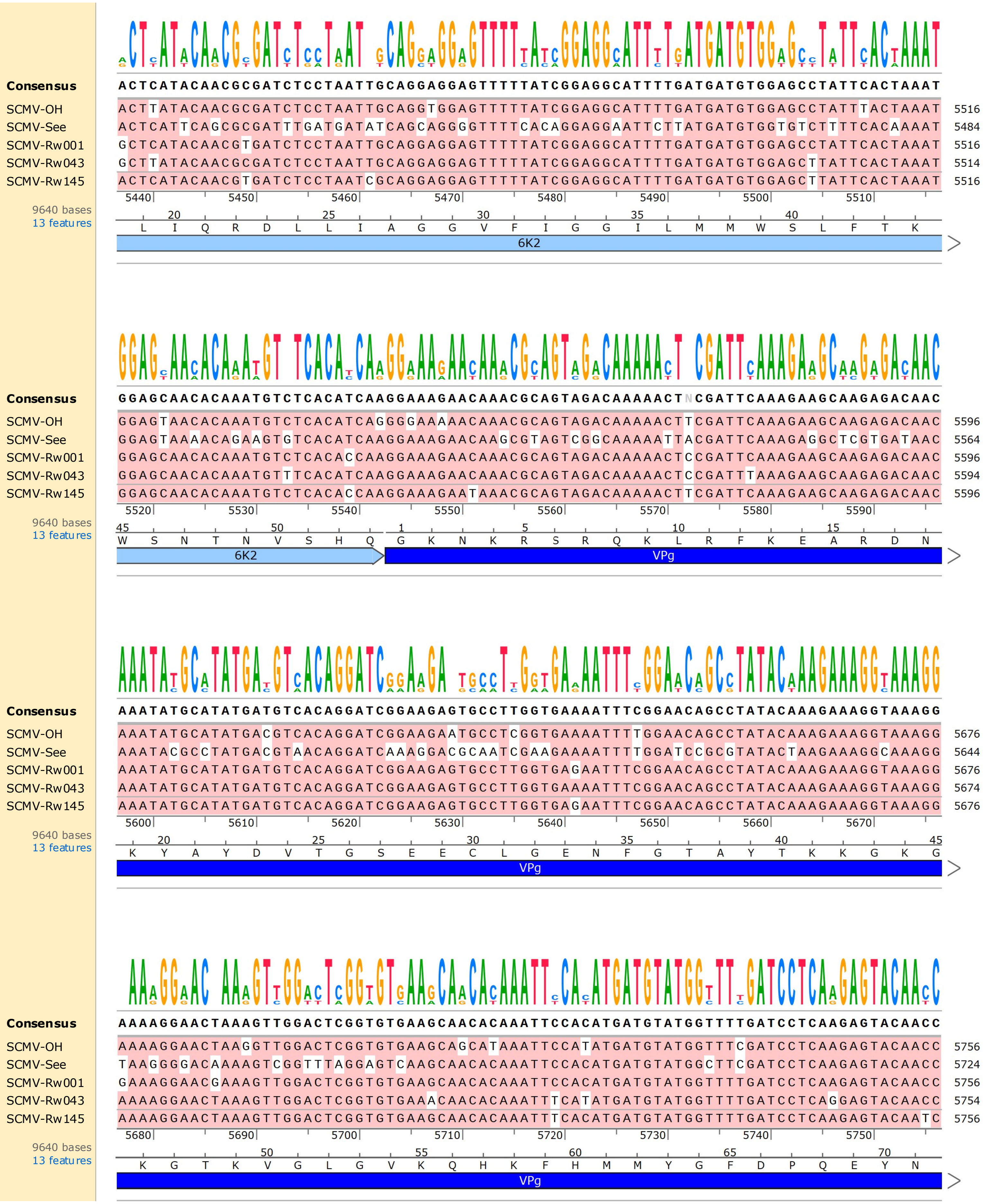

Consensus
ACTCATACAACGCGATCTCCTAATTGCAGGAGGAGTTTTTATCGGAGGCATTTTGATGATGTGGAGCCTATTCACTAAAT
SCMV-OH ACTTATACAACGCGATCTCCTAATTGCAGGTGGAGTTTTTATCGGAGGCATTTTGATGATGTGGAGCCTATTTACTAAAT 5516
SCMV-See ACTCATTCAGCGCGATTTGATGATATCAGCAGGGGTTTTCACAGGAGGAATTCTTATGATGTGGTGTCTTTTCACAAAAT 5484
SCMV-Rw001 GCTCATACAACGTGATCTCCTAATTGCAGGAGGAGTTTTTATCGGAGGCATTTTGATGATGTGGAGCCTATTCACTAAAT 5516
SCMV-Rw043 GCTTATACAACGCGATCTCCTAATTGCAGGAGGAGTTTTTATCGGAGGCATTTTGATGATGTGGAGCTTATTCACTAAAT 5514
SCMV-Rw145 ACTCATACAACGTGATCTCCTAATCGCAGGAGGAGTTTTTATCGGAGGCATTTTGATGATGTGGAGCTTATTCACTAAAT 5516
5440 5450 5460 5470 5480 5490 5500 5510
20 25 30 35 40
L I Q R D L L I A G G V F I G G I L M M W S L F T K
6K2
9640 bases
13 features
Consensus
GGAGCAACACAAATGTCTCACATCAAGGAAAGAACAAACGCAGTAGACAAAAACTNCGATTCAAAGAAGCAAGAGACAAC
SCMV-OH GGAGTAACACAAATGTCTCACATCAGGGGAAAAACAAACGCAGTAGACAAAAACTTCGATTCAAAGAAGCAAGAGACAAC 5596
SCMV-See GGAGTAAAACAGAAGTGTCACATCAAGGAAAGAACAAGCGTAGTCGGCAAAAATTACGATTCAAAGAGGCTCGTGATAAC 5564
SCMV-Rw001 GGAGCAACACAAATGTCTCACACCAAGGAAAGAACAAACGCAGTAGACAAAAACTCCGATTCAAAGAAGCAAGAGACAAC 5596
SCMV-Rw043 GGAGCAACACAAATGTTTCACATCAAGGAAAGAACAAACGCAGTAGACAAAAACTCCGATTTAAAGAAGCAAGAGACAAC 5594
SCMV-Rw145 GGAGCAACACAAATGTCTCACACCAAGGAAAGAATAAACGCAGTAGACAAAAACTTCGATTCAAAGAAGCAAGAGACAAC 5596
5520 5530 5540 5550 5560 5570 5580 5590
45 50 1 5 10 15
W S N T N V S H Q G K N K R S R Q K L R F K E A R D N
6K2
VPg
9640 bases
13 features
Consensus
AAATATGCATATGATGTCACAGGATCGGAAGAGTGCCTTGGTGAAAATTTCGGAACAGCCTATACAAAGAAAGGTAAAGG
SCMV-OH AAATATGCATATGACGTCACAGGATCGGAAGAATGCCTCGGTGAAAATTTTGGAACAGCCTATACAAAGAAAGGTAAAGG 5676
SCMV-See AAATACGCCTATGACGTAACAGGATCAAAGGACGCAATCGAAGAAAATTTTGGATCCGCGTATACTAAGAAAGGCAAAGG 5644
SCMV-Rw001 AAATATGCATATGATGTCACAGGATCGGAAGAGTGCCTTGGTGAGAATTTCGGAACAGCCTATACAAAGAAAGGTAAAGG 5676
SCMV-Rw043 AAATATGCATATGATGTCACAGGATCGGAAGAGTGCCTTGGTGAAAATTTCGGAACAGCCTATACAAAGAAAGGTAAAGG 5674
SCMV-Rw145 AAATATGCATATGATGTCACAGGATCGGAAGAGTGCCTTGGTGAGAATTTCGGAACAGCCTATACAAAGAAAGGTAAAGG 5676
5600 5610 5620 5630 5640 5650 5660 5670
20 25 30 35 40 45
K Y A Y D V T G S E E C L G E N F G T A Y T K K G K G
VPg
9640 bases
13 features
Consensus
AAAAGGAACTAAAGTTGGACTCGGTGTGAAGCAACACAAATTCCACATGATGTATGGTTTTGATCCTCAAGAGTACAACC
SCMV-OH AAAAGGAACTAAGGTTGGACTCGGTGTGAAGCAGCATAAATTCCATATGATGTATGGTTTCGATCCTCAAGAGTACAACC 5756
SCMV-See TAAGGGGACAAAAGTCGGTTTAGGAGTCAAGCAACACAAATTCCACATGATGTATGGCTTCGATCCTCAAGAGTACAACC 5724
SCMV-Rw001 GAAAGGAACGAAAGTTGGACTCGGTGTGAAGCAACACAAATTCCACATGATGTATGGTTTTGATCCTCAAGAGTACAACC 5756
SCMV-Rw043 AAAAGGAACTAAAGTTGGACTCGGTGTGAAACAACACAAATTTCATATGATGTATGGTTTTGATCCTCAGGAGTACAACC 5754
SCMV-Rw145 AAAAGGAACTAAAGTTGGACTCGGTGTGAAGCAACACAAATTTCACATGATGTATGGTTTTGATCCTCAAGAGTACAATC 5756
5680 5690 5700 5710 5720 5730 5740 5750
50 55 60 65 70
K G T K V G L G V K Q H K F H M M Y G F D P Q E Y N
VPg
9640 bases
13 features

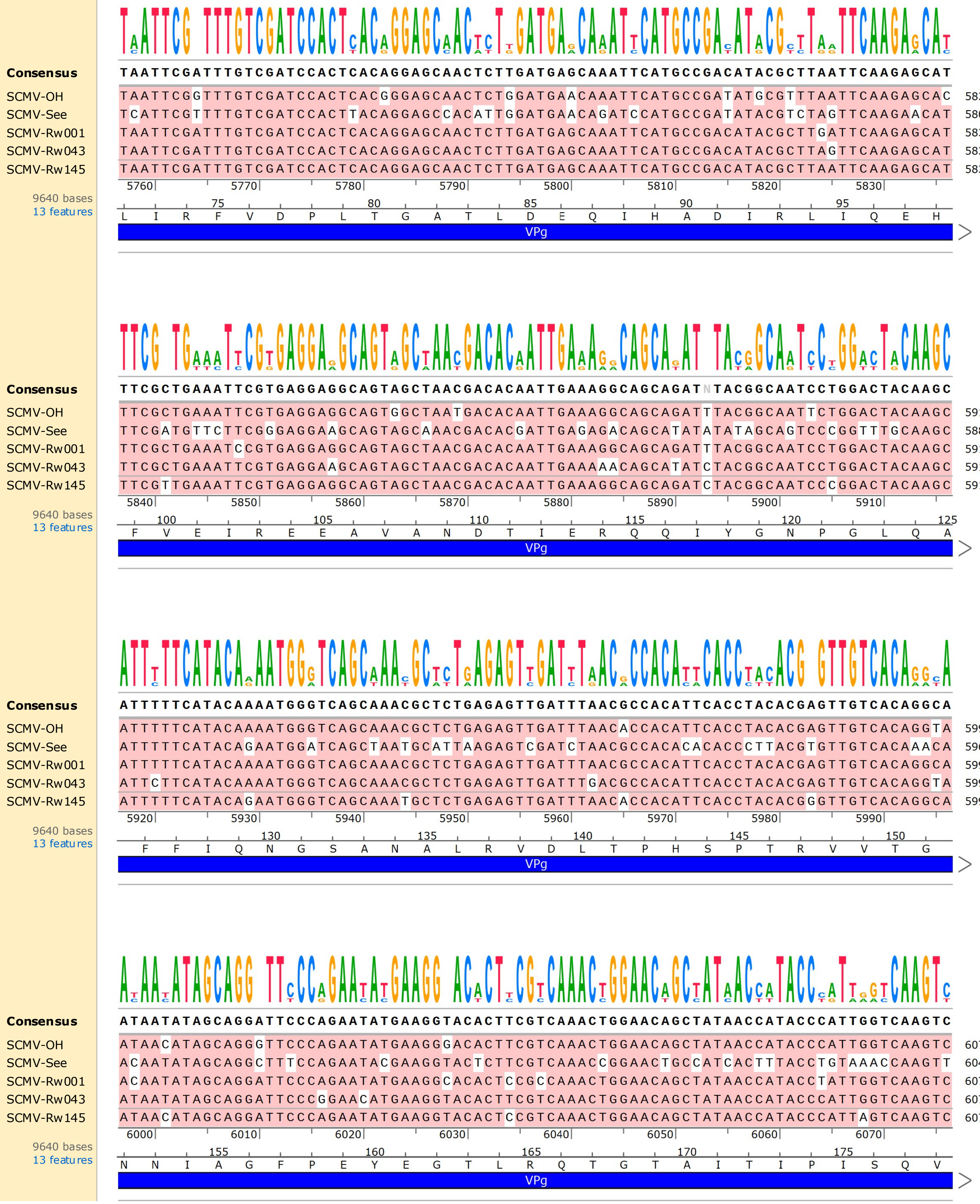
Consensus
TAATTCGATTTGTCGATCCACTCACAGGAGCAACTCTTGATGAGCAAATTCATGCCGACATACGCTTAATTCAAGAGCAT
SCMV-OH TAATTCGGTTTGTCGATCCACTCACGGGAGCAACTCTGGATGAACAAATTCATGCCGATATGCGTTTAATTCAAGAGCAC 5836
SCMV-See TCATTCGTTTTGTCGATCCACTTACAGGAGCCACATTGGATGAACAGATCCATGCCGATATACGTCTAGTTCAAGAACAT 5804
SCMV-Rw001 TAATTCGATTTGTCGATCCACTCACAGGAGCAACTCTTGATGAGCAAATTCATGCCGACATACGCTTGATTCAAGAGCAT 5836
SCMV-Rw043 TAATTCGATTTGTCGATCCACTCACAGGAGCAACTCTTGATGAGCAAATTCATGCCGACATACGCTTAGTTCAAGAGCAT 5834
SCMV-Rw145 TAATTCGATTTGTCGATCCACTCACAGGAGCAACTCTTGATGAGCAAATTCATGCCGACATACGCTTAATTCAAGAGCAT 5836
5760 5770 5780 5790 5800 5810 5820 5830
9640 bases
13 features
75 80 85 90 95
L I R F V D P L T G A T L D E Q I H A D I R L I Q E H
VPg
Consensus
TTCGCTGAAATTCGTGAGGAGGCAGTAGCTAACGACACAATTGAAAGGCAGCAGATNTACGGCAATCCTGGACTACAAGC
SCMV-OH TTCGCTGAAATTCGTGAGGAGGCAGTGGCTAATGACACAATTGAAAGGCAGCAGATTTACGGCAATTCTGGACTACAAGC 5916
SCMV-See TTCGATGTTCTTCGGGAGGAAGCAGTAGCAAACGACACGATTGAGAGACAGCATATATATAGCAGTCCCGGTTTGCAAGC 5884
SCMV-Rw001 TTCGCTGAAATCCGTGAGGAGGCAGTAGCTAACGACACAATTGAAAGGCAGCAGATTTACGGCAATCCTGGACTACAAGC 5916
SCMV-Rw043 TTCGCTGAAATTCGTGAGGAAGCAGTAGCTAACGACACAATTGAAAAACAGCATATCTACGGCAATCCTGGACTACAAGC 5914
SCMV-Rw145 TTCGTTGAAATTCGTGAGGAGGCAGTAGCTAACGACACAATTGAAAGGCAGCAGATCTACGGCAATCCCGGACTACAAGC 5916
5840 5850 5860 5870 5880 5890 5900 5910
9640 bases
13 features
100 105 110 115 120 125
F V E I R E E A V A N D T I E R Q Q I Y G N P G L Q A
VPg
Consensus
ATTTTTCATACAAAATGGGTCAGCAAACGCTCTGAGAGTTGATTTAACGCCACATTCACCTACACGAGTTGTCACAGGCA
SCMV-OH ATTTTTCATACAAAATGGGTCAGCAAACGCTCTGAGAGTTGATTTAACACCACATTCACCTACACGAGTTGTCACAGGTA 5996
SCMV-See ATTTTTCATACAGAATGGATCAGCTAATGCATTAAGAGTCGATCTAACGCCACACACACCCTTACGTGTTGTCACAAACA 5964
SCMV-Rw001 ATTTTTCATACAAAATGGGTCAGCAAACGCTCTGAGAGTTGATTTAACGCCACATTCACCTACACGAGTTGTCACAGGCA 5996
SCMV-Rw043 ATTCTTCATACAAAATGGGTCAGCAAACGCTCTGAGAGTTGATTTGACGCCACATTCACCTACACGAGTTGTCACAGGTA 5994
SCMV-Rw145 ATTTTTCATACAGAATGGGTCAGCAAATGCTCTGAGAGTTGATTTAACACCACATTCACCTACACGGGTTGTCACAGGCA 5996
5920 5930 5940 5950 5960 5970 5980 5990
9640 bases
13 features
130 135 140 145 150
F F I Q N G S A N A L R V D L T P H S P T R V V T G
VPg
Consensus
ATAATATAGCAGGATTCCCAGAATATGAAGGTACACTTCGTCAAACTGGAACAGCTATAACCATACCCATTGGTCAAGTC
SCMV-OH ATAACATAGCAGGGTTCCCAGAATATGAAGGGACACTTCGTCAAACTGGAACAGCTATAACCATACCCATTGGTCAAGTC 6076
SCMV-See ACAATATAGCAGGCTTTCCAGAATACGAAGGTACTCTTCGTCAAACCGGAACTGCCATCACTTTACCTGTAAACCAAGTT 6044
SCMV-Rw001 ACAATATAGCAGGATTCCCAGAATATGAAGGCACACTCCGCCAAACTGGAACAGCTATAACCATACCTATTGGTCAAGTC 6076
SCMV-Rw043 ATAATATAGCAGGATTCCCGGAACATGAAGGTACACTTCGTCAAACTGGAACAGCTATAACCATACCCATTGGTCAAGTC 6074
SCMV-Rw145 ATAACATAGCAGGATTCCCAGAATATGAAGGTACACTCCGTCAAACTGGAACAGCTATAACCATACCCATTAGTCAAGTC 6076
6000 6010 6020 6030 6040 6050 6060 6070
9640 bases
13 features
155 160 165 170 175
N N I A G F P E Y G T L R Q T G T A I T I P I S Q V
VPg

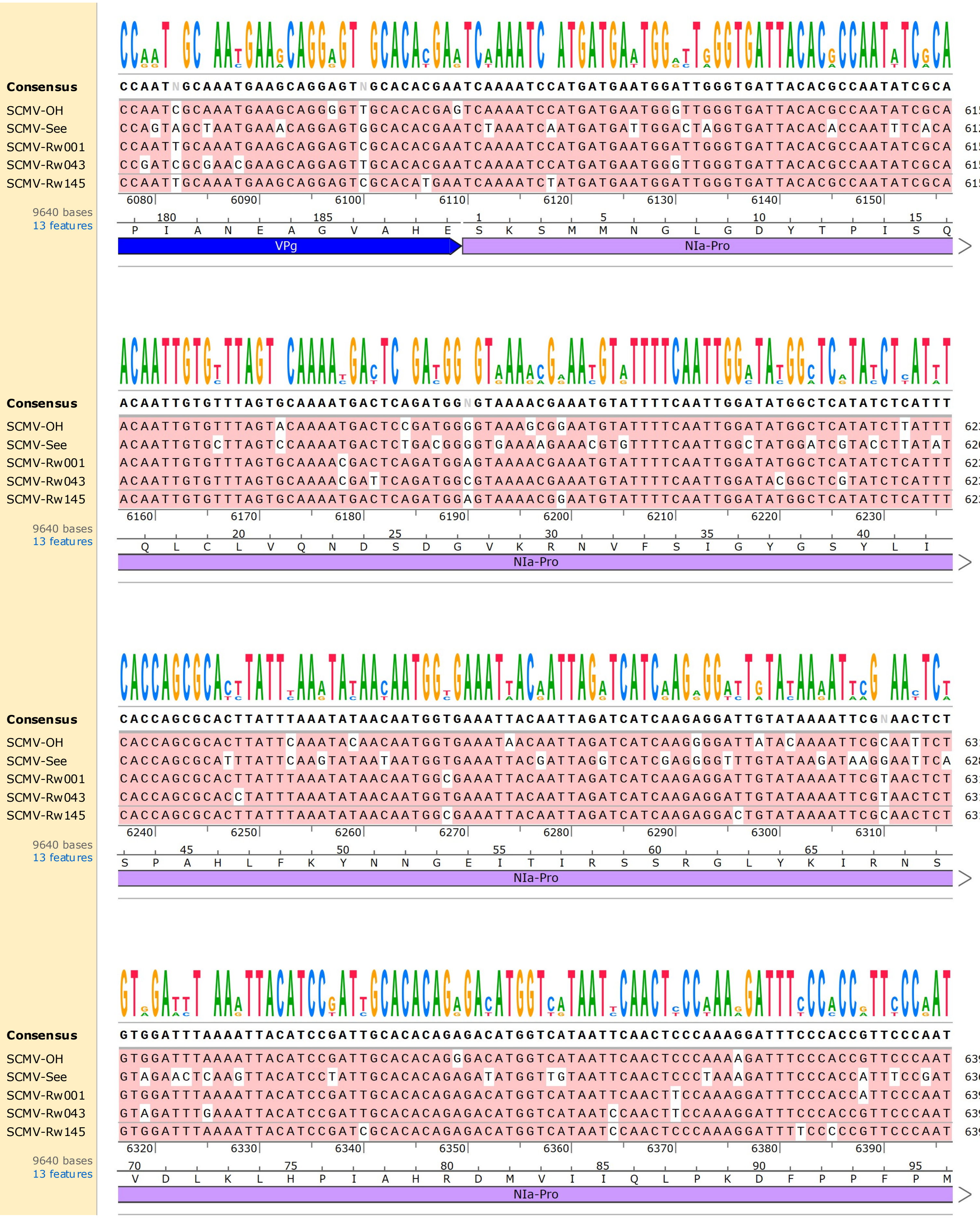
Consensus
CCAATNGCAAATGAAGCAGGAGTNGCACACGAATCAAAATCCATGATGAATGGATTGGGTGATTACACGCCAATATCGCA
SCMV-OH CCAATCGCAAATGAAGCAGGGGTTGCACACGAGTCAAAATCCATGATGAATGGGTTGGGTGATTACACGCCAATATCGCA 6156
SCMV-See CCAGTAGCTAATGAAACAGGAGTGGCACACGAATCTAAATCAATGATGATTGGACTAGGTGATTACACACCAATTTCACA 6124
SCMV-Rw001 CCAATTGCAAATGAAGCAGGAGTCGCACACGAATCAAAATCCATGATGAATGGATTGGGTGATTACACGCCAATATCGCA 6156
SCMV-Rw043 CCGATCGCGAACGAAGCAGGAGTTGCACACGAATCAAAATCCATGATGAATGGGTTGGGTGATTACACGCCAATATCGCA 6154
SCMV-Rw145 CCAATTGCAAATGAAGCAGGAGTCGCACATGAATCAAAATCTATGATGAATGGATTGGGTGATTACACGCCAATATCGCA 6156
6080 6090 6100 6110 6120 6130 6140 6150
9640 bases
13 features
180 185 1 5 10 15
P I A N E A G V A H E S K S M M N G L G D Y T P I S Q
VPg
NIa-Pro
Consensus
ACAATTGTGTTTAGTGCAAAATGACTCAGATGGNGTAAAACGAAATGTATTTTCAATTGGATATGGCTCATATCTCATTT
SCMV-OH ACAATTGTGTTTAGTACAAAATGACTCCGATGGGGTAAAGCGGAATGTATTTTCAATTGGATATGGCTCATATCTTATTT 6236
SCMV-See ACAATTGTGCTTAGTCCAAAATGACTCTGACGGGGTGAAAAGAAACGTGTTTTCAATTGGCTATGGATCGTACCTTATAT 6204
SCMV-Rw001 ACAATTGTGTTTAGTGCAAAACGACTCAGATGGAGTAAAACGAAATGTATTTTCAATTGGATATGGCTCATATCTCATTT 6236
SCMV-Rw043 ACAATTGTGTTTAGTGCAAAACGATTCAGATGGCGTAAAACGAAATGTATTTTCAATTGGATACGGCTCGTATCTCATTT 6234
SCMV-Rw145 ACAATTGTGTTTAGTGCAAAATGACTCAGATGGAGTAAAACGGAATGTATTTTCAATTGGATATGGCTCATATCTCATTT 6236
6160 6170 6180 6190 6200 6210 6220 6230
9640 bases
13 features
20 25 30 35 40
Q L C L V Q N D S D G V K R N V F S I G Y G S Y L I
NIa-Pro
Consensus
CACCAGCGCACTTATTTAAATATAACAATGGTGAAATTACAATTAGATCATCAAGAGGATTGTATAAAATTCGNAACTCT
SCMV-OH CACCAGCGCACTTATTCAAATACAACAATGGTGAAATAACAATTAGATCATCAAGGGGATTATACAAAATTCGCAATTCT 6316
SCMV-See CACCAGCGCATTTATTCAAGTATAATAATGGTGAAATTACGATTAGGTCATCGAGGGGTTGTATAAGATAAGGAATTCA 6284
SCMV-Rw001 CACCAGCGCACTTATTTAAATATAACAATGGCGAAATTACAATTAGATCATCAAGAGGATTGTATAAAATTCGTAACTCT 6316
SCMV-Rw043 CACCAGCGCACCTATTTAAATATAACAATGGTGAAATTACAATTAGATCATCAAGAGGATTGTATAAAATTCGTAACTCT 6314
SCMV-Rw145 CACCAGCGCACTTATTTAAATATAACAATGGCGAAATTACAATTAGATCATCAAGAGGACTGTATAAAATTCGCAACTCT 6316
6240 6250 6260 6270 6280 6290 6300 6310
9640 bases
13 features
45 50 55 60 65
S P A H L F K Y N N G E I T I R S S R G L Y K I R N S
NIa-Pro
Consensus
GTGGATTTAAAATTACATCCGATTGCACACAGAGACATGGTCATAATTCAACTCCCAAAGGATTTCCCACCGTTCCCAAT
SCMV-OH GTGGATTTAAAATTACATCCGATTGCACACAGGGACATGGTCATAATTCAACTCCCAAAAGATTTCCCACCGTTCCCAAT 6396
SCMV-See GTAGAACTCAAGTTACATCCTATTGCACACAGAGATATGGTTGTAATTCAACTCCCTAAAGATTTCCCACCATTTCCGAT 6364
SCMV-Rw001 GTGGATTTAAAATTACATCCGATTGCACACAGAGACATGGTCATAATTCAACTTCCAAAGGATTTCCCACCATTCCCAAT 6396
SCMV-Rw043 GTAGATTTGAAATTACATCCGATTGCACACAGAGACATGGTCATAATCCAACTTCCAAAGGATTTCCCACCGTTCCCAAT 6394
SCMV-Rw145 GTGGATTTAAAATTACATCCGATCGCACACAGAGACATGGTCATAATCCAACTCCCAAAGGATTTTCCCCCGTTCCCAAT 6396
6320 6330 6340 6350 6360 6370 6380 6390
9640 bases
13 features
70 75 80 85 90 95
V D L K L H P I A H R D M V I Q L P K D F P P F P M
NIa-Pro

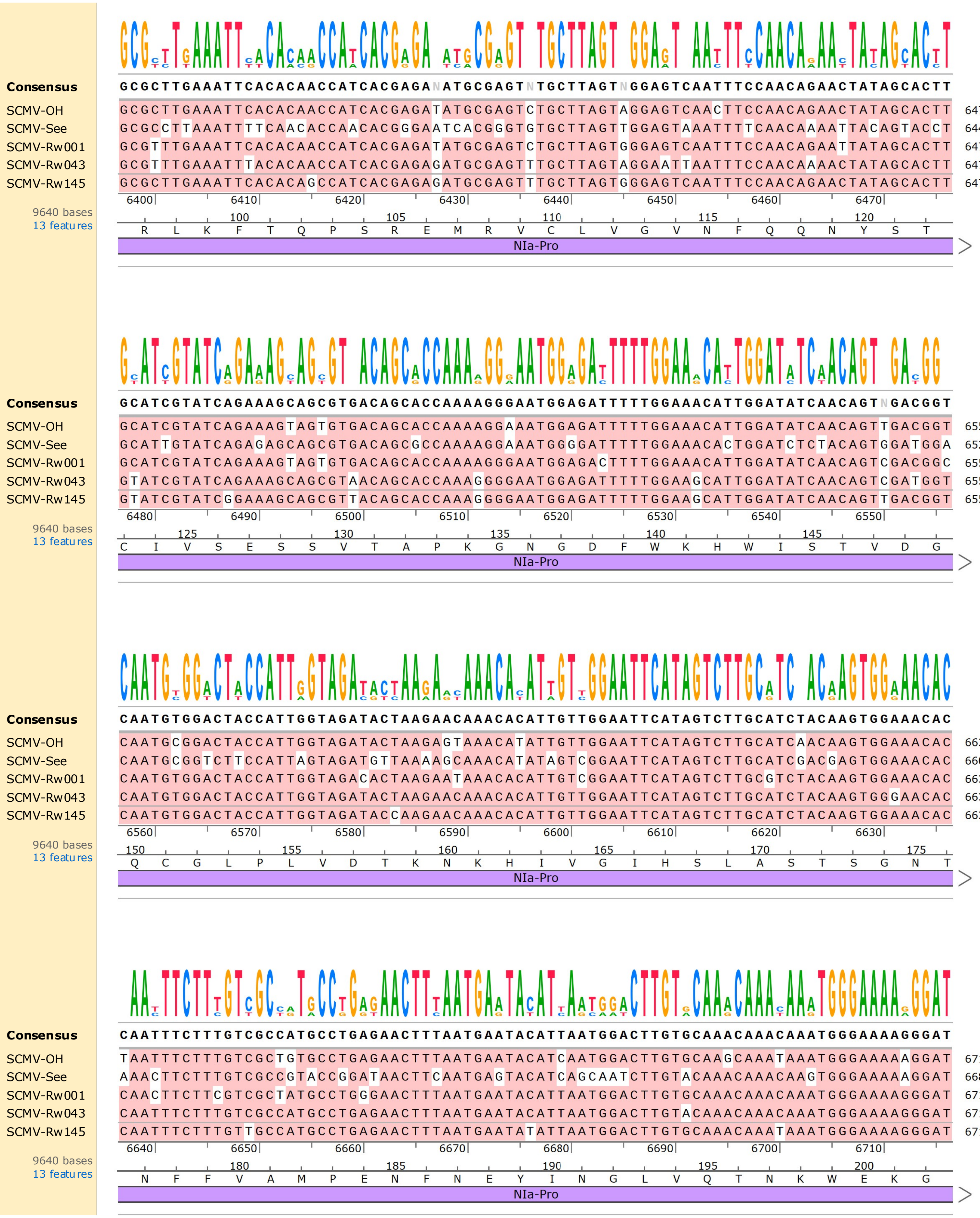
Consensus
GCGCTTGAAATTCACACAACCATCACGAGANATGCGAGTNTGCTTAGTNGGAGTCAATTTCCAACAGAACTATAGCACTT
SCMV-OH GCGCTTGAAATTCACACAACCATCACGAGATATGCGAGTCTGCTTAGTAGGAGTCAACTTCCAACAGAACTATAGCACTT 6476
SCMV-See GCGCCTTAAATTTTCAACACCAACACGGGAATCACGGGTGTGCTTAGTTGGAGTAAATTTTCAACAAAATTACAGTACCT 6444
SCMV-Rw001 GCGTTTGAAATTCACACAACCATCACGAGATATGCGAGTCTGCTTAGTGGGAGTCAATTTCCAACAGAATTATAGCACTT 6476
SCMV-Rw043 GCGTTTGAAATTTACACAACCATCACGAGAGATGCGAGTTTGCTTAGTAGGAATTAATTTCCAACAAAACTATAGCACTT 6474
SCMV-Rw145 GCGCTTGAAATTCACACAGCCATCACGAGAGATGCGAGTTTGCTTAGTGGGAGTCAATTTCCAACAGAACTATAGCACTT 6476
6400 6410 6420 6430 6440 6450 6460 6470
9640 bases
13 features
100 105 110 115 120
R L K F T Q P S R E M R V C L V G V N F Q Q N Y S T
NIa-Pro
Consensus
GCATCGTATCAGAAAGCAGCGTGACAGCACCAAAAGGGAATGGAGATTTTTGGAAACATTGGATATCAACAGTNGACGGT
SCMV-OH GCATCGTATCAGAAAGTAGTGTGACAGCACCAAAAGGAAATGGAGATTTTTGGAAACATTGGATATCAACAGTTGACGGT 6556
SCMV-See GCATTGTATCAGAGAGCAGCGTGACAGCGCCAAAAGGAAATGGGGATTTTTGGAAACACTGGATCTCTACAGTGGATGGA 6524
SCMV-Rw001 GCATCGTATCAGAAAGTAGTGTGACAGCACCAAAAGGGAATGGAGACTTTTGGAAACATTGGATATCAACAGTCGACGGC 6556
SCMV-Rw043 GTATCGTATCAGAAAGCAGCGTAACAGCACCAAAGGGGAATGGAGATTTTTGGAAGCATTGGATATCAACAGTCGATGGT 6554
SCMV-Rw145 GTATCGTATCGGAAAGCAGCGTTACAGCACCAAAGGGGAATGGAGATTTTTGGAAGCATTGGATATCAACAGTTGACGGT 6556
6480 6490 6500 6510 6520 6530 6540 6550
9640 bases
13 features
125 130 135 140 145
C I V S E S S V T A P K G N G D F W K H W I S T V D G
NIa-Pro
Consensus
CAATGTGGACTACCATTGGTAGATACTAAGAACAAACACATTGTTGGAATTCATAGTCTTGCATCTACAAGTGGAAACAC
SCMV-OH CAATGCGGACTACCATTGGTAGATACTAAGAGTAAACATATTGTTGGAATTCATAGTCTTGCATCAACAAGTGGAAACAC 6636
SCMV-See CAATGCGGTCTTCCATTAGTAGATGTTAAAAGCAAACATATAGTCGGAATTCATAGTCTTGCATCGACGAGTGGAAACAC 6604
SCMV-Rw001 CAATGTGGACTACCATTGGTAGACACTAAGAATAAACACATTGTCGGAATTCATAGTCTTGCGTCTACAAGTGGAAACAC 6636
SCMV-Rw043 CAATGTGGACTACCATTGGTAGATACTAAGAACAAACACATTGTTGGAATTCATAGTCTTGCATCTACAAGTGGGAACAC 6634
SCMV-Rw145 CAATGTGGACTACCATTGGTAGATACCAAGAACAAACACATTGTTGGAATTCATAGTCTTGCATCTACAAGTGGAAACAC 6636
6560 6570 6580 6590 6600 6610 6620 6630
9640 bases
13 features
150 155 160 165 170 175
Q C G L P L V D T K N K H I V G I H S L A S T S G N T
NIa-Pro
Consensus
CAATTTCTTTGTCGCCATGCCTGAGAACTTTAATGAATACATTAATGGACTTGTGCAAACAAACAAATGGGAAAAGGGAT
SCMV-OH TAATTTCTTTGTCGCTGTGCCTGAGAACTTTAATGAATACATCAATGGACTTGTGCAAGCAAATAAATGGGAAAAAGGAT 6716
SCMV-See AAACTTCTTTGTCGCCGTACCGGATAACTTCAATGAGTACATCAGCAATCTTGTACAAACAAACAAGTGGGAAAAAGGAT 6684
SCMV-Rw001 CAACTTCTTCGTCGCTATGCCTGGGAACTTTAATGAATACATTAATGGACTTGTGCAAACAAACAAATGGGAAAGGGAT 6716
SCMV-Rw043 CAATTTCTTTGTCGCCATGCCTGAGAACTTTAATGAATACATTAATGGACTTGTACAAACAAACAAATGGGAAAGGGAT 6714
SCMV-Rw145 CAATTTCTTTGTTGCCATGCCTGAGAACTTTAATGAATATATTAATGGACTTGTGCAAACAAATAAATGGGAAAGGGAT 6716
6640 6650 6660 6670 6680 6690 6700 6710
9640 bases
13 features
180 185 190 195 200
N F F V A M P E N F N E Y I N G L V Q T N K W E K G
NIa-Pro

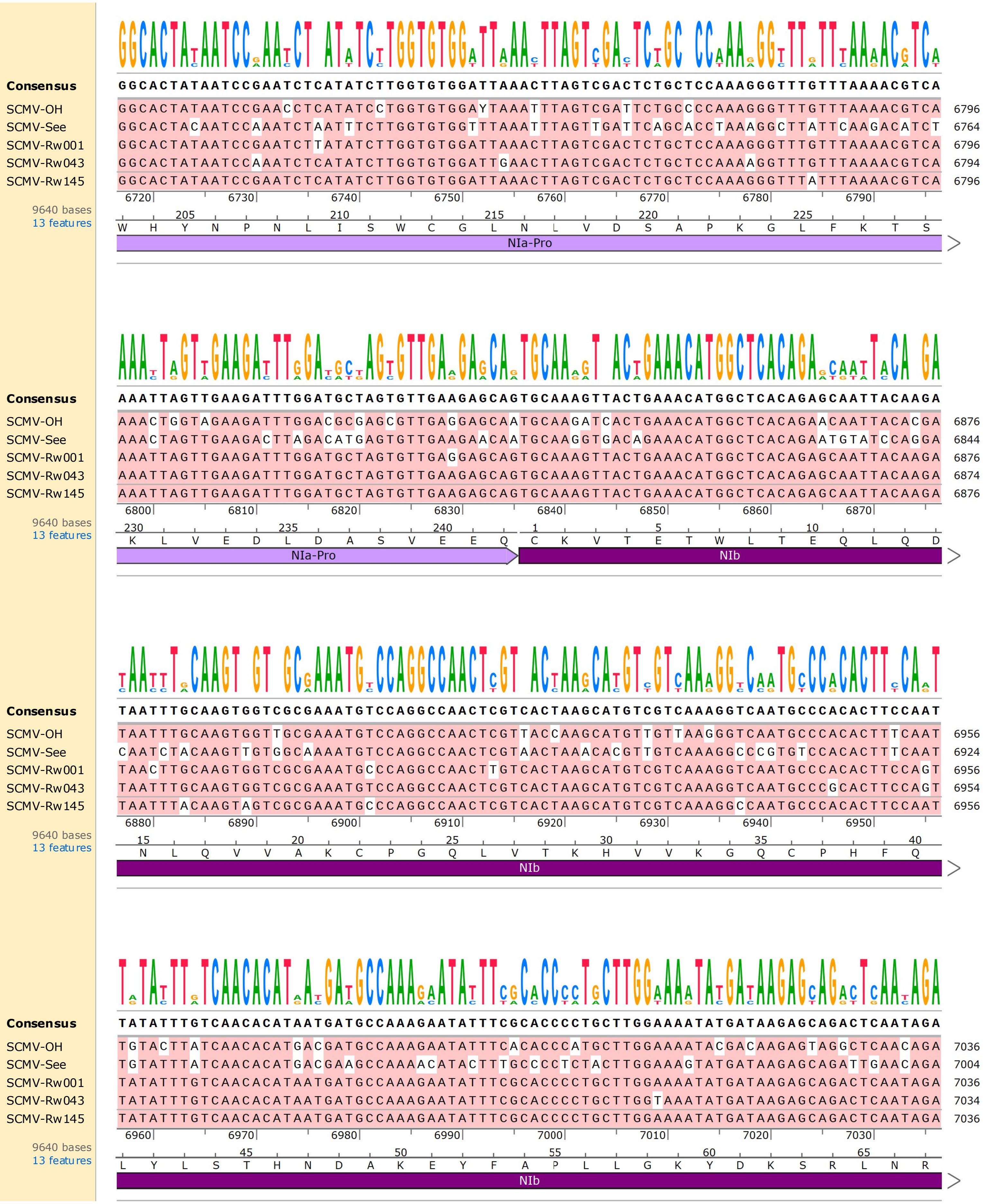
Consensus GGCACTATAATCCGAATCTCATATCTTGGTGTGGATTAAACTTAGTCGACTCTGCTCCAAAGGGTTTGTTTAAAACGTCA
SCMV-OH GGCACTATAATCCGAACCTCATATCCTGGTGTGGAYTAAATTTAGTCGATTCTGCCCCAAAGGGTTTGTTTAAAACGTCA 6796
SCMV-See GGCACTACAATCCAAATCTAATTTCTTGGTGTGGTTTAAATTTAGTTGATTCAGCACCTAAAGGCTTATTCAAGACATCT 6764
SCMV-Rw001 GGCACTATAATCCGAATCTTATATCTTGGTGTGGATTAAACTTAGTCGACTCTGCTCCAAAGGGTTTGTTTAAAACGTCA 6796
SCMV-Rw043 GGCACTATAATCCAAATCTCATATCTTGGTGTGGATTGAACTTAGTCGACTCTGCTCCAAAAGGTTTGTTTAAAACGTCA 6794
SCMV-Rw145 GGCACTATAATCCGAATCTCATATCTTGGTGTGGATTAAACTTAGTCGACTCTGCTCCAAAGGGTTTATTTAAAACGTCA 6796
6720 6730 6740 6750 6760 6770 6780 6790
9640 bases
13 features
205 210 215 220 225
W H Y N P N L I S W C G L N L V D S A P K G L F K T S
NIa-Pro
Consensus AAATTAGTTGAAGATTTGGATGCTAGTGTTGAAGAGCAGTGCAAAGTTACTGAAACATGGCTCACAGAGCAATTACAAGA
SCMV-OH AAACTGGTAGAAGATTTGGACGCGAGCGTTGAGGAGCAATGCAAGATCACTGAAACATGGCTCACAGAACAATTACACGA 6876
SCMV-See AAACTAGTTGAAGACTTAGACATGAGTGTTGAAGAACAATGCAAGGTGACAGAAACATGGCTCACAGAATGTATCCAGGA 6844
SCMV-Rw001 AAATTAGTTGAAGATTTGGATGCTAGTGTTGAGGAGCAGTGCAAAGTTACTGAAACATGGCTCACAGAGCAATTACAAGA 6876
SCMV-Rw043 AAATTAGTTGAAGATTTGGATGCTAGTGTTGAAGAGCAGTGCAAAGTTACTGAAACATGGCTCACAGAGCAATTACAAGA 6874
SCMV-Rw145 AAATTAGTTGAAGATTTGGATGCTAGTGTTGAAGAGCAGTGCAAAGTTACTGAAACATGGCTCACAGAGCAATTACAAGA 6876
6800 6810 6820 6830 6840 6850 6860 6870
9640 bases
13 features
230 235 240 1 5 10
K L V E D L D A S V E E Q C K V T E T W L T E Q L Q D
NIa-Pro
NIb
Consensus TAATTTGCAAGTGGTCGCGAAATGTCCAGGCCAACTCGTCACTAAGCATGTCGTCAAAGGTCAATGCCCACACTTCCAAT
SCMV-OH TAATTTGCAAGTGGTTGCGAAATGTCCAGGCCAACTCGTTACCAAGCATGTTGTTAAGGGTCAATGCCCACACTTTCAAT 6956
SCMV-See CAATCTACAAGTTGTGGCAAAATGTCCAGGCCAACTCGTAACTAAACACGTTGTCAAAGGCCCGTGTCCACACTTTCAAT 6924
SCMV-Rw001 TAACTTGCAAGTGGTCGCGAAATGCCCAGGCCAACTTGTCACTAAGCATGTCGTCAAAGGTCAATGCCCACACTTCCAGT 6956
SCMV-Rw043 TAATTTGCAAGTGGTCGCGAAATGTCCAGGCCAACTCGTCACTAAGCATGTCGTCAAAGGTCAATGCCCGCACTTCCAGT 6954
SCMV-Rw145 TAATTTACAAGTAGTCGCGAAATGCCCAGGCCAACTCGTCACTAAGCATGTCGTCAAAGGCCAATGCCCACACTTCCAAT 6956
6880 6890 6900 6910 6920 6930 6940 6950
9640 bases
13 features
15 20 25 30 35 40
N L Q V V A K C P G Q L V T K H V V K G Q C P H F Q
NIb
Consensus TATATTTGTCAACACATAATGATGCCAAAGAATATTTCGCACCCCTGCTTGGAAAATATGATAAGAGCAGACTCAATAGA
SCMV-OH TGTACTTATCAACACATGACGATGCCAAAGAATATTTCACACCCATGCTTGGAAAATACGACAAGAGTAGGCTCAACAGA 7036
SCMV-See TGTATTTATCAACACATGACGAAGCCAAAACATACTTTGCCCCTCTACTTGGAAAGTATGATAAGAGCAGATTGAACAGA 7004
SCMV-Rw001 TATATTTGTCAACACATAATGATGCCAAAGAATATTTCGCACCCCTGCTTGGAAAATATGATAAGAGCAGACTCAATAGA 7036
SCMV-Rw043 TATATTTGTCAACACATAATGATGCCAAAGAATATTTCGCACCCCTGCTTGGTAAATATGATAAGAGCAGACTCAATAGA 7034
SCMV-Rw145 TATATTTGTCAACACATAATGATGCCAAAGAATATTTCGCACCCCTGCTTGGAAAATATGATAAGAGCAGACTCAATAGA 7036
6960 6970 6980 6990 7000 7010 7020 7030
9640 bases
13 features
45 50 55 60 65
L Y L S T H N D A K E Y F A P L L G K Y D K S R L N R
NIb

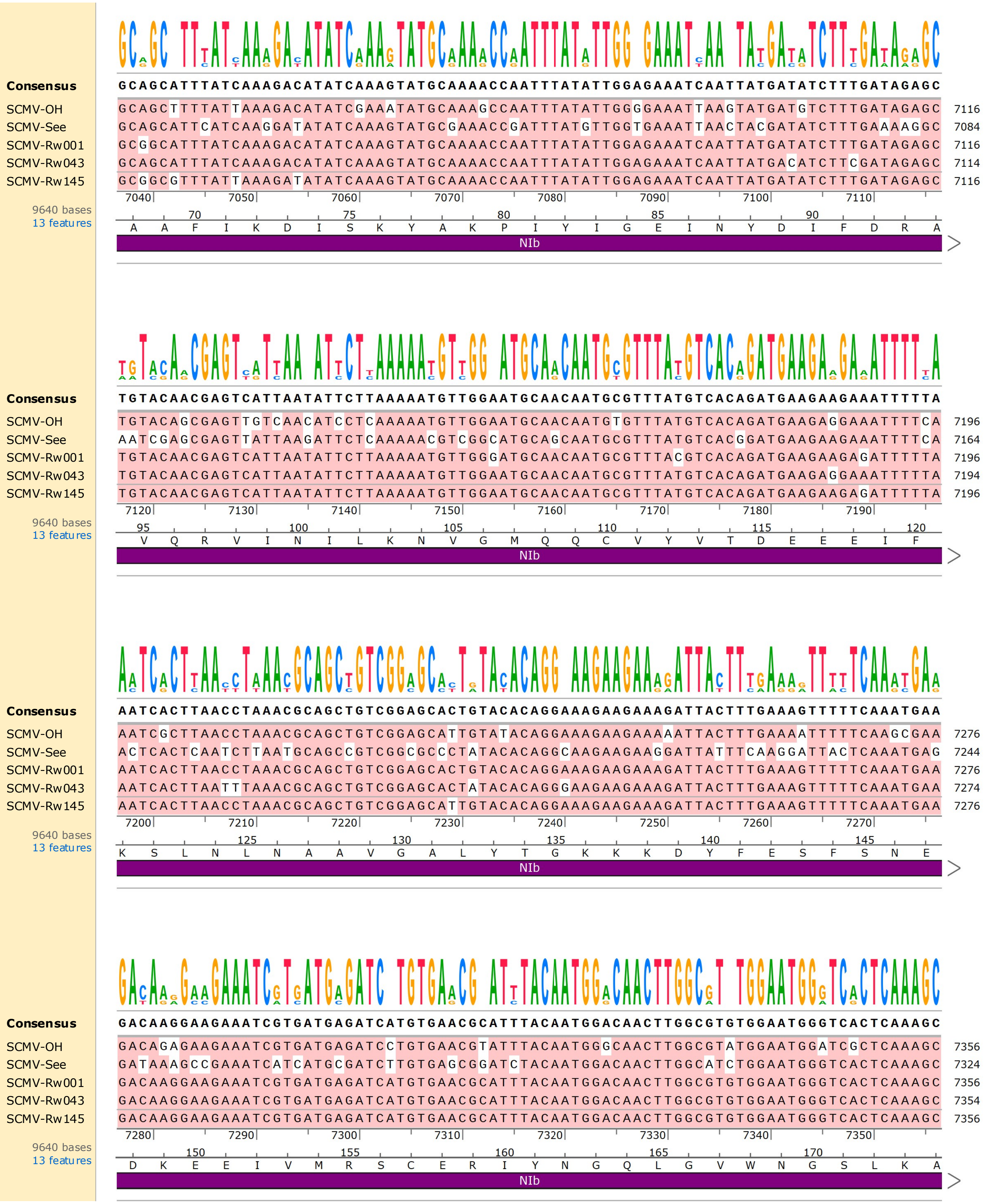
Consensus
SCMV-OH
SCMV-See
SCMV-Rw001
SCMV-Rw043
SCMV-Rw145
9640 bases
13 features
NIb

```
Consensus    TGAAATCAGNCCAATAGAGAAAACCATGTTAAATAAGACTCGAACCTTTACAGCAGCTCCATTGGAAACTTTGCTTGGAG
SCMV-OH      TGAGATCAGACCAATAGAGAAAACCATGCTGAATAAGACTCGAACCTTCACAGCAGCCCCATTAGAAACTTTGCTCGGAG  7436
SCMV-See     TGAAATACGCCCAATTGAGAAAACCATGTTGAATAAAACACGCACTTTCACAGCAGCACCATTGGAAACTCTACTTGGCG  7404
SCMV-Rw001   TGAAATCAGGCCAATAGAGAAAACCATGTTAAATAAGACTCGAACCTTTACAGCAGCTCCATTGGAAACTTTGCTTGGAG  7436
SCMV-Rw043   TGAAATCAGGCCAATAGAGAAAACCATGTTAAATAAGACTCGAACCTTTACAGCAGCTCCATTGGAAACTTTGCTTGGAG  7434
SCMV-Rw145   TGAAATCAGACCAATAGAGAAAACCATGTTAAATAAGACTCGAACCTTTACAGCAGCTCCATTGGAAACTTTGCTTGGAG  7436
             7360 7370 7380 7390 7400 7410 7420 7430
9640 bases   175 180 185 190 195 200
13 features  E I R P I E K T M L N K T R T F T A A P L E T L L G
             NIb
```

```
Consensus    GAAAAGTGTGCGTGGACGATTTTAATAATCAATTTTATTCACACCATTTAGAAGGCCCATGGACTGTTGGGATAACAAAA
SCMV-OH      GAAAAGTGTGCGTGGATGATTTTAATAATCAATTCTATTCACATCATTTAGAAGGTCCATGGACTGTTGGGATAACAAAA  7516
SCMV-See     GAAAGGTTTGCGTGGACGATTTTAATAATCAGTTTTATTCGCATCACCTTGAAGGCCCATGGACAGTTGGAATCACAAAA  7484
SCMV-Rw001   GAAAAGTGTGTGTGGACGATTTTAATAATCAATTTTATTCACACCATTTAGAAGGCCCGTGGACTGTTGGGATAACGAAA  7516
SCMV-Rw043   GAAAAGTGTGTGTGGACGATTTTAATAATCAATTTTATTCACACCATTTAGAAGGCCCGTGGACTGTTGGAATAACAAAA  7514
SCMV-Rw145   GAAAAGTGTGCGTGGACGATTTTAATAATCAATTTTACTCACACCATTTAGAAGGCCCATGGACTGTTGGGATAACAAAA  7516
             7440 7450 7460 7470 7480 7490 7500 7510
9640 bases   205 210 215 220 225
13 features  G K V C V D D F N N Q F Y S H H L E G P W T V G I T K
             NIb
```

```
Consensus    TTCTATGGAGGTTGGAATCGCTTACTTGAGAAGTTGCCAGAAGGATGGGTTTACTGCGACGCTGACGGNTCCCAATTCGA
SCMV-OH      TTCTATGGAGGTTGGAATCGCTTACTCGAGAAGTTACCAGAAGGATGGGTTTACTGCGATGCTGACGGGTCTCAATTTGA  7596
SCMV-See     TTTTATGGAGGATGGAATCGTCTACTCGAGAAGTTACCAGAAGGCTGGATTTATTGCGACGCCGATGGTTCACAATTTGA  7564
SCMV-Rw001   TTCTATGGAGGTTGGAATCGCTTACTTGAGAAGTTGCCGGAAGGATGGATTTACTGTGACGCTGACGGATCCCAGTTCGA  7596
SCMV-Rw043   TTCTATGGAGGTTGGAATCGCTTACTTGAGAAGTTGCCAGAAGGATGGGTTTACTGCGATGCTGACGGGTCCCAATTCGA  7594
SCMV-Rw145   TTTTATGGAGGTTGGAATCGCTTACTTGAGAAGTTGCCAGAAGGATGGGTTTATTGCGACGCTGACGGATCCCAATTCGA  7596
             7520 7530 7540 7550 7560 7570 7580 7590
9640 bases   230 235 240 245 250
13 features  F Y G G W N R L L E K L P E G W V Y C D A D G S Q F D
             NIb
```

```
Consensus    TAGTTCATTAACACCATATCTCATTAATGCAGTGTTGAATATTCGATTACAGTTCATGGAAGATTGGAACATAGGAGCAC
SCMV-OH      TAGTTCATTAACACCATATCTCATCAATGCAGTATTAAATATTCGATTGCAATTTATGGAAAATTGGGATATAGGAGCAC  7676
SCMV-See     TAGCTCATTAACGCCATACCTTATCAACGCCGTTTTACATATTCGGCTGCAATTTATGGAAGAATGGGCATTAGGGGCAC  7644
SCMV-Rw001   CAGTTCATTAACACCATATCTTATTAATGCAGTGTTGAATATTCGATTACAGTTCATGGAAGATTGGAACATAGGAGCAC  7676
SCMV-Rw043   TAGTTCATTAACACCATATCTCATTAATGCAGTGTTGAATATTCGATTACAGTTCATGGAAGATTGGAACATAGGAGCGC  7674
SCMV-Rw145   TAGTTCATTAACACCATATCTCATTAATGCAGTGTTGAATATTCGATTACAGTTCATGGAAGATTGGAACATAGGAGCGC  7676
             7600 7610 7620 7630 7640 7650 7660 7670
9640 bases   255 260 265 270 275 280
13 features  S S L T P Y L I N A V L N I R L Q F M E D W N I G A
             NIb
```

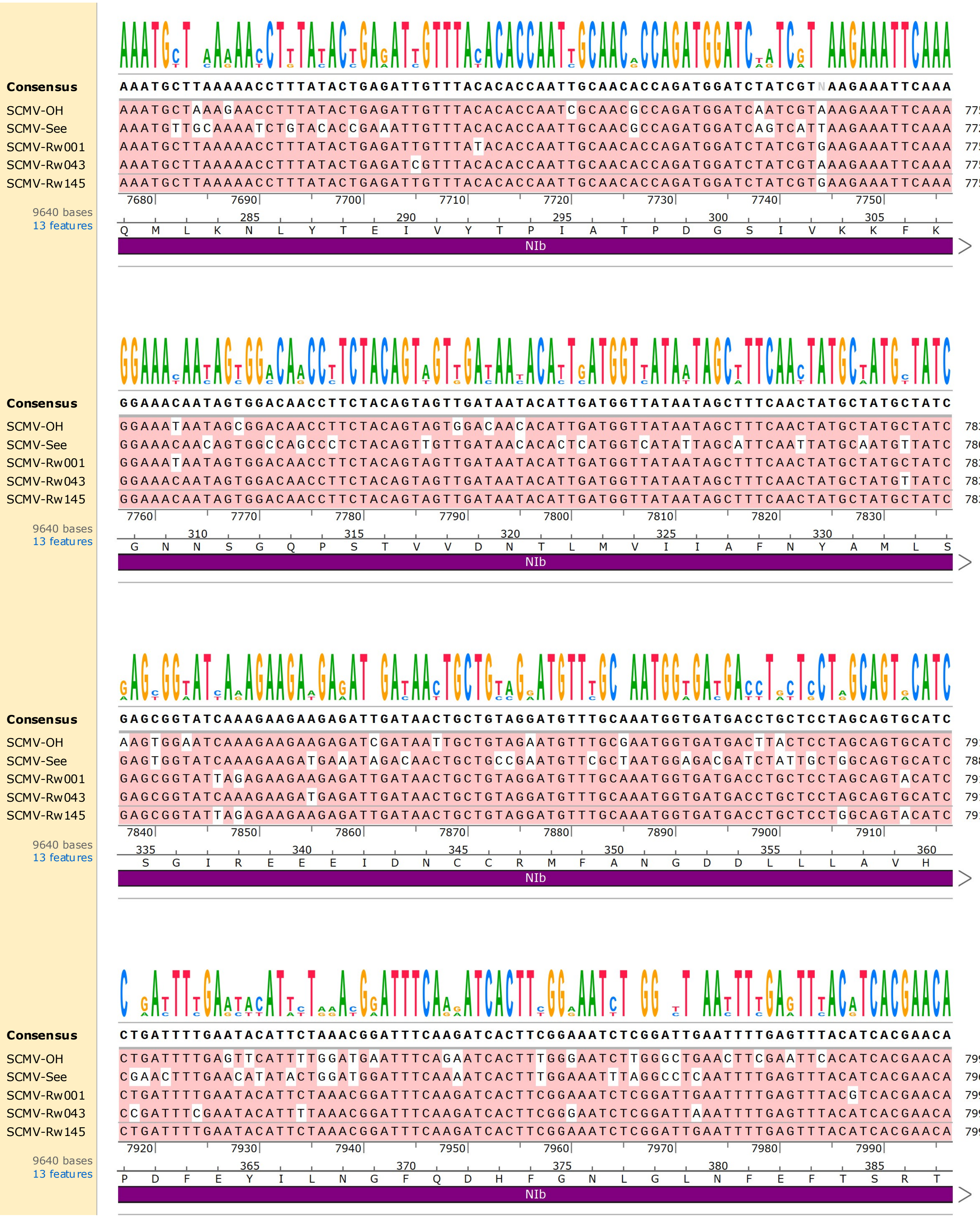

Consensus AAATGCTTAAAAACCTTTATACTGAGATTGTTTACACCAATTGCAACACCAGATGGATCTATCGTNAAGAAATTCAAA
SCMV-OH AAATGCTAAAGAACCTTTATACTGAGATTGTTTACACCAATCGCAACGCCAGATGGATCAATCGTAAAGAAATTCAAA 7756
SCMV-See AAATGTTGCAAAATCTGTACACCGAAATTGTTTACACCAATTGCAACGCCAGATGGATCAGTCATTAAGAAATTCAAA 7724
SCMV-Rw001 AAATGCTTAAAAACCTTTATACTGAGATTGTTTATACACCAATTGCAACACCAGATGGATCTATCGTGAAGAAATTCAAA 7756
SCMV-Rw043 AAATGCTTAAAAACCTTTATACTGAGATCGTTTACACCAATTGCAACACCAGATGGATCTATCGTAAAGAAATTCAAA 7754
SCMV-Rw145 AAATGCTTAAAAACCTTTATACTGAGATTGTTTACACCAATTGCAACACCAGATGGATCTATCGTGAAGAAATTCAAA 7756
7680 7690 7700 7710 7720 7730 7740 7750
9640 bases
13 features
285 290 295 300 305
Q M L K N L Y T E I V Y T P I A T P D G S I V K K F K
NIb
Consensus GGAAACAATAGTGGACAACCTTCTACAGTAGTTGATAATACATTGATGGTTATAATAGCTTTCAACTATGCTATGCTATC
SCMV-OH GGAAATAATAGCGGACAACCTTCTACAGTAGTGGACAACACATTGATGGTTATAATAGCTTTCAACTATGCTATGCTATC 7836
SCMV-See GGAAACAACAGTGGCCAGCCCTCTACAGTTGTTGATAACACACTCATGGTCATATTAGCATTCAATTATGCAATGTTATC 7804
SCMV-Rw001 GGAAATAATAGTGGACAACCTTCTACAGTAGTTGATAATACATTGATGGTTATAATAGCTTTCAACTATGCTATGCTATC 7836
SCMV-Rw043 GGAAACAATAGTGGACAACCTTCTACAGTAGTTGATAATACATTGATGGTTATAATAGCTTTCAACTATGCTATGTTATC 7834
SCMV-Rw145 GGAAACAATAGTGGACAACCTTCTACAGTAGTTGATAATACATTGATGGTTATAATAGCTTTCAACTATGCTATGCTATC 7836
7760 7770 7780 7790 7800 7810 7820 7830
9640 bases
13 features
310 315 320 325 330
G N N S G Q P S T V V D N T L M V I I A F N Y A M L S
NIb
Consensus GAGCGGTATCAAAGAAGAAGAGATTGATAACTGCTGTAGGATGTTTGCAAATGGTGATGACCTGCTCCTAGCAGTGCATC
SCMV-OH AAGTGGAATCAAAGAAGAAGAGATCGATAATTGCTGTAGAATGTTTGCGAATGGTGATGACTTACTCCTAGCAGTGCATC 7916
SCMV-See GAGTGGTATCAAAGAAGATGAAATAGACAACTGCTGCCGAATGTTCGCTAATGGAGACGATCTATTGCTGGCAGTGCATC 7884
SCMV-Rw001 GAGCGGTATTAGAGAAGAAGAGATTGATAACTGCTGTAGGATGTTTGCAAATGGTGATGACCTGCTCCTAGCAGTACATC 7916
SCMV-Rw043 GAGCGGTATCAAAGAAGATGAGATTGATAACTGCTGTAGGATGTTTGCAAATGGTGATGACCTGCTCCTAGCAGTGCATC 7914
SCMV-Rw145 GAGCGGTATTAGAGAAGAAGAGATTGATAACTGCTGTAGGATGTTTGCAAATGGTGATGACCTGCTCCTGGCAGTACATC 7916
7840 7850 7860 7870 7880 7890 7900 7910
9640 bases
13 features
335 340 345 350 355 360
S G I R E E E I D N C C R M F A N G D D L L L A V H
NIb
Consensus CTGATTTTGAATACATTCTAAACGGATTTCAAGATCACTTCGGAAATCTCGGATTGAATTTTGAGTTTACATCACGAACA
SCMV-OH CTGATTTTGAGTTCATTTTGGATGAATTTCAGAATCACTTTGGGAATCTTGGGCTGAACTTCGAATTCACATCACGAACA 7996
SCMV-See CGAACTTTGAACATATACTGGATGGATTTCAAAATCACTTTGGAAATTTAGGCCTCAATTTTGAGTTTACATCACGAACA 7964
SCMV-Rw001 CTGATTTTGAATACATTCTAAACGGATTTCAAGATCACTTCGGAAATCTCGGATTGAATTTTGAGTTTACGTCACGAACA 7996
SCMV-Rw043 CCGATTTCGAATACATTTTAAACGGATTTCAAGATCACTTCGGGAATCTCGGATTAAATTTTGAGTTTACATCACGAACA 7994
SCMV-Rw145 CTGATTTTGAATACATTCTAAACGGATTTCAAGATCACTTCGGAAATCTCGGATTGAATTTTGAGTTTACATCACGAACA 7996
7920 7930 7940 7950 7960 7970 7980 7990
9640 bases
13 features
365 370 375 380 385
P D F E Y I L N G F Q D H F G N L G L N F E F T S R T
NIb

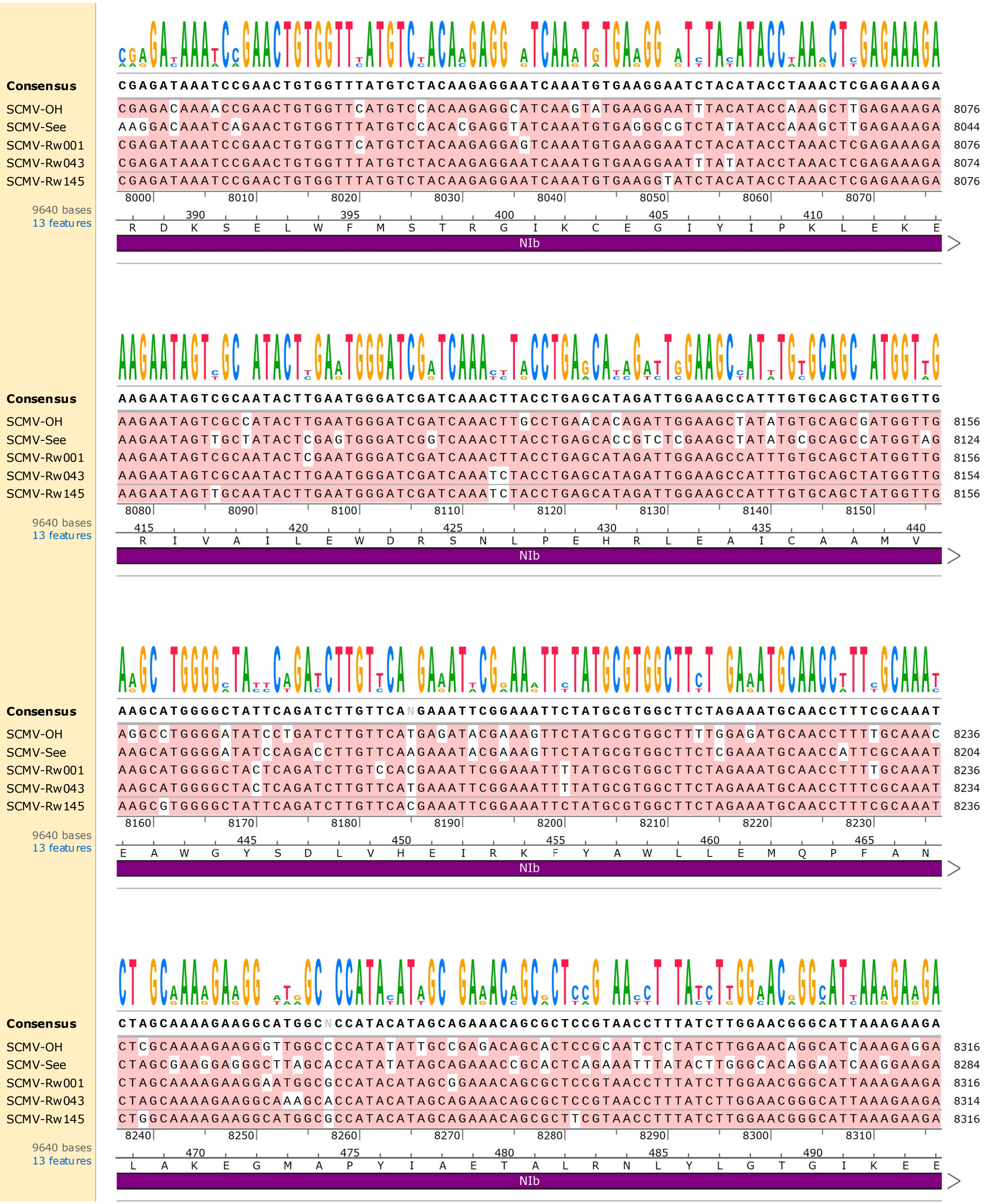
Consensus
CGAGATAAATCCGAACTGTGGTTTATGTCTACAAGAGGAATCAAATGTGAAGGAATCTACATACCTAAACTCGAGAAAGA
SCMV-OH CGAGACAAAACCGAACTGTGGTTCATGTCCACAAGAGGCATCAAGTATGAAGGAATTTACATACCAAAGCTTGAGAAAGA 8076
SCMV-See AAGGACAAATCAGAACTGTGGTTTATGTCCACACGAGGTATCAAATGTGAGGGCGTCTATATACCAAAGCTTGAGAAAGA 8044
SCMV-Rw001 CGAGATAAATCCGAACTGTGGTTCATGTCTACAAGAGGAGTCAAATGTGAAGGAATCTACATACCTAAACTCGAGAAAGA 8076
SCMV-Rw043 CGAGATAAATCCGAACTGTGGTTTATGTCTACAAGAGGAATCAAATGTGAAGGAATTTATATACCTAAACTCGAGAAAGA 8074
SCMV-Rw145 CGAGATAAATCCGAACTGTGGTTTATGTCTACAAGAGGAATCAAATGTGAAGGTATCTACATACCTAAACTCGAGAAAGA 8076
8000 8010 8020 8030 8040 8050 8060 8070
9640 bases
13 features
390 395 400 405 410
R D K S E L W F M S T R G I K C E G I Y I P K L E K E
NIb
Consensus
AAGAATAGTCGCAATACTTGAATGGGATCGATCAAACTTACCTGAGCATAGATTGGAAGCCATTTGTGCAGCTATGGTTG
SCMV-OH AAGAATAGTCGCCATACTTGAATGGGATCGATCAAACTTGCCTGAACACAGATTGGAAGCTATATGTGCAGCGATGGTTG 8156
SCMV-See AAGAATAGTTGCTATACTCGAGTGGGATCGGTCAAACTTACCTGAGCACCGTCTCGAAGCTATATGCGCAGCCATGGTAG 8124
SCMV-Rw001 AAGAATAGTCGCAATACTCGAATGGGATCGATCAAACTTACCTGAGCATAGATTGGAAGCCATTTGTGCAGCTATGGTTG 8156
SCMV-Rw043 AAGAATAGTCGCAATACTTGAATGGGATCGATCAAATCTACCTGAGCATAGATTGGAAGCCATTTGTGCAGCTATGGTTG 8154
SCMV-Rw145 AAGAATAGTTGCAATACTTGAATGGGATCGATCAAATCTACCTGAGCATAGATTGGAAGCCATTTGTGCAGCTATGGTTG 8156
8080 8090 8100 8110 8120 8130 8140 8150
9640 bases
13 features
415 420 425 430 435 440
R I V A I L E W D R S N L P E H R L E A I C A A M V
NIb
Consensus
AAGCATGGGGCTATTCAGATCTTGTTCANGAAATTCGGAAATTCTATGCGTGGCTTCTAGAAATGCAACCTTTCGCAAAT
SCMV-OH AGGCCTGGGGATATCCTGATCTTGTTCATGAGATACGAAAGTTCTATGCGTGGCTTTTGGAGATGCAACCTTTTGCAAAC 8236
SCMV-See AAGCATGGGGATATCCAGACCTTGTTCAAGAAATACGAAAGTTCTATGCGTGGCTTCTCGAAATGCAACCATTCGCAAAT 8204
SCMV-Rw001 AAGCATGGGGCTACTCAGATCTTGTCCACGAAATTCGGAAATTTTATGCGTGGCTTCTAGAAATGCAACCTTTTGCAAAT 8236
SCMV-Rw043 AAGCATGGGGCTACTCAGATCTTGTTCATGAAATTCGGAAATTTTATGCGTGGCTTCTAGAAATGCAACCTTTCGCAAAT 8234
SCMV-Rw145 AAGCGTGGGGCTATTCAGATCTTGTTCACGAAATTCGGAAATTCTATGCGTGGCTTCTAGAAATGCAACCTTTCGCAAAT 8236
8160 8170 8180 8190 8200 8210 8220 8230
9640 bases
13 features
445 450 455 460 465
E A W G Y S D L V H E I R K F Y A W L L E M Q P F A N
NIb
Consensus
CTAGCAAAAGAAGGCATGGCNCCATACATAGCAGAAACAGCGCTCCGTAACCTTTATCTTGGAACGGGCATTAAAGAAGA
SCMV-OH CTCGCAAAAGAAGGGTTGGCCCCATATATTGCCGAGACAGCACTCCGCAATCTCTATCTTGGAACAGGCATCAAAGAGGA 8316
SCMV-See CTAGCGAAGGAGGGCTTAGCACCATATATAGCAGAAACCGCACTCAGAAATTTATACTTGGGCACAGGAATCAAGGAAGA 8284
SCMV-Rw001 CTAGCAAAAGAAGGAATGGCGCCATACATAGCGGAAACAGCGCTCCGTAACCTTTATCTTGGAACGGGCATTAAAGAAGA 8316
SCMV-Rw043 CTAGCAAAAGAAGGCAAAGCACCATACATAGCAGAAACAGCGCTCCGTAACCTTTATCTTGGAACGGGCATTAAAGAAGA 8314
SCMV-Rw145 CTGGCAAAAGAAGGCATGGCGCCATACATAGCAGAAACAGCGCTTCGTAACCTTTATCTTGGAACGGGCATTAAAGAAGA 8316
8240 8250 8260 8270 8280 8290 8300 8310
9640 bases
13 features
470 475 480 485 490
L A K E G M A P Y I A E T A L R N L Y L G T G I K E E
NIb

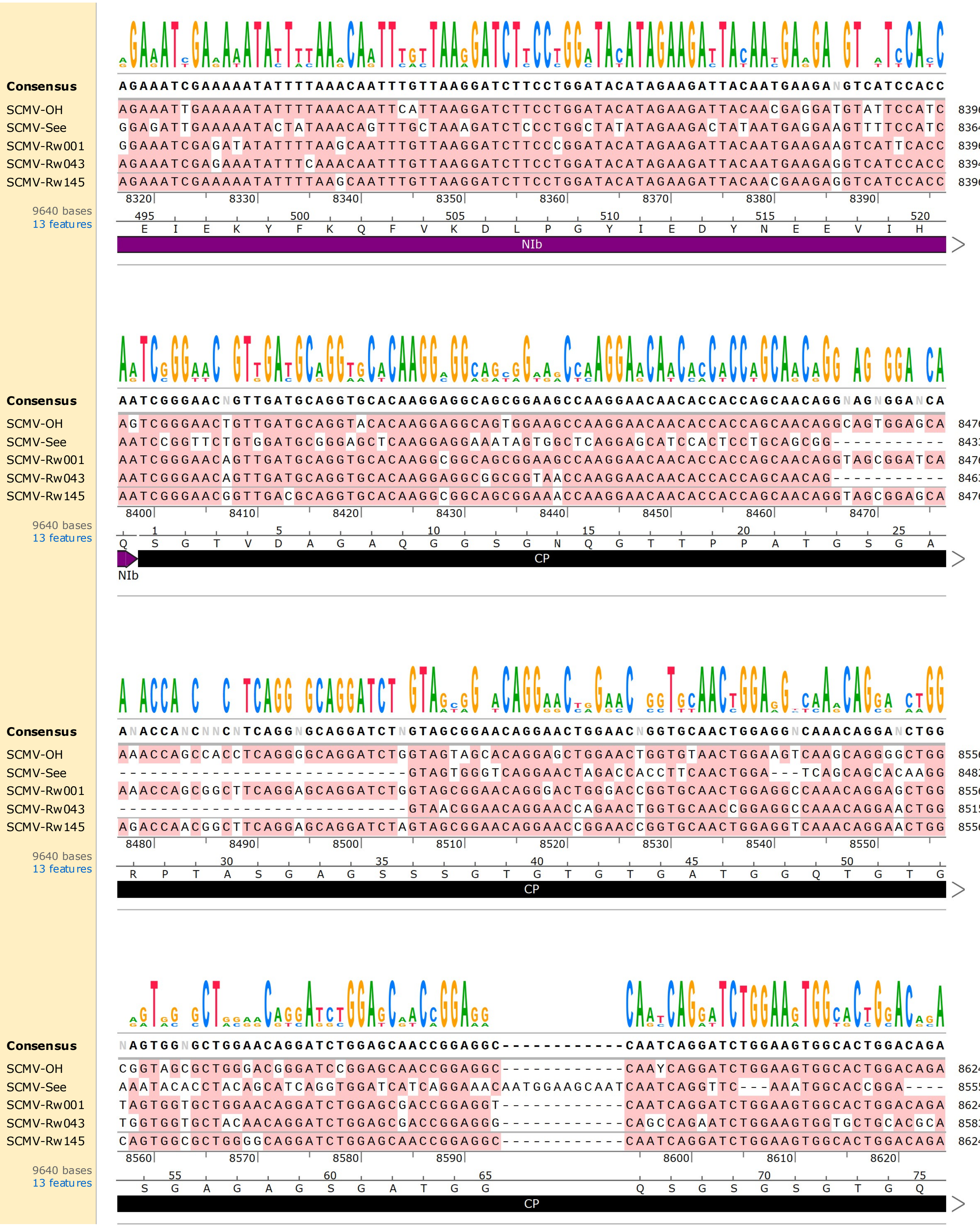
Consensus
AGAAATCGAAAAATATTTTAAACAATTTGTTAAGGATCTTCCTGGATACATAGAAGATTACAATGAAGANGTCATCCACC
SCMV-OH AGAAATTGAAAAATATTTTAAACAATTCATTAAGGATCTTCCTGGATACATAGAAGATTACAACGAGGATGTATTCCATC 8396
SCMV-See GGAGATTGAAAAATACTATAAACAGTTTGCTAAAGATCTCCCTGGCTATATAGAAGACTATAATGAGGAAGTTTTCCATC 8364
SCMV-Rw001 GGAAATCGAGATATATTTTAAGCAATTTGTTAAGGATCTTCCCGGATACATAGAAGATTACAATGAAGAAGTCATTCACC 8396
SCMV-Rw043 AGAAATCGAGAAATATTTCAAACAATTTGTTAAGGATCTTCCTGGATACATAGAAGATTACAATGAAGAGGTCATCCACC 8394
SCMV-Rw145 AGAAATCGAAAAATATTTTAAGCAATTTGTTAAGGATCTTCCTGGATACATAGAAGATTACAACGAAGAGGTCATCCACC 8396
8320 8330 8340 8350 8360 8370 8380 8390
9640 bases
13 features
495 500 505 510 515 520
E I E K Y F K Q F V K D L P G Y I E D Y N E E V I H
NIb
Consensus
AATCGGGAACNGTTGATGCAGGTGCACAAGGAGGCAGCGGAAGCCAAGGAACAACACCACCAGCAACAGGNAGNGGANCA
SCMV-OH AGTCGGGAACTGTTGATGCAGGTACACAAGGAGGCAGTGGAAGCCAAGGAACAACACCACCAGCAACAGGCAGTGGAGCA 8476
SCMV-See AATCCGGTTCTGTGGATGCGGGAGCTCAAGGAGGAAATAGTGGCTCAGGAGCATCCACTCCTGCAGCGG----------- 8433
SCMV-Rw001 AATCGGGAACAGTTGATGCAGGTGCACAAGGCGGCAGCGGAAGCCAAGGAACAACACCACCAGCAACAGGTAGCGGATCA 8476
SCMV-Rw043 AATCGGGAACAGTTGATGCAGGTGCACAAGGAGGCGGCGGTAACCAAGGAACAACACCACCAGCAACAG----------- 8463
SCMV-Rw145 AATCGGGAACGGTTGACGCAGGTGCACAAGGCGGCAGCGGAAACCAAGGAACAACACCACCAGCAACAGGTAGCGGAGCA 8476
8400 8410 8420 8430 8440 8450 8460 8470
9640 bases
13 features
1 5 10 15 20 25
Q S G T V D A G A Q G G S G N Q G T T P P A T G S G A
CP
NIb
Consensus
ANACCANCNNCNTCAGGNGCAGGATCTNGTAGCGGAACAGGAACTGGAACNGGTGCAACTGGAGGNCAAACAGGANCTGG
SCMV-OH AAACCAGCCACCTCAGGGGCAGGATCTGGTAGTAGCACAGGAGCTGGAACTGGTGTAACTGGAAGTCAAGCAGGGGCTGG 8556
SCMV-See ----------------------------GTAGTGGGTCAGGAACTAGACCACCTTCAACTGGA---TCAGCAGCACAAGG 8482
SCMV-Rw001 AAACCAGCGGCTTCAGGAGCAGGATCTGGTAGCGGAACAGGGACTGGGACCGGTGCAACTGGAGGCCAAACAGGAGCTGG 8556
SCMV-Rw043 ----------------------------GTAACGGAACAGGAACCAGAACTGGTGCAACCGGAGGCCAAACAGGAACTGG 8515
SCMV-Rw145 AGACCAACGGCTTCAGGAGCAGGATCTAGTAGCGGAACAGGAACCGGAACCGGTGCAACTGGAGGTCAAACAGGAACTGG 8556
8480 8490 8500 8510 8520 8530 8540 8550
9640 bases
13 features
30 35 40 45 50
R P T A S G A G S S S G T G T G T G A T G G Q T G T G
CP
Consensus
NAGTGGNGCTGGAACAGGATCTGGAGCAACCGGAGGC------------CAATCAGGATCTGGAAGTGGCACTGGACAGA
SCMV-OH CGGTAGCGCTGGACGGGATCCGGAGCAACCGGAGGC------------CAAYCAGGATCTGGAAGTGGCACTGGACAGA 8624
SCMV-See AAATACACCTACAGCATCAGGTGGATCATCAGGAAACAATGGAAGCAATCAATCAGGTTC---AAATGGCACCGGA---- 8555
SCMV-Rw001 TAGTGGTGCTGGAACAGGATCTGGAGCGACCGGAGGT------------CAATCAGGATCTGGAAGTGGCACTGGACAGA 8624
SCMV-Rw043 TGGTGGTGCTACAACAGGATCTGGAGCGACCGGAGGG------------CAGCCAGAATCTGGAAGTGGTGCTGCACGCA 8583
SCMV-Rw145 CAGTGGCGCTGGGGCAGGATCTGGAGCAACCGGAGGC------------CAATCAGGATCTGGAAGTGGCACTGGACAGA 8624
8560 8570 8580 8590 8600 8610 8620
9640 bases
13 features
55 60 65 70 75
S G A G A G S G A T G G Q S G S G S G T G Q
CP

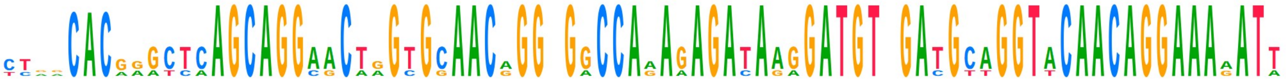

| | | |
|---|---|---|
| **Consensus** | CTNNCACGGGCTCAGCAGGAACTGGTGCAACAGGNGGCCAAAGAGATAAGGATGTAGATGCAGGTACAACAGGAAAAATT | |
| SCMV-OH | TTAACACGGGTTCAGCAGGAACTAGTGCAACAGGAGGCCAAAGAGATAGGGATGTGGATGCAGGTACAACAGGAAAAATT | 8704 |
| SCMV-See | --------AACCAAGCAGGCGCAAGCGGAACAGGGGACCAAAGAGACAAAGATGTCGACGTTGGTTCAACAGGAAAGATA | 8627 |
| SCMV-Rw001 | CTGGCACAGGCTCAGCAGGAACTGGTGCAACGGGAGGCCAGAGAGATAAGGATGTAGATGCAGGTACAACAGGAAAAATT | 8704 |
| SCMV-Rw043 | CCAACACGAGCTCAGCAGGAACTGGTGCAACGGGTGGCCAGAAAGATAAGGATGTAGATGCAGGTACAACAGGAAAAATT | 8663 |
| SCMV-Rw145 | CTGGCACGGGCTCAGCAGGAACTGGTGCAACAGGGGGCCAAAGAGATAAGGATGTAGATGCAGGTACAACAGGAAAGATT | 8704 |

8630 8640 8650 8660 8670 8680 8690 8700

9640 bases
13 features

80 85 90 95 100
T G T G S A G T G A T G G Q R D K D V D A G T T G K I
CP

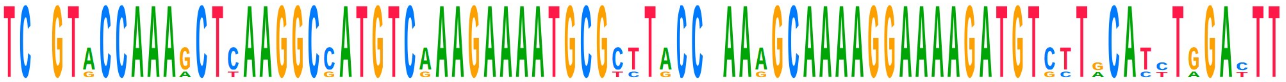

| | | |
|---|---|---|
| **Consensus** | TCTGTACCAAAGCTCAAGGCCATGTCAAAGAAAATGCGCTTACCGAAAGCAAAAGGAAAAGATGTCTTGCATCTGGACTT | |
| SCMV-OH | TCTGTACCAAAGCTCAAGGCCATGTCAAAGAAAATGCGCTTACCTAAAGCAAAAGGAAAAGATGTGCTACATTTGGATTT | 8784 |
| SCMV-See | TCAGTGCCAAAACTTAAGGCGATGTCGAAGAAAATGCGTCTGCCAAAGGCAAAAGGAAAAGATGTCTTACACTTAGATTT | 8707 |
| SCMV-Rw001 | TCTGTACCAAAGCTCAAGGCCATGTCAAAGAAAATGCGCTTACCGAAAGCAAAAGGAAAAGATGTCTTGCATCTGGACTT | 8784 |
| SCMV-Rw043 | TCTGTACCAAAGCTCAAGGCCATGTCAAAGAAAATGCGCTTACCGAAAGCAAAAGGAAAAGATGTCTTGCATCTGGACTT | 8743 |
| SCMV-Rw145 | TCCGTACCAAAGCTCAAGGCCATGTCAAAGAAAATGCGCTTACCGAAAGCAAAAGGAAAAGATGTCTTGCATCTGGACTT | 8784 |

8710 8720 8730 8740 8750 8760 8770 8780

9640 bases
13 features

105 110 115 120 125
S V P K L K A M S K K M R L P K A K G K D V L H L D F
CP

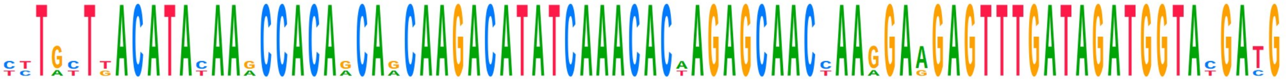

| | | |
|---|---|---|
| **Consensus** | CTTGCTTACATACAAGCCACAGCAGCAAGACATATCAAACACAAGAGCAACCAAGGAAGAGTTTGATAGATGGTACGATG | |
| SCMV-OH | TCTATTGACATATAAACCACAACAACAAGACATATCAAACACTAGAGCAACCAAAGAAGAGTTTGATAGATGGTATGATG | 8864 |
| SCMV-See | TCTGTTGACATATAAACCACAACAACAAGACATATCAAACACTAGAGCAACCAAGGAAGAGTTTGATAGATGGTATGACG | 8787 |
| SCMV-Rw001 | CTTGCTTACATACAAGCCACAGCAGCAAGACATATCAAACACAAGAGCAACTAAGGAAGAGTTTGATAGATGGTACGATG | 8864 |
| SCMV-Rw043 | CTTGCTTACATACAAGCCACAGCAGCAAGACATATCAAACACAAGAGCAACTAAGGAGGAGTTTGATAGATGGTACGATG | 8823 |
| SCMV-Rw145 | CTTGCTTACATACAAGCCACAGCAGCAAGACATATCAAACACAAGAGCAACCAAGGAAGAGTTTGATAGATGGTACGATG | 8864 |

8790 8800 8810 8820 8830 8840 8850 8860

9640 bases
13 features

130 135 140 145 150 155
L L T Y K P Q Q Q D I S N T R A T K E E F D R W Y D
CP

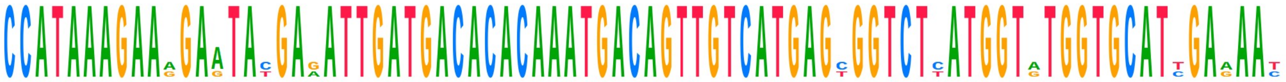

| | | |
|---|---|---|
| **Consensus** | CCATAAAGAAAGAATACGAGATTGATGACACACAAATGACAGTTGTCATGAGCGGTCTCATGGTATGGTGCATTGAAAAT | |
| SCMV-OH | CCATAAAGAAGGAATATGAAATTGATGACACACAAATGACAGTTGTCATGAGTGGTCTTATGGTATGGTGCATCGAAAAT | 8944 |
| SCMV-See | CCATAAAGAAGGAGTATGAAATTGATGACACACAAATGACAGTTGTCATGAGTGGTCTCATGGTATGGTGCATCGAAAAT | 8867 |
| SCMV-Rw001 | CCATAAAGAAAGAATACGAGATTGATGACACACAAATGACAGTTGTCATGAGCGGTCTCATGGTGTGGTGCATTGAGAAC | 8944 |
| SCMV-Rw043 | CCATAAAGAAAGAATACGAGATTGATGACACACAAATGACAGTTGTCATGAGCGGTCTTATGGTGTGGTGCATTGAAAAT | 8903 |
| SCMV-Rw145 | CCATAAAGAAAGAATACGAGATTGATGACACACAAATGACAGTTGTCATGAGCGGTCTCATGGTATGGTGCATTGAGAAC | 8944 |

8870 8880 8890 8900 8910 8920 8930 8940

9640 bases
13 features

160 165 170 175 180
A I K K E Y E I D D T Q M T V V M S G L M V W C I E N
CP

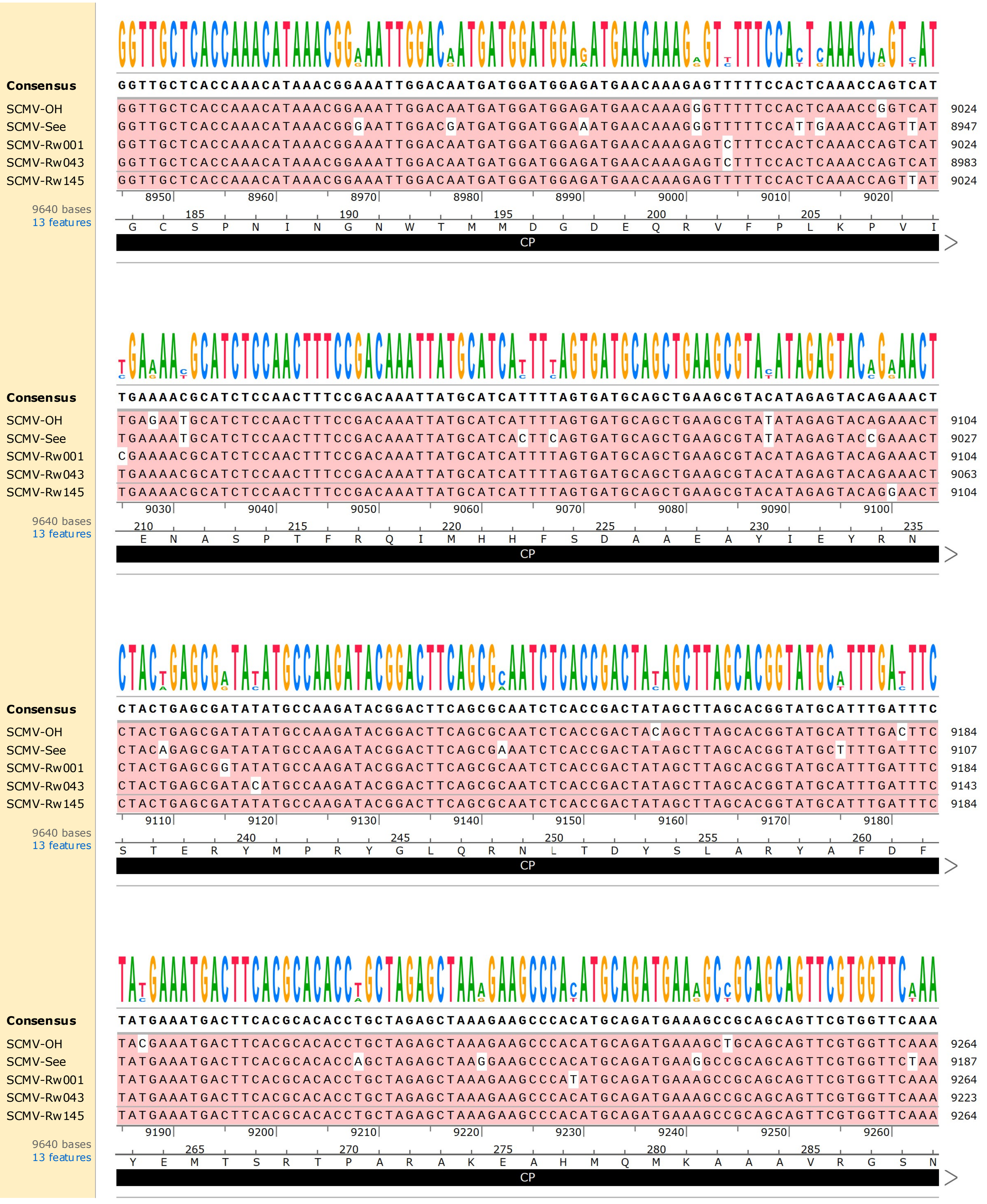
Consensus GGTTGCTCACCAAACATAAACGGAAATTGGACAATGATGGATGGAGATGAACAAAGAGTTTTTCCACTCAAACCAGTCAT
SCMV-OH GGTTGCTCACCAAACATAAACGGAAATTGGACAATGATGGATGGAGATGAACAAAGGGTTTTTCCACTCAAACCGGTCAT 9024
SCMV-See GGTTGCTCACCAAACATAAACGGGAATTGGACGATGATGGATGGAAATGAACAAAGGGTTTTTCCATTGAAACCAGTTAT 8947
SCMV-Rw001 GGTTGCTCACCAAACATAAACGGAAATTGGACAATGATGGATGGAGATGAACAAAGAGTCTTTCCACTCAAACCAGTCAT 9024
SCMV-Rw043 GGTTGCTCACCAAACATAAACGGAAATTGGACAATGATGGATGGAGATGAACAAAGAGTCTTTCCACTCAAACCAGTCAT 8983
SCMV-Rw145 GGTTGCTCACCAAACATAAACGGAAATTGGACAATGATGGATGGAGATGAACAAAGAGTTTTTCCACTCAAACCAGTTAT 9024
8950 8960 8970 8980 8990 9000 9010 9020
9640 bases
13 features
185 190 195 200 205
G C S P N I N G N W T M M D G D E Q R V F P L K P V I
CP
Consensus TGAAAACGCATCTCCAACTTTCCGACAAATTATGCATCATTTTAGTGATGCAGCTGAAGCGTACATAGAGTACAGAAACT
SCMV-OH TGAGAATGCATCTCCAACTTTCCGACAAATTATGCATCATTTTAGTGATGCAGCTGAAGCGTATATAGAGTACAGAAACT 9104
SCMV-See TGAAAATGCATCTCCAACTTTCCGACAAATTATGCATCACTTCAGTGATGCAGCTGAAGCGTATATAGAGTACCGAAACT 9027
SCMV-Rw001 CGAAAACGCATCTCCAACTTTCCGACAAATTATGCATCATTTTAGTGATGCAGCTGAAGCGTACATAGAGTACAGAAACT 9104
SCMV-Rw043 TGAAAACGCATCTCCAACTTTCCGACAAATTATGCATCATTTTAGTGATGCAGCTGAAGCGTACATAGAGTACAGAAACT 9063
SCMV-Rw145 TGAAAACGCATCTCCAACTTTCCGACAAATTATGCATCATTTTAGTGATGCAGCTGAAGCGTACATAGAGTACAGGAACT 9104
9030 9040 9050 9060 9070 9080 9090 9100
9640 bases
13 features
210 215 220 225 230 235
E N A S P T F R Q I M H H F S D A A E A Y I E Y R N
CP
Consensus CTACTGAGCGATATATGCCAAGATACGGACTTCAGCGCAATCTCACCGACTATAGCTTAGCACGGTATGCATTTGATTTC
SCMV-OH CTACTGAGCGATATATGCCAAGATACGGACTTCAGCGCAATCTCACCGACTACAGCTTAGCACGGTATGCATTTGACTTC 9184
SCMV-See CTACAGAGCGATATATGCCAAGATACGGACTTCAGCGAAATCTCACCGACTATAGCTTAGCACGGTATGCTTTTGATTTC 9107
SCMV-Rw001 CTACTGAGCGGTATATGCCAAGATACGGACTTCAGCGCAATCTCACCGACTATAGCTTAGCACGGTATGCATTTGATTTC 9184
SCMV-Rw043 CTACTGAGCGATACATGCCAAGATACGGACTTCAGCGCAATCTCACCGACTATAGCTTAGCACGGTATGCATTTGATTTC 9143
SCMV-Rw145 CTACTGAGCGATATATGCCAAGATACGGACTTCAGCGCAATCTCACCGACTATAGCTTAGCACGGTATGCATTTGATTTC 9184
9110 9120 9130 9140 9150 9160 9170 9180
9640 bases
13 features
240 245 250 255 260
S T E R Y M P R Y G L Q R N L T D Y S L A R Y A F D F
CP
Consensus TATGAAATGACTTCACGCACACCTGCTAGAGCTAAAGAAGCCCACATGCAGATGAAAGCCGCAGCAGTTCGTGGTTCAAA
SCMV-OH TACGAAATGACTTCACGCACACCTGCTAGAGCTAAAGAAGCCCACATGCAGATGAAAGCTGCAGCAGTTCGTGGTTCAAA 9264
SCMV-See TATGAAATGACTTCACGCACACCAGCTAGAGCTAAGGAAGCCCACATGCAGATGAAGGCCGCAGCAGTTCGTGGTTCTAA 9187
SCMV-Rw001 TATGAAATGACTTCACGCACACCTGCTAGAGCTAAAGAAGCCCATATGCAGATGAAAGCCGCAGCAGTTCGTGGTTCAAA 9264
SCMV-Rw043 TATGAAATGACTTCACGCACACCTGCTAGAGCTAAAGAAGCCCACATGCAGATGAAAGCCGCAGCAGTTCGTGGTTCAAA 9223
SCMV-Rw145 TATGAAATGACTTCACGCACACCTGCTAGAGCTAAAGAAGCCCACATGCAGATGAAAGCCGCAGCAGTTCGTGGTTCAAA 9264
9190 9200 9210 9220 9230 9240 9250 9260
9640 bases
13 features
265 270 275 280 285
Y E M T S R T P A R A K E A H M Q M K A A A V R G S N
CP

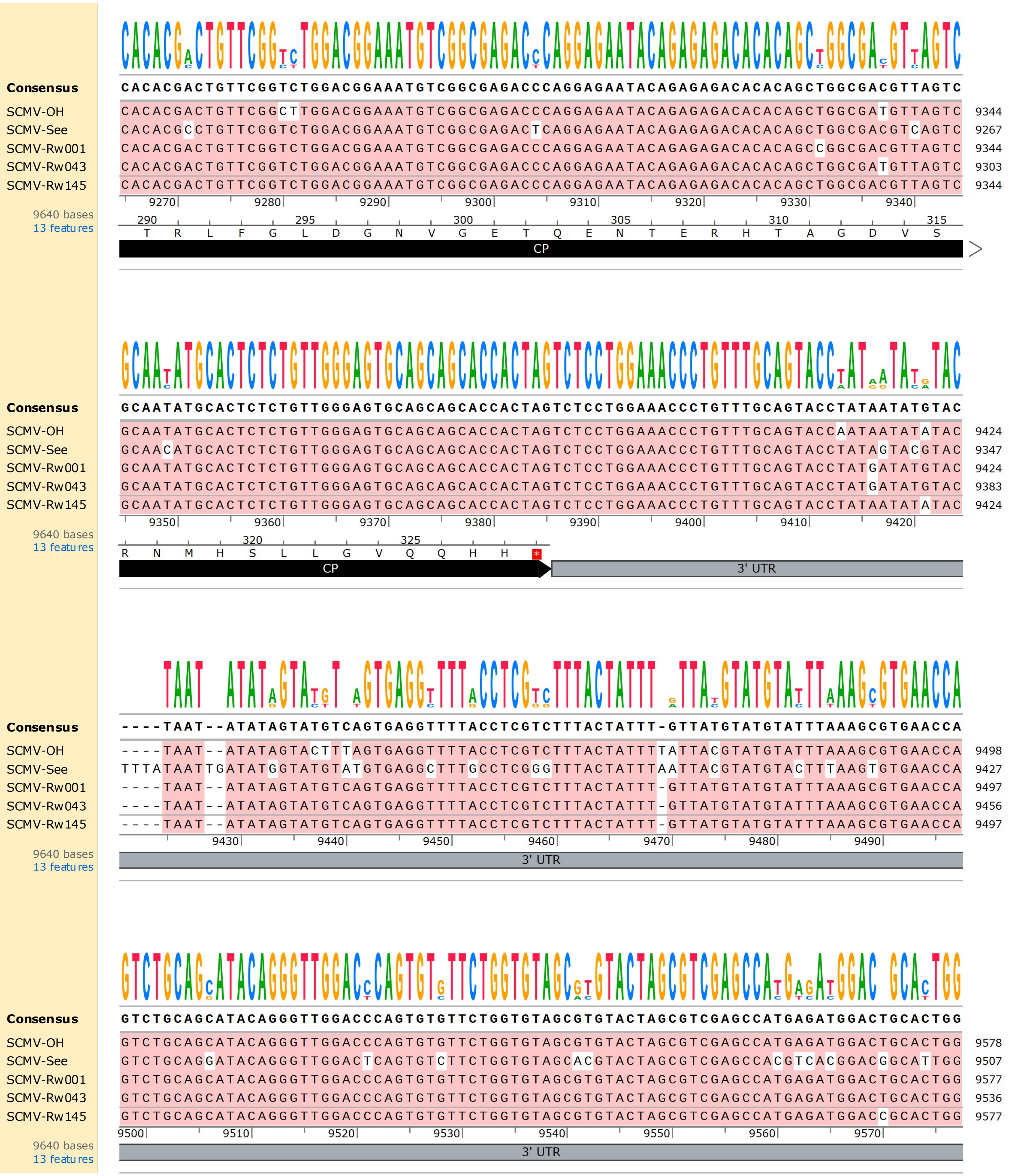
Consensus
CACACGACTGTTCGGTCTGGACGGAAATGTCGGCGAGACCCAGGAGAATACAGAGAGACACACAGCTGGCGACGTTAGTC
SCMV-OH CACACGACTGTTCGGCTTGGACGGAAATGTCGGCGAGACCCAGGAGAATACAGAGAGACACACAGCTGGCGATGTTAGTC 9344
SCMV-See CACACGCCTGTTCGGTCTGGACGGAAATGTCGGCGAGACTCAGGAGAATACAGAGAGACACACAGCTGGCGACGTCAGTC 9267
SCMV-Rw001 CACACGACTGTTCGGTCTGGACGGAAATGTCGGCGAGACCCAGGAGAATACAGAGAGACACACAGCCGGCGACGTTAGTC 9344
SCMV-Rw043 CACACGACTGTTCGGTCTGGACGGAAATGTCGGCGAGACCCAGGAGAATACAGAGAGACACACAGCTGGCGATGTTAGTC 9303
SCMV-Rw145 CACACGACTGTTCGGTCTGGACGGAAATGTCGGCGAGACCCAGGAGAATACAGAGAGACACACAGCTGGCGACGTTAGTC 9344
9270 9280 9290 9300 9310 9320 9330 9340
9640 bases
13 features
290 295 300 305 310 315
T R L F G L D G N V G E T Q E N T E R H T A G D V S
CP
Consensus
GCAATATGCACTCTCTGTTGGGAGTGCAGCAGCACCACTAGTCTCCTGGAAACCCTGTTTGCAGTACCTATAATATGTAC
SCMV-OH GCAATATGCACTCTCTGTTGGGAGTGCAGCAGCACCACTAGTCTCCTGGAAACCCTGTTTGCAGTACCAATAATATATAC 9424
SCMV-See GCAACATGCACTCTCTGTTGGGAGTGCAGCAGCACCACTAGTCTCCTGGAAACCCTGTTTGCAGTACCTATAGTACGTAC 9347
SCMV-Rw001 GCAATATGCACTCTCTGTTGGGAGTGCAGCAGCACCACTAGTCTCCTGGAAACCCTGTTTGCAGTACCTATGATATGTAC 9424
SCMV-Rw043 GCAATATGCACTCTCTGTTGGGAGTGCAGCAGCACCACTAGTCTCCTGGAAACCCTGTTTGCAGTACCTATGATATGTAC 9383
SCMV-Rw145 GCAATATGCACTCTCTGTTGGGAGTGCAGCAGCACCACTAGTCTCCTGGAAACCCTGTTTGCAGTACCTATAATATATAC 9424
9350 9360 9370 9380 9390 9400 9410 9420
9640 bases
13 features
320 325
R N M H S L L G V Q Q H H *
CP
3' UTR
Consensus
----TAAT--ATATAGTATGTCAGTGAGGTTTTACCTCGTCTTTACTATTT-GTTATGTATGTATTTAAAGCGTGAACCA
SCMV-OH ----TAAT--ATATAGTACTTTAGTGAGGTTTTACCTCGTCTTTACTATTTTATTACGTATGTATTTAAAGCGTGAACCA 9498
SCMV-See TTTATAATTGATATGGTATGTATGTGAGGCTTTGCCTCGGGTTTACTATTTAATTACGTATGTACTTTAAGTGTGAACCA 9427
SCMV-Rw001 ----TAAT--ATATAGTATGTCAGTGAGGTTTTACCTCGTCTTTACTATTT-GTTATGTATGTATTTAAAGCGTGAACCA 9497
SCMV-Rw043 ----TAAT--ATATAGTATGTCAGTGAGGTTTTACCTCGTCTTTACTATTT-GTTATGTATGTATTTAAAGCGTGAACCA 9456
SCMV-Rw145 ----TAAT--ATATAGTATGTCAGTGAGGTTTTACCTCGTCTTTACTATTT-GTTATGTATGTATTTAAAGCGTGAACCA 9497
9430 9440 9450 9460 9470 9480 9490
9640 bases
13 features
3' UTR
Consensus
GTCTGCAGCATACAGGGTTGGACCCAGTGTGTTCTGGTGTAGCGTGTACTAGCGTCGAGCCATGAGATGGACTGCACTGG
SCMV-OH GTCTGCAGCATACAGGGTTGGACCCAGTGTGTTCTGGTGTAGCGTGTACTAGCGTCGAGCCATGAGATGGACTGCACTGG 9578
SCMV-See GTCTGCAGGATACAGGGTTGGACTCAGTGTCTTCTGGTGTAGCACGTACTAGCGTCGAGCCACGTCACGGACGGCATTGG 9507
SCMV-Rw001 GTCTGCAGCATACAGGGTTGGACCCAGTGTGTTCTGGTGTAGCGTGTACTAGCGTCGAGCCATGAGATGGACTGCACTGG 9577
SCMV-Rw043 GTCTGCAGCATACAGGGTTGGACCCAGTGTGTTCTGGTGTAGCGTGTACTAGCGTCGAGCCATGAGATGGACTGCACTGG 9536
SCMV-Rw145 GTCTGCAGCATACAGGGTTGGACCCAGTGTGTTCTGGTGTAGCGTGTACTAGCGTCGAGCCATGAGATGGACCGCACTGG 9577
9500 9510 9520 9530 9540 9550 9560 9570
9640 bases
13 features
3' UTR

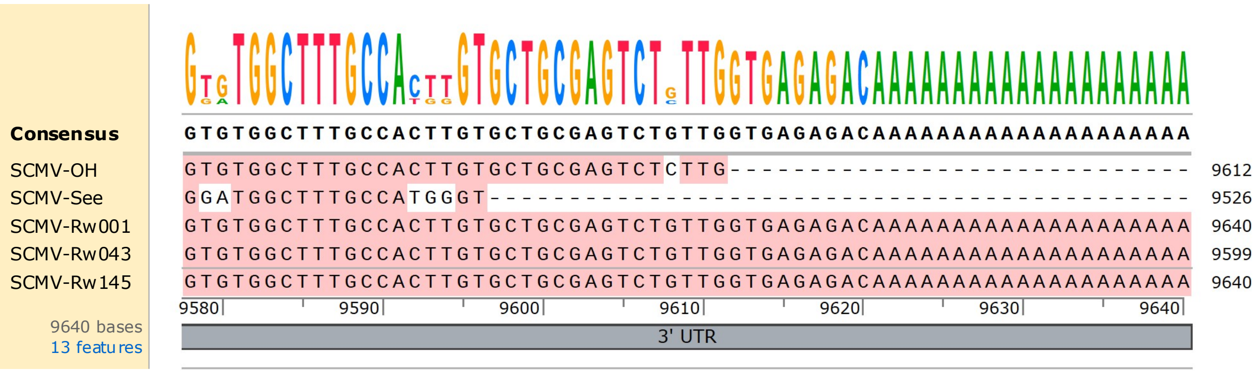


**Supplementary Figure S2.** Nucleotide alignment of the near-full genome sequences of the five SCMV isolates. Bases that match the consensus sequence (bold) are shaded pink. Above the consensus sequence is a sequence logo indicating the most prevalent base at each position. Untranslated regions (UTR, grey) and open reading frames of viral genes are indicated below the alignment: P1 (bright pink), helper component protease (HC-Pro, red), P3 (orange), Pretty Interesting Potyviridae ORF (PIPO, teal), 6KI (yellow), cylindrical inclusion protein (CIP, green), 6K2 (light blue), viral protein genome-linked (VPg, dark blue), nuclear inclusion a protease (NIa-Pro, light purple), nuclear inclusion b (NIb, dark purple), and coat protein (CP, black). Amino acids are indicated above each position in the polyprotein based on translation of the SCMV-Rw145 sequence. Alignment was performed in SnapGene v 7.0.3 using MUSCLE and default parameters.
